# Allosteric Disordering of eIF2B Regulates the Integrated Stress Response

**DOI:** 10.1101/2025.09.11.675390

**Authors:** Udit Dalwadi, Advait Subramanian, Aniliese Deal, Julia E. Conrad, Meera Venkatesh, Morgane Boone, Pascal F. Egea, Lingjie He, Nimit Jain, D. John Lee, Yuwei Liu, Lucas C. Reineke, Kazuki Saito, Nathaniel Talledge, Hannah Toutkoushian, Maxence Le Vasseur, Francesca Zappa, Raoul J. de Groot, Diego Acosta-Alvear, Christopher P. Arthur, Jodi Nunnari, Mauro Costa-Mattioli, James J. Crawford, Frank J. M. van Kuppeveld, Tristan I. Croll, Peter Walter, Adam Frost

**Author notes:** Corresponding authors (AF), (PW). These authors contributed equally to this work (Udit Dalwadi, Advait Subramanian).

## Abstract

The ternary complex (TC), composed of translation initiation factor eIF2, GTP, and initiator methionyl tRNA, delivers the first amino acid to the ribosome to initiate protein synthesis. The activity of the decameric eukaryotic initiation factor 2B complex (eIF2B) initiates TC assembly by catalyzing GDP to GTP exchange on eIF2, thereby setting the TC levels in the cell. Stress-induced phosphorylation converts eIF2 from the substrate of the GDP/GTP exchange reaction into an inhibitor (eIF2-P) of eIF2B. This conversion reduces the cell’s TC levels and induces the widespread reprogramming of translation known as the Integrated Stress Response (ISR). Here, we chart an allosteric axis running through eIF2B, revealing the importance of a protrusive α-helix in its β-subunit, the ‘latch-helix’, that locks onto the α-subunit to induce eIF2B activity. eIF2-P binding unhooks the latch-helix, opening eIF2B, which inhibits its GDP/GTP exchange activity. Distinct viral proteins have convergently evolved to bind to eIF2Bα and stabilize the latch-helix-bound active state. Using these insights, we generated <u>ISR</u>-<u>ACT</u>ivating compounds, ISRACTs, that stabilize eIF2B in its inhibited, unlatched state. Our study thus highlights how state-transitions in eIF2B are regulated via long-range allostery.

## Introduction

The eukaryotic translation initiation factor 2B (eIF2B) is a guanine nucleotide exchange factor (GEF) and gatekeeper of protein synthesis. It is a complex comprised of two tetramers containing the β-, γ-, δ-, and ε-subunits that further assemble into a decamer by binding the homodimeric α-subunit (*1–5*). When fully assembled, the eIF2B decamer catalyzes GDP ejection from its substrate, the heterotrimeric eukaryotic initiation factor 2 (eIF2; composed of α-, β-, and γ-subunits) (*4–7*), allowing for GTP binding to eIF2γ. By catalyzing eIF2 nucleotide exchange, eIF2B facilitates the formation of the eIF2-GTP-methionyl-initiator tRNA “Ternary Complex” (TC). TC formation is a prerequisite for mRNA translation initiation as it delivers the first aminoacyl-tRNA to 40S ribosomal subunits scanning for AUG initiation codons (*8–12*). Stress-inducing perturbations, such as viral infection (*13*, *14*), ribosome stalling and collisions (*15*, *16*), secretory protein folding (*17*, *18*) and trafficking imbalances (*19*), redox changes (*20*, *21*), mitochondrial dysfunction (*22*, *23*), hypoxia (*24*, *25*), and amino acid/nutrient deficiencies (*26–32*), cause the activation of any of four eIF2 kinases (PKR, GCN2, PERK, and HRI). These kinases all phosphorylate eIF2α at a single site, serine 51, producing eIF2-P – a potent eIF2B inhibitor. This conversion of eIF2B’s substrate, eIF2, into its inhibitor, eIF2-P, is the hallmark of the Integrated Stress Response (ISR) (*7*, *21*, *33–37*). As eIF2-P accumulates, TC levels diminish, which leads to a decline in translation initiation rates for all AUG-initiated transcripts.

While the ISR is characterized by a decline in general translation, a subset of transcripts with inhibitory upstream open reading frames (uORFs) are translated at higher rates, leading to enhanced accumulation of their protein products (*31*, *38–41*), such as the transcriptional activator *ATF4* (*39*, *42*, *43*). The translation of these uORF-containing transcripts and the resulting transcriptomic changes enable cells to adapt to and counteract the stressful perturbation (*39*, *44–47*). This broad reprogramming of protein synthesis rates is typically cytoprotective (*48*, *49*), but chronic activation of the ISR can become maladaptive and is associated with pathologies that include leukodystrophy (*50–54*), cancer (*24*, *55–57*), and cognitive impairments caused by Alzheimer’s disease (*58–62*), traumatic brain injuries (*63*), and Down syndrome (*64*).

In these disease contexts, the drug-like small molecule ISRIB (*41*, *49*) and its clinical derivatives maintain protein synthesis rates despite elevated eIF2-P levels. Mechanistically, ISRIB binds to eIF2B in a two-fold symmetric pocket formed at the interface between two eIF2Bβ and two eIF2Bδ subunits proximal to the substrate eIF2 binding site. In doing so, ISRIB stabilizes the active “A”-state of the eIF2B decamer (*4*, *5*, *65–67*). The A-state is defined, in part, by the distance between the β- and δ-subunits from opposing tetramers. When the cleft between these subunits is ∼32 Å wide, the α-subunit of eIF2 binds favorably and this positions the GDP-bound γ-subunit over the catalytic eIF2B ε-subunit to drive nucleotide exchange. The inactive “I”-state of eIF2B is induced by eIF2-P binding, which occurs at a distinct allosteric site between the α- and δ-subunits more than 40 Å away from the ISRIB and eIF2 binding sites (*36*, *37*). The I-state is characterized by widening of the eIF2α binding pocket to ∼35 Å, reducing substrate binding affinity and guanine nucleotide exchange (*65*, *66*).

ISRIB thus renders eIF2B resistant to inhibition by eIF2-P through allosteric stabilization of the A-state (*65–67*). Although there are no known endogenous or exogenous ligands that engage the ISRIB pocket, viruses inhibit the ISR via effector proteins that engage at other sites on eIF2B (*68–71*). Viral suppression of the ISR promotes synthesis of viral proteins despite activation of the innate sensors that lead to eIF2α phosphorylation (*13*, *14*). The best-characterized example to date comes from studies of the Sandfly fever Sicilian virus (SFSV) protein, NSs (*68–70*). NSs binds to eIF2B at a site that overlaps with the eIF2-P inhibitor binding site, thereby sterically blocking eIF2-P binding. However, rather than inducing the I-state as eIF2-P does, it allosterically promotes the A-state conformation (*67*, *69*, *70*).

These observations inform current models of how eIF2-P activates a long-range allosteric change through the eIF2B core to regulate eIF2B GEF activity, including the discovery of the eIF2Bδ subunit’s fulcrum-helix, which acts as a switch for propagating changes in the angle between the eIF2Bα and eIF2Bδ subunits (*67*). However, our understanding of how information is relayed between the site of eIF2-P inhibitor binding, the site of substrate binding, and the ISRIB-binding pocket in eIF2B is incomplete. Furthermore, how diverse eIF2B ligands—including pharmaceuticals and pathogen effector proteins—influence allosteric chain linkage despite their disparate and distant binding sites remains poorly understood. Using structural, biochemical and pharmacological approaches, we map the axis of coupled allosteric changes that transmits information over the 40 Å distance between eIF2B’s diverse binding sites. Our analyses reveal a remarkable convergence in viral evolution where unrelated viruses utilize dissimilar proteins to stabilize the A-state of eIF2B. We leverage the new structural understanding of allosteric eIF2B regulation to develop small molecules, ISRACTs, that target the ISRIB binding pocket to activate, rather than inhibit, the ISR.

## Results

### A short α-helix in eIF2Bβ allosterically competes with eIF2α-P binding

To gain mechanistic understanding of the conformational dynamics that govern eIF2B activity, we segmented and classified 569,415 single-particle cryo-EM images of eIF2B in its apo-form and subjected the images to 3D variability analysis (3DVA) (Table 1). Visualization of 3DVA highlighted motion trajectories of eIF2B consistent with the particle distributions (Fig. S1 and Movie S1). The trajectories included (1) the A- to I-state opening of the eIF2 substrate binding pocket between the two eIF2Bβγδε tetramers (Fig. 1A; Motion 1), (2) rotation of eIF2Bα away from eIF2Bβ/δ (Motion 2), and (3) swiveling of the eIF2Bγ beta-solenoid domains perpendicular to the tetramer rocking motion (Motion 3).

**Fig 1.**
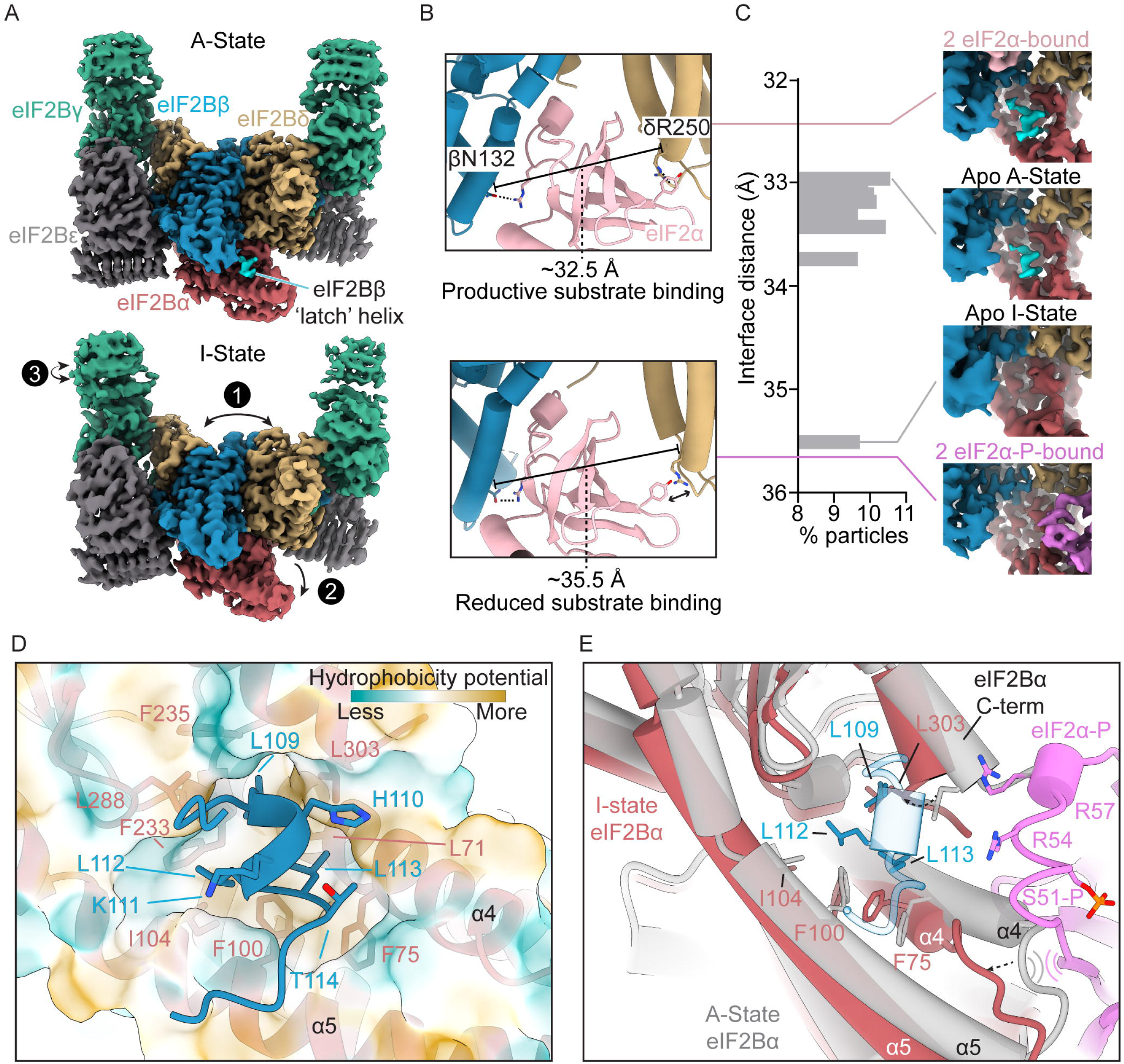
Disordering of a helical latch that bridges eIF2Bβ and eIF2Bα demarcates the inactivation of eIF2B. (A) (Top) Cryo-EM density of wild-type eIF2B(αβγδε)_2_ in the apo, A-state with the location of the latch-helix highlighted in cyan. (Bottom) Cryo-EM density of eIF2B in an apo, I-state where the latch-helix is no longer resolvable. Direction of three trajectories of motion identified by 3D variability analysis are indicated by **1**, **2**, and **3**, respectively (see Supp movie S1) (B) Zoom-in on the eIF2 binding interfaces defined by residues βN132 and δ’R250 (‘ indicates a subunit from the opposing half of eIF2B) from the eIF2 substrate-bound (PDB: 6O81, top) and the eIF2α-P inhibitor-bound (PDB: 9Y5R, bottom) structures with the model of eIF2α (pink) from 6O81 overlaid. (C) (Left) Distribution of eIF2B particles after 3D classification, as measured by the distance between βN132 and δ’R250 in structures from 8 distinct 3D classes. (Right) Close-up view of the eIF2B α-β interface from the cryo-EM structures of eIF2-eIF2B (EMD-0649), eIF2α-P (EMD-72523), and representative A-state and I-state structures from 3D classification. (D) Zoom-in of the latch-helix and its binding pocket with a transparent surface representation of eIF2Bα colored by molecular hydrophobicity potential. Key residues of eIF2Bα and eIF2Bβ are labeled according to the color of each subunit. (E) The structural model of eIF2α-P (pink)-bound eIF2Bα (maroon) overlaid with a transparent model of A-state apo eIF2Bα (gray) and eIF2Bβ latch-helix (blue). Conformational changes of interest are indicated with dashed arrows and potential steric clashes indicated by curved lines.

To estimate the probability of particle distribution within these states, and thus the energy barriers associated with conformational transitions, we performed heterogeneous 3D classification of all particles into 10 classes. This procedure yielded 8 distinct reconstructions and 2 noise classes. Unexpectedly, we found that ∼10% of the particles yielded eIF2B completely in the I-state conformation, with (i) the substrate interfaces as far apart as eIF2-P bound eIF2B (distance of 35.5 Å versus 32.5 Å observed in the substrate-bound A-state, Fig. 1B-C), (ii) the ISRIB-binding pocket lengthened by 4.3 Å (Fig. S2A), (iii) the eIF2Bδ “fulcrum-helix”, or switch-helix (*67*), positioned in its rotational I-state (Fig. S2B), and (iv) the dimer/dimer interface histidine βH160 in its resting I-state-characteristic rotamer position that precludes the A-state-stabilizing interface-zipper interaction (Fig. S2C; Movie S2), as first suggested in an I-state-inducing point mutant, βH160D (*72*).

Further inspection of the minority I-state fraction of eIF2B revealed missing density for a protrusive α-helix at the tip of eIF2Bβ. In the A-state, this short α-helix, henceforth referred to as the “latch-helix”, bridges eIF2Bβ to eIF2Bα (colored in *cyan*, Fig. 1A, 1C; Movie S1). Density for the latch-helix is absent in reconstructions of the eIF2-P-bound I-state (Fig. 1A - C). These observations suggest that latch-helix binding to eIF2Bα could be a driving force in the inactive (I-) to active (A-) state transition that “locks” eIF2B in the A-state and hence must unhook to allow eIF2B to relax into the I-state. Notably, we observed two 3D classes of eIF2B with defined density for the latch-helix only on one half of the decamer; however, these classes are otherwise in the A-state (Fig. S2D, Table 2). Thus, latch-helix ordering on only one of the two halves of eIF2B is sufficient to drive the I- to A-state transition across the rest of the eIF2B complex.

The latch-helix comprises eIF2Bβ residues ^108^SLHKLLT^114^ within a flexible loop linking two helices in the eIF2Bβ core (Fig. 1D). Latch-helix side chains of βL109, βL112, and βL113 nestle into a hydrophobic pocket in eIF2Bα burying a surface area of approximately 623 Å^2^. Within the pocket, each latch-helix leucine ‘bump’ occupies a unique ‘hole’ created by the specific orientation of eIF2Bα residues, highlighting the importance of both the hydrophobic effect and geometric complementarity as drivers of this interaction. A reanalysis of published hydrogen-deuterium exchange coupled to mass spectrometry (HDX-MS) data (*67*) confirmed that the latch-helix is more solvent protected in the bound A-state than in the unbound eIF2-P-induced I-state (Fig. S3).

Comparison of the latch-helix binding pocket in A-state eIF2B (*grey*) with an eIF2α-P-bound I-state eIF2B (*red*) reveals the determinants of eIF2α-P-induced latch-helix release (Fig. 1E). Specifically, the pocket occupied by latch-helix amino acid βL109 is occluded by the C-terminus of eIF2Bα rotating inward to place αL303 in this space. Its position is stabilized by the A- to I-state rearrangement of the adjacent fulcrum-helix (*67*) (Movie S3). The rearrangement of eIF2Bα helices α4 and α5 and rotation of αF75 remodels the pocket further and precludes binding of latch-helix amino acids βL112 and βL113. Accordingly, latch-helix and eIF2-P binding are mutually exclusive. Moreover, interaction of eIF2α-P sterically blocks eIF2Bα from adopting the A-state conformation. Together, these structures demonstrate how latch-helix engagement and the linked movements in eIF2Bα govern eIF2B activity.

### Deletion of the latch-helix deactivates eIF2B allosterically and induces the ISR

Next, we asked how the latch-helix regulates the conformation and catalytic GEF activity of eIF2B. We observed that deletion of the latch-helix (ΔeIF2Bβ_100-116_, hereafter designated as eIF2B^Δ-latch^) resulted in an ∼8-fold decrease in eIF2B GEF activity as measured by *in vitro* nucleotide exchange (t_1/2_WT = 2.3 +/- 0.3 min vs. t_1/2_Δ-latch = 18.4 +/- 2.4 min) (Fig. 2A). We found no difference in the sedimentation velocity between eIF2B^WT^ and eIF2B^Δ-latch^ (Fig. 2B), indicating that the loss of catalytic activity was not due to defective eIF2B decamer assembly. We then assessed how eIF2B^Δ-latch^ affects ISR activation *in cellula*. To this end, we depleted endogenous eIF2Bβ with small interfering RNA (siRNA) (Fig. 2C) and expressed siRNA-resistant epitope-tagged *eIF2Bβ^WT^* or *eIF2Bβ^Δ-latch^* (Fig. 2D). In line with the *in vitro* results, eIF2Bβ-depleted cells expressing eIF2Bβ^Δ-latch^ exhibited a robust ISR, indicated by ATF4 accumulation, whereas eIF2Bβ^WT^ expressing cells did not (Fig. 2D). Together, the *in vitro* and *in cellula* experiments demonstrate that the engagement of the latch-helix is required for optimal eIF2B GEF activity.

**Fig 2.**
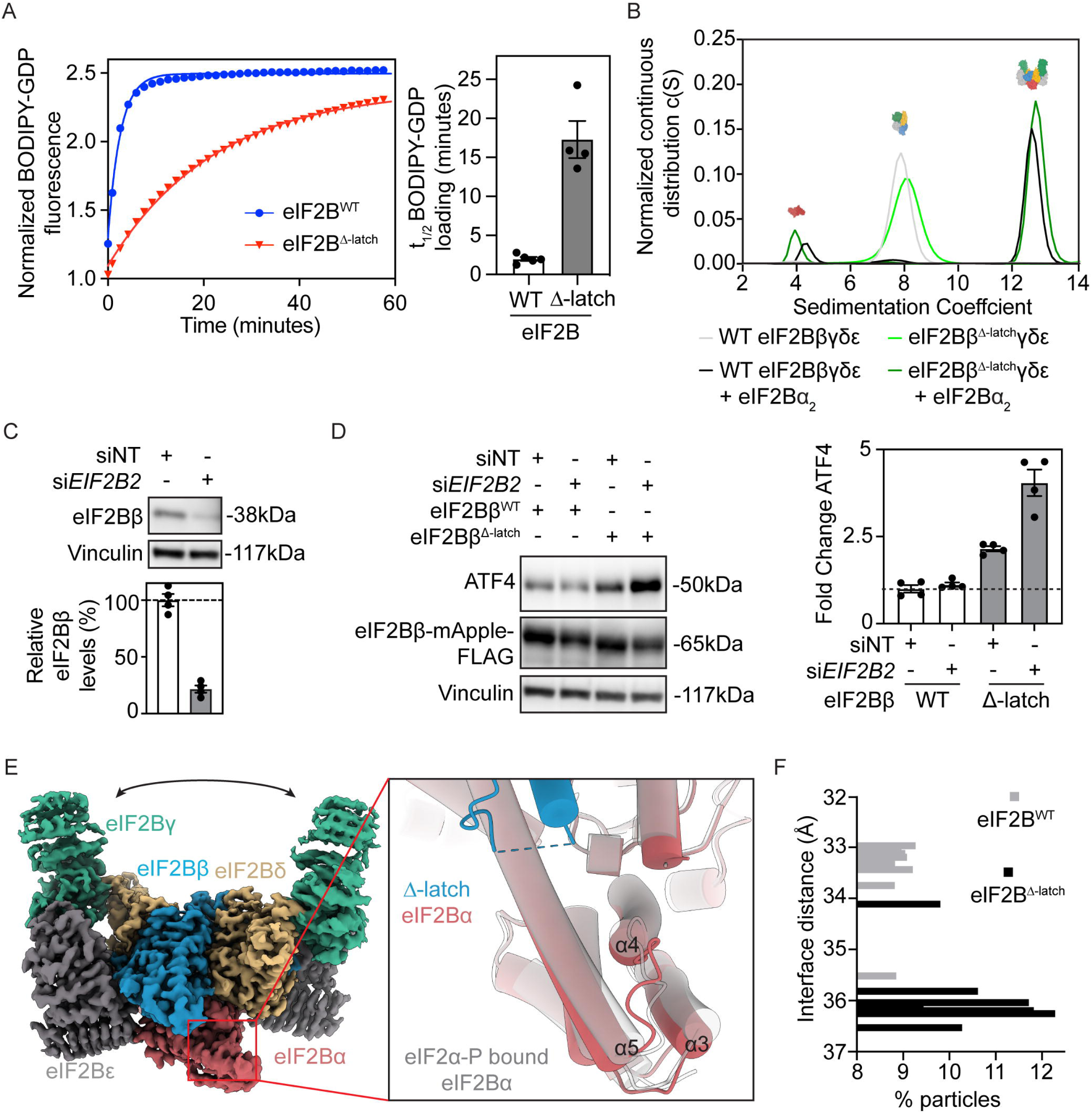
Removal of the eIF2Bβ latch-helix is sufficient to inactivate eIF2B. (A) Guanine nucleotide exchange factor (GEF) activity as measured by BODIPY-GDP loading onto eIF2, using 10 nM wild-type (eIF2B^WT^) or β100-116 deletion (eIF2B^Δ-latch^) eIF2B as the enzyme and 100 nM eIF2 as the substrate. The left panel shows a representative time course readout of a 60-minute nucleotide exchange reaction. The t_1/2_ of nucleotide loading from three experiments are plotted and averaged in the right panel. Data represent the mean±s.e.m (n**≥**4). (B) Sedimentation velocity of WT and Δ-latch eIF2Bβγδε in the presence or absence of eIF2Bα_2_ as measured by analytical ultracentrifugation. (C) Lysates of HEK293T cells transfected for 72 hours with either non-targeting (siNT) or *EIF2B2*-targeting (si*EIF2B2*) siRNA were subjected to immunoblotting using antibodies to eIF2Bβ and vinculin. Densitometry of the eIF2Bβ blot intensity normalized to the intensity of Vinculin and plotted relative to the non-targeting control. Data represent the mean±s.e.m (n=4). (D) Immunoblots and quantifications of lysates of HEK293T cells co-transfected with either non-targeting (siNT) or *EIF2B2*-targeting (si*EIF2B2*) knockdown siRNA and rescue plasmids expressing siRNA resistant and mApple-FLAG tagged *eIF2Bβ^WT^*or *eIF2Bβ^Δ-latch^*. ATF4 mean fold changes are normalized to the amount of Vinculin measured by densitometry. Data represent the mean±s.e.m (n=4). (E). The I-state cryo-EM structure of eIF2B^Δ-latch^ (left) with a zoom-in view of the eIF2Bα latch-helix binding pocket in red overlaid with the eIF2α-P bound I-state conformation in transparent grey (right). (F) Distribution of eIF2B^WT^ (gray) and eIF2B^Δ-latch^ (black) particle conformations as measured by the distance between residues βN132 and δ’R250 from structural models fit into cryo-EM maps generated by 3D classification.

Cryo-EM analysis allowed us to obtain a consensus structure of the eIF2B^Δ-latch^ decamer at 2.1 Å (Fig. S4A-E). Comparison of the latch-helix binding pocket from this structure to the I-state eIF2α-P-bound eIF2B^WT^ structure showed that eIF2B^Δ-latch^ adopts the I-state (Fig. 2E) with eIF2Bα and the eIF2Bδ fulcrum-helix in their characteristic I-state conformations (Fig. S4F). Consequently, the eIF2α substrate binding pocket opens by ∼3 Å (Fig. 2F), readily explaining the loss of GEF activity described above (Fig. 2A-D).

We next 3D-classified the refined particles to probe the conformational spectrum of eIF2B^Δ-latch^. Reminiscent of the 3D classification results for the eIF2B^WT^ particles (see Fig. 1C), we found that a small fraction of eIF2B^Δ-latch^ particles (∼12%) classified into the opposing state (A-like) from the majority consensus (I-state) (Fig. 2F). Contrasting this minor class to a canonical A-state structure, we find that while the hallmarks such as the eIF2α substrate-binding cleft distance and the fulcrum-helix are in an A-state conformation, the latch-helix pocket of eIF2Bα remains in the I-state in the absence of latch-helix engagement (Fig. S4E).

Importantly, comparison of the eIF2B^Δ-latch^ α subunit conformation with multiple crystal structures of eIF2Bα [(*73*, *74*); PDB: 3ECS, 7KMA] shows that eIF2Bα adopts the I-state conformation as its *in crystallo* ground state (Fig. S5C). These observations indicate that eIF2B decamer assembly itself does not induce the A-state unless and until the latch-helix locks onto eIF2Bα. Thus, in the absence of the binding energy provided by the eIF2Bβ latch-helix, eIF2Bα remains in its low-energy ground state. In turn, the ground state of eIF2Bα favors the open and inhibited state of eIF2B, which induces the ISR even in the absence of eIF2-P.

### Viral effector proteins convergently evolved to suppress the ISR

The results presented so far indicate that during ISR activation, eIF2-P binding promotes unbinding of the eIF2Bβ latch-helix by engaging with eIF2Bα (Fig. 1). By contrast, the Sandfly fever Sicilian virus (SFSV) effector protein NSs hijacks the eIF2-P binding pocket between eIF2Bα and eIF2Bδ (*69*, *70*) but stabilizes an eIF2B A-state, suggesting a conformation complicit with latch-helix ordering. Indeed, inspection of the cryo-EM structures of NSs (*69*, *70*) (EMD-24535, EMD-31472) bound to eIF2B confirmed the presence of an ordered eIF2Bβ latch-helix (Fig. S6).

We next considered if proteins encoded by other viruses might mimic the mechanism exhibited by SFSV NSs to stabilize the latch-helix-bound state of eIF2Bα and eIF2Bβ to suppress the ISR. For example, the Beluga whale coronavirus accessory protein 10 (Bw CoV-AcP10) was shown to bind to eIF2B and compete with eIF2-P (*71*). AcP10, however, shares no sequence similarity with NSs, and the mechanism by which it inhibits the ISR remained unknown. To address this knowledge gap, we expressed an N-terminal MBP-tagged version of AcP10 in *E. coli* (Fig. S7A-B) and tested the purified protein to see if it bound to eIF2B directly. Pulldown assays with recombinant eIF2Bα dimers (α_2_), eIF2B tetramers (βγδε), or eIF2B decamers (αβγδε)_2_ *in vitro* indicated that MBP-AcP10 bound directly only to fully assembled eIF2B decamers (Fig. 3A), indicating that a competent AcP10-binding interface requires decamer assembly.

**Fig 3.**
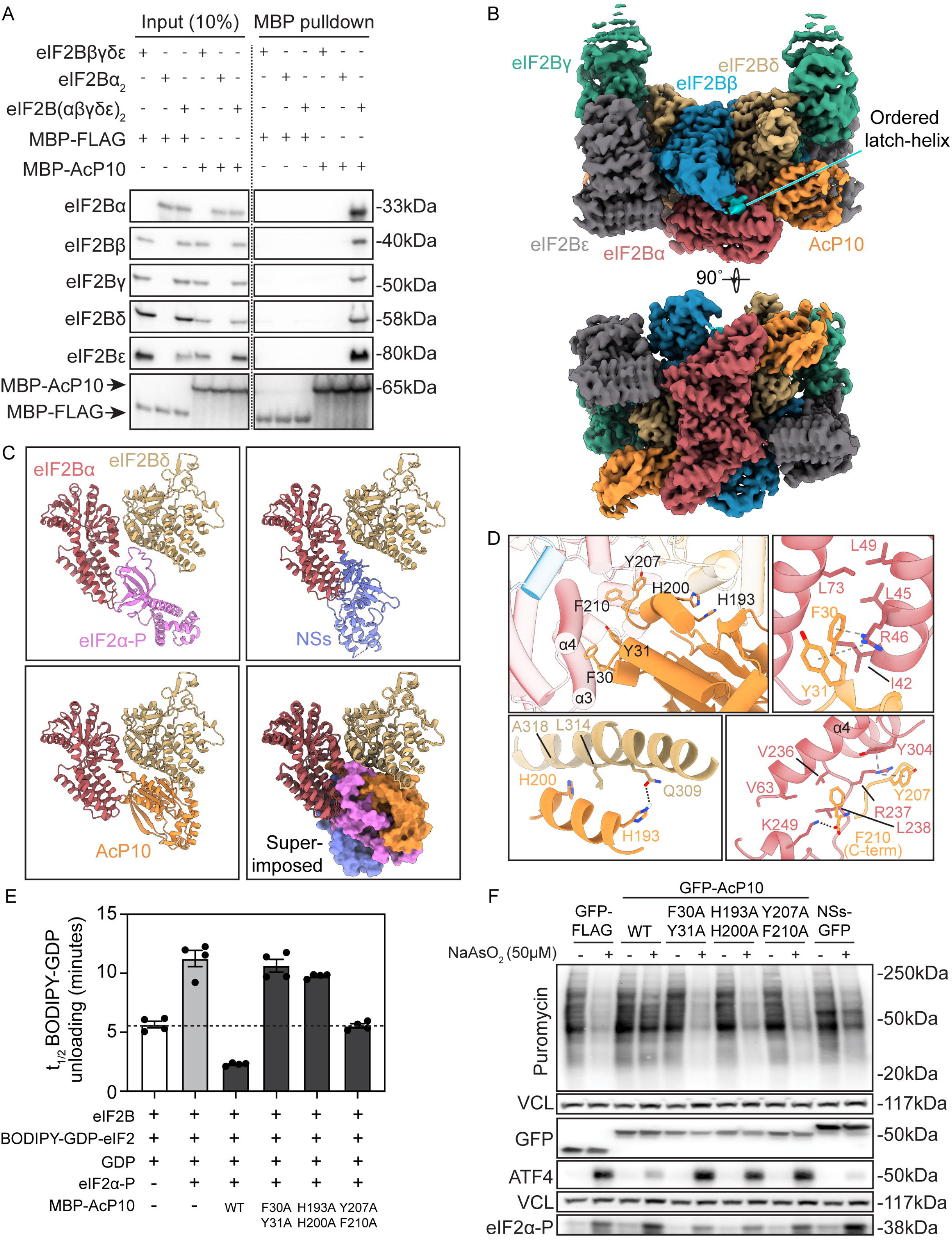
Viral proteins AcP10 and NSs convergently evolved as activators of eIF2B that order the latch-helix and outcompete eIF2-P to suppress the ISR. (A) *In vitro* pulldowns of purified eIF2Bα_2_ (200 nM), eIF2Bβδγε (400 nM) or eIF2B(αβδγε)_2_ (200 nM) with MBP-FLAG (1 µM) and MBP-AcP10 (1 µM). Immunoblotting was performed using antibodies to eIF2B subunits and MBP. Data are representative of three independent experiments. (B) Cryo-EM density of the AcP10-eIF2B^WT^ complex showing two copies of AcP10 (orange) bound. (C) Comparison of the binding locations on eIF2Bα/δ occupied by either eIF2α-P (top left), NSs (top right) or AcP10 (bottom left). The bottom right panel depicts the overlay of the interface. eIF2Bα - red; eIF2Bδ - yellow; eIF2α-P - violet; NSs - blue; AcP10 - orange. (D) Zoomed-in view of the entire AcP10-eIF2B interaction interface (top left) with key interactions made by AcP10 residues (clockwise from top right): F30/Y31 with eIF2Bα, Y207/F210 with eIF2Bα, and H193/H200 with eIF2Bδ. Hydrogen bond, electrostatic, or cation–π interactions are denoted by dashed lines. AcP10 - orange; eIF2Bα - red; eIF2Bδ - yellow; eIF2Bβ latch-helix - light blue. (E) GEF activity as measured by BODIPY-GDP unloading from eIF2 (100 nM) by eIF2B^WT^ decamers (10 nM) with or without the addition of eIF2α-P (1 µM), MBP-AcP10 WT (1 µM) or MBP-AcP10 interface mutants (1 µM), as indicated. Shown are mean±s.e.m. of t_1/2_ values derived from a one-phase decay fit of four replicates. (F) HEK293T cells were transfected for 24 hours with plasmids expressing GFP-tagged AcP10, AcP10 interface mutants or NSs and treated with sodium arsenite (50 µM) for 4 hours to induce the ISR. For a set of experiments, cells were subjected to puromycin pulse-chase analyses to assess nascent protein synthesis rates. Immunoblotting of lysates was performed using antibodies to puromycin, vinculin, GFP, ATF4 and eIF2α-P. Puromycin pulse-chase (n=2) and ATF4/eIF2α-P accumulation (n=4) assay samples were processed in parallel on separate gels. Representative blots are shown.

To determine the interaction interface between AcP10 and eIF2B decamers, we solved a 2.9 Å structure of this complex by cryo-EM (Fig S7). The interface was well-resolved (Fig. 3B, Fig. S7C-F), allowing us to model two molecules of AcP10 bound to eIF2B. Like the structurally unrelated SFSV NSs, AcP10 bound at the eIF2-P binding pockets, forming symmetric interfaces between eIF2Bα and eIF2Bδ on either side of the decamer (Fig. 3B-C) (*69*, *70*) (Fig. S7). Cross-linking mass spectrometry confirmed that AcP10 interacts only with eIF2Bα and eIF2Bδ (Fig S8). 3D classification revealed that ∼24% of the particles were bound by one molecule of AcP10, while the remaining particles had two AcP10 molecules bound. As expected and akin to NSs, the AcP10-eIF2B structure exhibited an ordered eIF2Bβ latch-helix (Fig. 3B) and the fulcrum-helix in its A-state conformation (Fig. S7F). These hallmarks are indicative of an A-state-stabilized eIF2B with a productive eIF2α substrate-binding interface.

An intricate set of interactions between eIF2Bα and AcP10 anchors this viral effector to eIF2B. AcP10 adopts a uridine-cytidine kinase fold, comprising a core of five-stranded parallel β-sheets flanked on both sides by a pair of α-helices and additional α-helices sitting above the core sheet structure together with a large β-hairpin loop. eIF2B is engaged by AcP10 in a wedge-like manner by inserting its N and C-termini in-between the concave surface formed by subunits eIF2Bα and eIF2Bδ (Fig. 3D, S9A). A stabilizing edge-to-face π-π interaction between the aromatic rings of AcP10 F30 and Y31 intercalates F30 into the hydrophobic groove formed in eIF2Bα between helices α3 and α4. The AcP10 aromatic F30-Y31 dyad further engages eIF2Bα via a strong cation-*π* interaction with the guanidinium moiety of αR46 on helix α3 (Fig. 3D; top right panel). These interactions, in turn, stabilize the opposite face of helix α4, allowing for strong interaction with the latch-helix.

A second aromatic dyad within AcP10, composed of Y207 and F210 at the C-terminus, also latches onto eIF2Bα’s C-terminal helix (Fig. 3D; bottom right panel). These interactions likely prevent the eIF2Bα C-terminus from moving inward, thereby stabilizing a conformation of the latch-helix pocket in eIF2Bα that is conducive for latch-helix binding. Additionally, the AcP10 C-terminal helix packs parallel to the longest helix in eIF2Bδ forming a large interface stabilized by H-bonds and the hydrophobic effect (Fig. 3D; bottom left panel).

A salient feature of the AcP10-eIF2B interaction is the use of ‘aromatic fingers’ that intercalate in between helices α3 and α4 of eIF2Bα (Fig. 3D, S10). This arrangement is reminiscent of interactions observed between NSs and eIF2B. However, diverging from AcP10, NSs projects both sets of its aromatic fingers on either side of the eIF2Bα helix α3, effectively sandwiching this helix. AcP10, by contrast, extends a pair of aromatic fingers to contact the surface patch of eIF2Bα just next to eIF2Bδ, as well as a set of polar residues to contact eIF2Bδ. Indeed, AcP10 buries ∼621 Å^2^ and ∼757 Å^2^ of solvent accessible area against subunits eIF2Bα and eIF2Bδ, respectively, while NSs buries ∼995 Å^2^ and ∼149 Å^2^, explaining their high binding affinities to eIF2B.

To probe the influence of the eIF2B-contacting amino acids, we purified AcP10 mutants carrying alanine substitutions on key residues and measured their ability to bind eIF2B decamers *in vitro*. Pairwise mutations to AcP10 F30/Y31 and H193/H200, resulted in a strong loss of binding to eIF2B (Fig. S9B; lanes 9-10), while mutations to AcP10 M196 and Y207/F210 had partial effects (Fig. S9B; lanes 11-12). An AcP10 D29A mutation had no effect on binding. Next, we explored the functional importance of these residues by measuring competitive eIF2B binding between AcP10 and eIF2α-P. First, we titrated AcP10 against pre-formed BODIPY-GDP-eIF2 complexes in the presence or absence of eIF2α-P and measured nucleotide exchange *in vitro*. As expected, eIF2B GEF activity was strongly inhibited by eIF2α-P [Fig. 3E - condition 2; S9C - condition 6, (*65*, *66*)]. The inhibition was completely reversed upon addition of AcP10 (Fig. 3E - condition 3; S9C - conditions 9-10). In accordance with our binding assays, AcP10 carrying F30A/Y31A or H193A/H200A mutations did not rescue the inhibitory effect of eIF2α-P, while mutations to AcP10 Y207/F210 had subtle effects (Fig. 3E). We note here that even in the absence of eIF2α-P, AcP10 stimulated eIF2B GEF activity (Fig. S9C, conditions 1-5). This is consistent with our observation that apo-eIF2B continuously samples the A- and I-state conformations (Fig. 1, S1), leading us to reason that AcP10 binding strongly shifts the equilibrium to the A-state. Supporting this notion, 3D variability analyses and multi-class cryo-EM reconstructions of eIF2B bound to AcP10 yielded no particles in the I-state (Fig. S7).

Second, we pre-incubated streptavidin-tagged eIF2B decamers with AcP10^WT^ or AcP10^H193A/H200A^ and then added purified eIF2α-P to the respective mixtures. We affinity purified eIF2B-bound complexes and determined the relative amounts of eIF2α-P or AcP10 that co-purified with eIF2B by immunoblotting. As expected, in reaction mixtures containing only eIF2B and eIF2α-P, eIF2B bound efficiently to eIF2α-P (Fig. S9D, lane 4). In reaction mixtures containing eIF2B, AcP10^WT^ and eIF2α-P, eIF2B robustly bound to AcP10 but failed to efficiently co-purify eIF2α-P (Fig. S9D, lane 5). AcP10^H193A/H200A^ did not co-precipitate with eIF2B efficiently and did not prevent the interaction between eIF2α-P and eIF2B (Fig. S9D, lane 6), corroborating the stabilizing role of these polar residues in the AcP10-eIF2B interface (*71*).

Third, we transfected GFP-tagged versions of NSs, AcP10, or the AcP10 interface mutants into HEK293T cells and tested their ability to inhibit the ISR induced by sodium arsenite, a commonly used activator of the ISR-inducing kinase HRI (*20*, *75*). AcP10 and NSs, but not the AcP10 interface mutants, strongly inhibited the accumulation of ATF4 and restored nascent protein synthesis rates even as eIF2-P levels remained high (Fig. 3F).

Together, these results indicate a remarkable convergence of evolutionary constraints that have shaped how structurally unrelated effectors from distinct viruses inhibit the ISR. BwCoV AcP10, like SFSV NSs, leverages molecular interactions involving ‘aromatic fingers’ combining strong hydrophobic and electrostatic components to engage the same ‘hotspot’ groove between helices α3 and α4 of eIF2Bα (Fig. S10). In doing so, both AcP10 and NSs hold eIF2Bα in the A-state with an open pocket for latch-helix binding while occluding eIF2-P binding.

### AcP10 compensates for the loss of the eIF2Bβ latch-helix

We next tested whether AcP10 binding could restore the enzymatic function of eIF2B^Δ-latch^. To this end, we first established that AcP10 binds to both eIF2B^WT^ and eIF2B^Δ-latch^ decamers (Fig. S11), indicating that, as expected, the latch-helix is not required for AcP10 binding. We then incubated wild-type AcP10 or interface mutants of AcP10 with eIF2B^Δ-latch^ decamers and assayed for enzymatic function *in vitro*. Remarkably, AcP10 accelerated the guanine-nucleotide exchange rate catalyzed by eIF2B^Δ-latch^ (Fig. 4A; t_1/2_Δ-latch = 15.7 +/- 0.2 min vs. t_1/2_Δ-latch + AcP10 = 4.8 +/- 0.7 min). This restorative effect was lost in reactions incubated with the AcP10 interface mutants (Fig. 4A). Corroborating these data, GFP-AcP10, but not GFP-AcP10^F30A/Y31A^, robustly reversed ATF4 accumulation in *EIF2B2* depleted cells expressing eIF2Bβ^Δ-latch^ (Fig. 4B-C). Therefore, AcP10 binding compensates for the loss of the latch-helix and inhibits the eIF2B^Δ-latch^-induced ISR.

**Fig 4.**
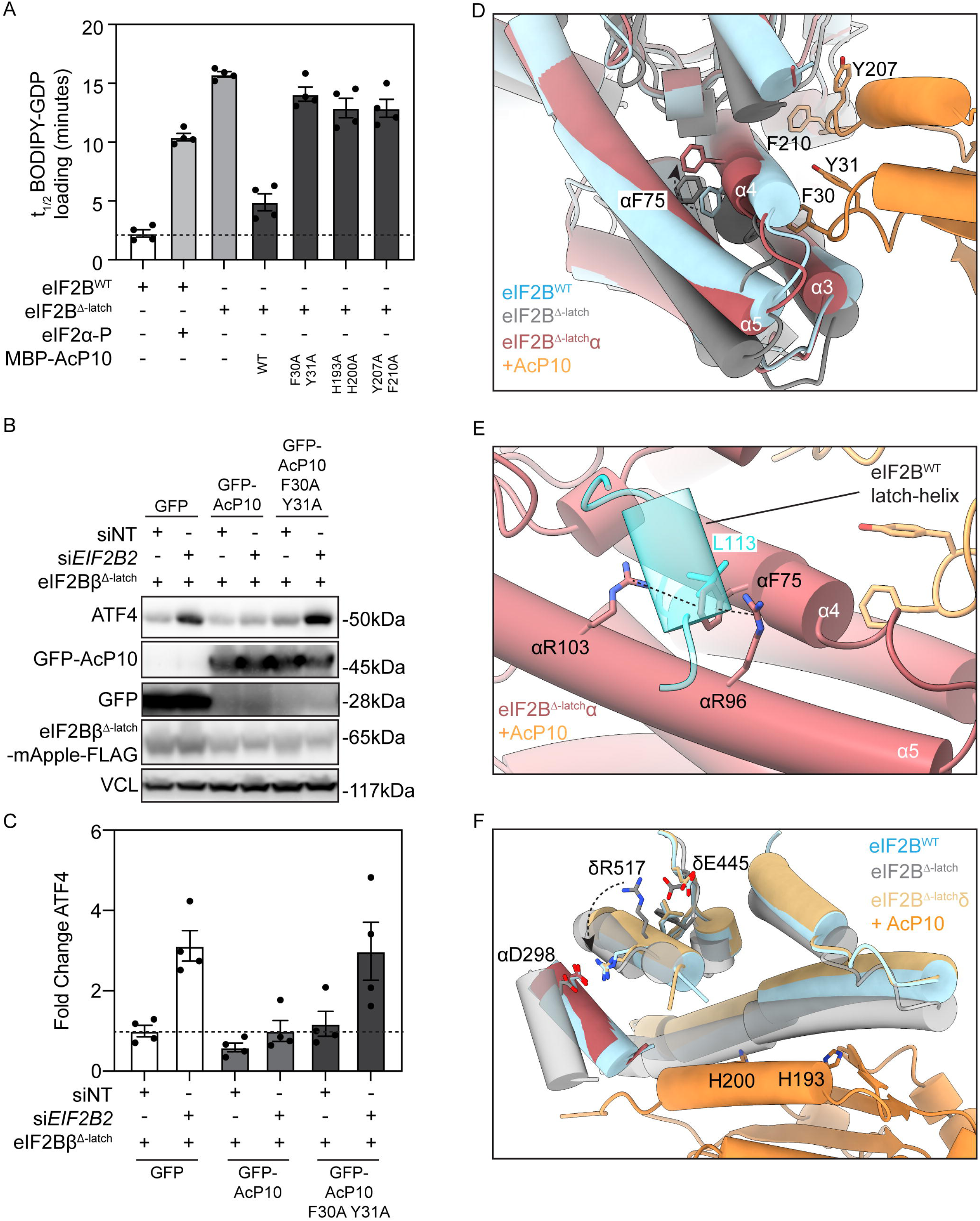
AcP10 rescues the activity of eIF2B^Δ-latch^ and suppresses ISR activation in eIF2B^Δ-latch^ mutant cells. (A) GEF activity as measured by BODIPY-GDP loading onto eIF2 (100 nM) by eIF2B^WT^ (10 nM) with or without the addition of eIF2α-P (1 µM) or eIF2B^Δ-latch^ decamers (10 nM) with or without the addition of MBP-AcP10 WT (1 µM) or MBP-AcP10 interface mutants (1 µM), as indicated. Shown are mean±s.e.m. of t_1/2_ values derived from a one-phase association fit for four experimental replicates. (B) Immunoblots of HEK293T cell lysates co-transfected with either non-targeting (siNT) or *EIF2B2*-targeting (si*EIF2B2*) siRNA and rescue plasmids expressing siRNA resistant and mApple-FLAG tagged *eIF2Bβ^Δ-latch^*. (C) Fold changes in the levels of ATF4 normalized to the amount of Vinculin in cells by densitometry. Data represent the mean±s.e.m (n=4). (D - F) The structural models of eIF2Bα (top and middle panels) and eIF2Bδ (bottom panel) when AcP10 (orange) is bound to eIF2B^Δ-latch^ (multi-colored) overlaid with A-state apo eIF2B^WT^ (blue) and I-state apo eIF2B^Δ-latch^ (gray) with the latch-helix highlighted (cyan). Conformational changes of interest are indicated with dashed arrows and denote I- to A-state-transition. The cation–π–cation interaction in (E) is denoted with dashed lines.

To characterize this reaction, we determined the AcP10-eIF2B^Δ-latch^ complex structure by cryo-EM (Fig. S12A-D). These maps revealed that AcP10 bound to eIF2B^Δ-latch^ through the eIF2-P binding pocket in the same manner as it did to eIF2B^WT^ (Fig. S12C). Structures of apo-eIF2B^Δ-latch^ overlaid onto AcP10-eIF2B^Δ-latch^ indicated a pronounced set of conformational changes in eIF2B^Δ-latch^ upon AcP10 binding (Fig. 4D-F). First, AcP10 induced a narrowing of the eIF2B^Δ-latch^ substrate binding pocket from 36 Å to 33 Å, conducive to productive eIF2 substrate binding (Fig. S12E). Second, by seizing hold of the groove between helices α3 and α4 of eIF2Bα, AcP10 reorients eIF2Bα to adopt an A-state position relative to eIF2Bβ and δ (Fig. 4D). However, the eIF2Bα C-terminal helix does not fully reset the latch-helix binding pocket to the wild-type A-state conformation. Instead, the aromatic ring of αF75 shifts to occupy the hydrophobic pocket of eIF2Bα that remains solvent-accessible upon loss of the latch-helix, thereby mimicking the A-state stabilizing environment created by the latch-helix residue βL113 (Fig 4D-E). This compensatory shift of αF75 is further stabilized by the sidechain of αR96 that reorients to sandwich the phenyl ring of αF75 in a cation-***π***-cation interaction with αR103 (Fig. 4E). Third, the fulcrum-helix on eIF2Bδ rotates back to the A-state conformation where δR517 engages in a salt bridge with αD298 (Fig. 4F).

Upon analysis of particles by 3D classification, we noted that an AcP10:eIF2B^Δ-latch^ ratio of 2:1 in solution yielded ∼35% of particles that were bound by two AcP10 molecules, while ∼23% of the particles classified into an asymmetric structure decorated by a single AcP10 molecule (Fig. S12B; Class 2). The remaining particles lacked AcP10 density. Despite asymmetric AcP10 binding, the reorientation of eIF2Bα and the switch of the eIF2Bδ fulcrum-helix to the A-state conformation was symmetrical on either side of eIF2B^Δ-latch^, a conformational state identical to eIF2B^Δ-latch^ bound by two molecules of AcP10 (Fig. 4D-F, Fig. S12F). This observation further demonstrates that coaxing the A-state on one side of the decamer cooperatively favors the A-state on the opposite side. Thus, despite the loss of the latch-helix, AcP10 pushes the eIF2Bα_2_ dimer uphill from its ground I-state into an A-state conformation, leading to stabilization of enzymatically active eIF2B.

### Pharmacological regulation of eIF2B via the ISRIB pocket depends on latch-helix interactions with eIF2Bα

While NSs and AcP10 access the eIF2-P binding pocket to regulate eIF2B, the small molecule ISRIB functions as an ISR inhibitor by binding a distinct pocket formed by the two-fold symmetry interface between the β- and δ-subunits in eIF2B, ∼50 Å distant from the binding sites of these viral effectors (*4*, *5*, *65*, *66*). ISRIB enhanced the GEF activity of eIF2B^Δ-latch^ three-fold, although it did not restore the activity to eIF2B^WT^ levels (Fig. 5A), prompting us to ask how ISRIB-binding allosterically relates to the latch-helix.

**Fig 5.**
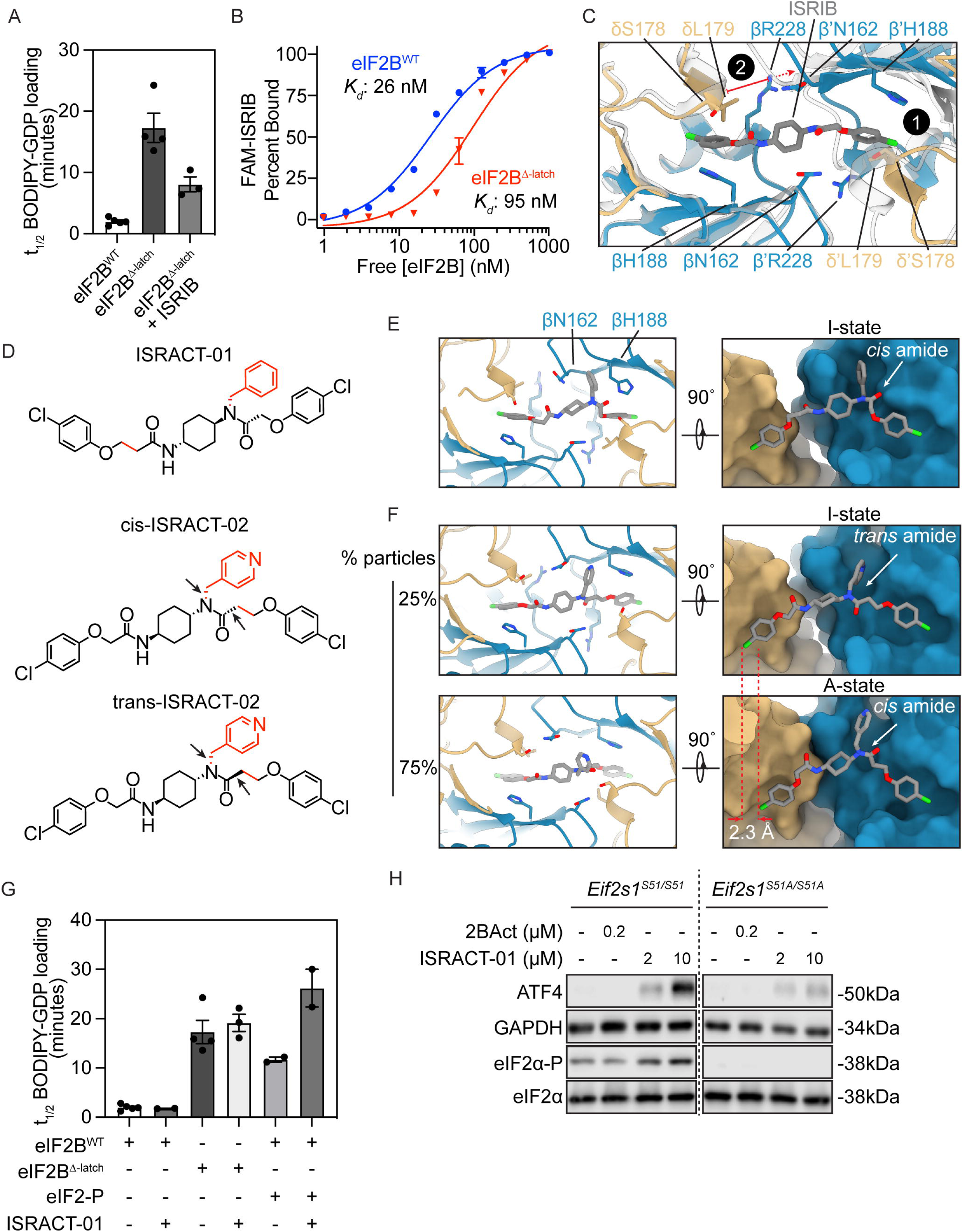
The eIF2B pharmacophore is allosterically modulated by latch-helix ordering. (A) Guanine nucleotide exchange factor (GEF) activity as measured by BODIPY-GDP loading, using 10 nM eIF2B^WT^ or eIF2B^Δ-latch^ (with and without 1 µM ISRIB pre-incubation) as the enzyme and 100 nM eIF2 as the substrate. (B) Fluorescence polarization based binding assay to measure binding of fluorescein conjugated ISRIB (FAM-ISRIB) to eIF2B^WT^ and eIF2B^Δ-latch^. (C) Close-up view of the ISRIB binding pocket of eIF2B^Δ-latch^ (colored) overlaid with a model of ISRIB in complex with eIF2B^WT^ (PDB: 7L7G) in gray. Key residues are shown as sticks and labeled by subunit color with (‘) indicating residues from the opposite tetramer. The observed widening of the pocket is indicated in red. (1) and (2) indicate major structural differences targeted for rational design of eIF2B inhibitors. (D) Chemical structures of compounds ISRACT-01 and ISRACT-02 with the modifications relative to ISRIB shown in red. Cis/trans isomerization of ISRACT-02 around the amide bond are indicated by arrows. (E) Zoom- in view of the ISRACT/ISRIB binding pocket from cryo-EM structures of eIF2B^Δ-latch^ bound to ISRACT-01 I-state shown as cartoons (left) and an alternate viewing direction as surface view (right). (F) Structures of ISRACT-02 in complex with eIF2B^Δ-latch^ in the I-state (top) and A-state (bottom). Percent of particles per conformation are indicated to the left. Alternate surface views on the right demonstrate the change in amide isomerization and end-to-end length of ISRACT-02. (G) GEF activity of eIF2B^WT^ and eIF2B^Δ-latch^ upon treatment with 10 µM of ISRACT-01. eIF2B^WT^ inhibition was also tested upon treatment with 50 nM eIF2-P, or 50 nM eIF2-P + 10 µM ISRACT-01. (H) Immunoblots of mouse embryonic fibroblast lysates of *Eif2fs1^S51/S51^*or *Eif2fs1^S51A/S51A^* cells treated with the indicated concentrations of ISRACT-01 or the ISRIB analogue 2BAct for 3 hours. Panels were cropped from the same gel. Representative blots from n = 4 experiments are shown.

To this end, we first measured the binding affinities of fluorescein-labeled ISRIB (FAM-ISRIB) to both eIF2B^WT^ and eIF2B^Δ-latch^. FAM-ISRIB bound to eIF2B^Δ-latch^ with an approximate four-fold affinity decrease as compared to eIF2B^WT^ (eIF2B^WT^ *K_d_* = 26 nM +/- 0.5 nM; eIF2B^Δ-latch^ *K_d_* = 95 nM +/- 4.7 nM) (Fig. 5B). The structural basis for this reduction in affinity becomes apparent when comparing the ISRIB-binding pocket between the ISRIB-bound eIF2B^WT^ structure (PDB: 7L7G; *grey*) and the eIF2B^Δ-latch^ structure (*colored*) (Fig. 5C). Specifically, the length of the binding pocket increases by more than 4 Å, displacing the hydrophobic pockets that receive the two terminal halogenated benzene rings of ISRIB (*76*). βH188 and δL179 from one half of eIF2B also move apart from the other half, disfavoring the centered binding of ISRIB orthogonal to the symmetry axis. The I-state conformation of the ISRIB-binding pocket is further stabilized by the lever-like rotation of the adjacent βH160 from its trans-interface zippering interaction with β’R228 (‘prime’ indicates an amino acid in the opposing tetramer across the symmetry interface) (Fig. S2C, Movie S2), facilitating opening of the clefts between eIF2Bβ and eIF2Bδ’ from the opposite tetramer.

The lengthening of the ISRIB pocket observed in eIF2B^Δ-latch^ suggested that the corresponding lengthening of ISRIB might stabilize the I-state and pharmacologically activate the ISR. Following this idea, we designed and synthesized ISRIB derivatives that conformed to the longer pocket. To this end, we introduced two key modifications to the glycolamide groups within ISRIB: (i) addition of a methylene group between the amide and ether moieties; and (ii) addition of an aromatic group to the amide nitrogen (based upon observations first reported by A. Zyryanova and D. Ron) (Fig. S13A). We rationalized that modification (i) would increase the overall length of the molecule and allow the terminal aryls to contact their cognate binding sites in the extended pocket, while modification (ii) could potentially interact within the cleft that forms between eIF2Bβ and eIF2Bδ’ (Fig. 5C). We designed two molecules following this scheme, named ISRACT-01 and ISRACT-02 (Fig. 5D). The primary difference between the two compounds is the addition of the methylene extension on the opposing (ISRACT-01) or same (ISRACT-02) side as the aromatic amide adduct.

To determine how these compounds fit within the I-state pocket, we determined cryo-EM structures of eIF2B^Δ-latch^ bound to both compounds (Fig. S14A-F). Our reconstruction of the ISRACT-01-bound eIF2B^Δ-latch^ complex allowed unambiguous modelling of the compound binding pose. Despite stabilizing the Δ-latch I-state conformation as expected, the branching benzyl group that we introduced on ISRACT-01 occupied a shallow surface groove in the space between residues βN162 and βH188 rather than the β/δ’ cleft as we had expected (Fig. 5E).

The consensus structure of eIF2B^Δ-latch^ in complex with ISRACT-02 similarly showed prominent density for the compound but required 3D classification to resolve heterogeneous states. We found that ∼25% of the particles sorted into an I-state decamer in which ISRACT-02 was bound in the same manner as ISRACT-01. By contrast, reconstruction of the remaining ∼75% of the particles revealed ISRACT-02 bound, surprisingly, to an A-state decamer with the aromatic pyridine facing the solvent rather than inserting into the β/δ’ cleft or occupying the pocket between βN162 and βH188 (Fig. 5F).

Closer inspection revealed that in the I-state particles, the ISRACT-02 tertiary amide was in the *trans* conformation, while in the unexpected A-state it was in *cis*. In contrast to secondary amides where *trans* is energetically favored, in tertiary amides both *trans* and *cis* states are expected to have similar prevalence in the free ligand and will interconvert in solution. The remarkable dual activity of ISRACT-02 appears to be a result of the *trans*-amide conformation end-to-end length being ∼2.3 Å longer, which better fits the I-state pocket (Fig. 5F, right). Conversely, from our observations, the *cis*-amide conformation of ISRACT-01 binds eIF2B^Δ-latch^ in the I-state, in which the *trans*-amide conformation would be unable to bind.

These structures were corroborated by measurements of the binding affinities of the FAM-derivatized ISRACT compounds toward eIF2B^WT^ or eIF2B^Δ-latch^ (Fig. S14I): FAM-ISRACT-01 bound eIF2B^WT^ very weakly (*K_d_* = not measurable with confidence), while it bound eIF2B^Δ-latch^ with high nanomolar affinity (*K_d_* = 557 nM +/- 97 nM). As expected, FAM-ISRACT-02 bound eIF2B^WT^ (*K_d_* = 174 nM +/- 32.2 nM) with a two-fold higher affinity compared to eIF2B^Δ-latch^ (*K_d_* = 385 nM +/- 7.8 nM). Together with the structures, these data indicate that ISRACT-01 represents a *bona fide* <u>ISR ACT</u>ivator, while ISRACT-02 can stabilize either the A- or I-state of eIF2B.

To characterize ISRACT-01 further, we next measured its effect on the GEF activity of either eIF2B^WT^ or eIF2B^Δ-latch^ *in vitro*. At saturating concentrations, ISRACT-01 did not have an effect when incubated alone with either the active eIF2B^WT^ or the inactive eIF2B^Δ-latch^ complex (Fig. 5G; conditions 2 and 4). However, ISRACT-01 notably potentiated the effect of limiting amounts of eIF2-P to decelerate nucleotide exchange by eIF2B^WT^ (Fig. 5G; condition 6). These results underscore the notion that ISRACT-01 binding to eIF2B is facilitated by I-state stabilization, leading to a synergistic inhibition in the presence of both eIF2-P and ISRACT-01.

When tested in cells, we found that treatment with ISRACT-01 caused an increase in the expression of ATF4 (Fig. 5H; *Eif2s1^S51/S51^* panel). Furthermore, ISRACT-01 protected cells from apoptosis activated by tunicamycin, an N-linked glycosylation inhibitor that causes ER stress and activation of the eIF2 kinase PERK (Fig. S15A). In contrast, eIF2B activation with ISRIB accelerated tunicamycin-induced cell death, as reported previously (*49*) (Fig. S15A). Polysome profiling of cells treated with ISRACT-01 showed an increase in the 60S and 80S ribosome subunit fractions accompanied by a concomitant decrease in the heavier polysome fractions, phenocopying protein synthesis defects observed during the ISR (*49*) (Fig. S15B,C). In line with the *in vitro* GEF assays described above, ISRACT-01 treatments of cells deficient for eIF2α phosphorylation (*Eif2s1^S51A/S51A^*) (*77*) did not stimulate strong ATF4 accumulation (Fig. 5H; *Eif2s1^S51A/S51A^* panel). Together, these observations indicate that ISRACT-01 binding alone is unable to displace the latch-helix. It is, however, sufficient to stabilize the I-state in the presence of limiting amounts of cellular eIF2-P, thus enhancing a cytoprotective ISR during tunicamycin-induced ER stress.

## Discussion

### The allosteric communication axes in eIF2B

The regulatory allosteric signaling axes described in this work explain how the antagonistic binding of substrate (eIF2) and inhibitor (eIF2-P) is communicated over the 40 Å distance along eIF2B’s central interface. Four conformational elements aligned along two identical, symmetry-related axes operate in concert: (i) the latch-helix docking to its binding pocket in eIF2Bα, (ii) the rotation of the fulcrum-helix in eIF2Bδ, (iii) the βH160 tetramer-tetramer interface zipper, and (iv) the substrate eIF2 binding pocket (Fig. 6). We establish that discrete conformational changes in these four elements, synchronized both within- and across-axes, define the active-to-inactive A- to I-state transition in eIF2B (Movie S4). Two convergently evolved viral effectors exert their effect by locking eIF2Bα, and with it the entire eIF2B complex, into the A-state.

**Fig 6.**
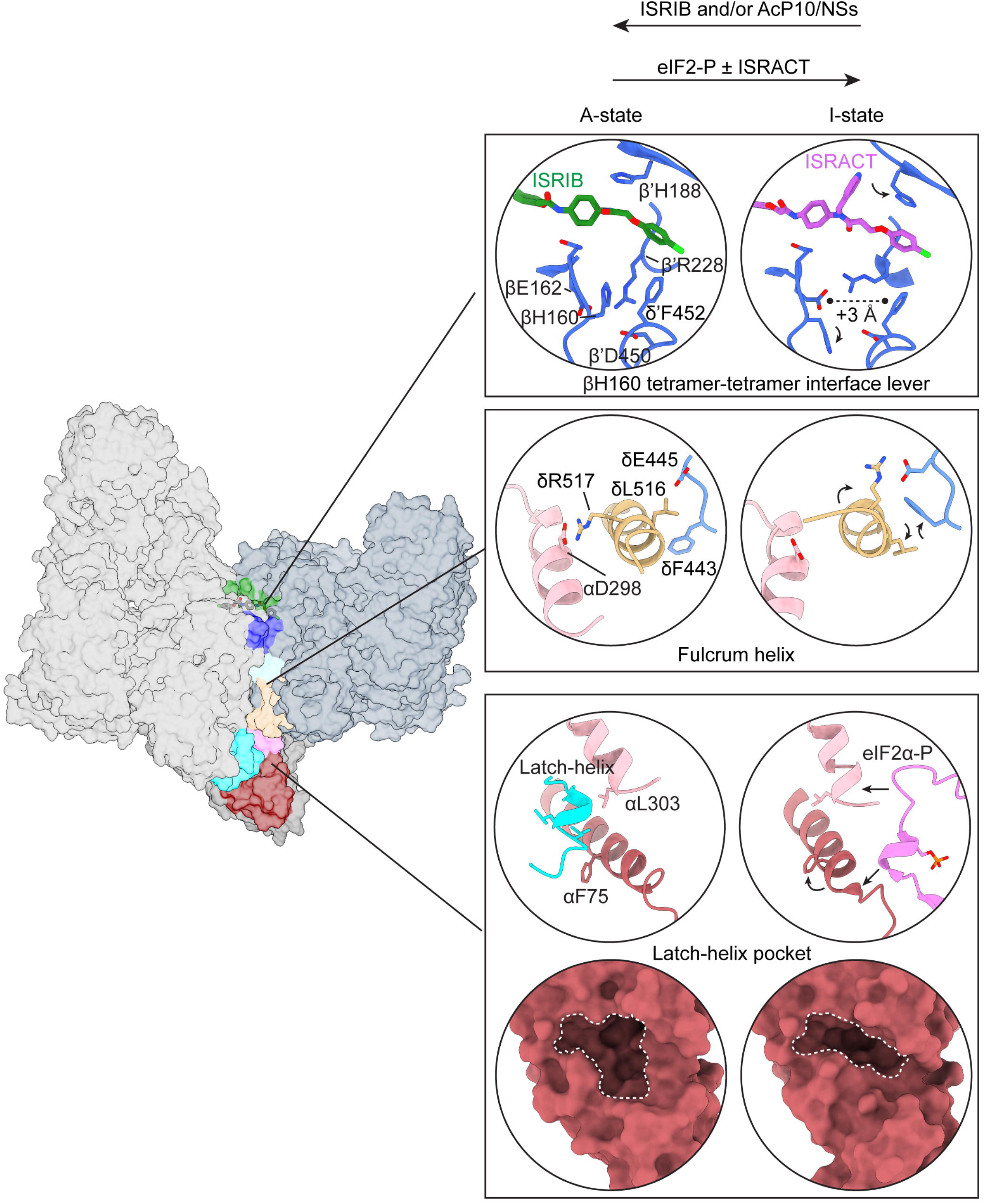
Latch-helix ordering propagates along an allosteric axis that links eIF2B’s substrate, inhibitor, and pharmacophore sites. Model of eIF2B A- to I-state transition encompassing shifts in the conformational equilibrium within the α, β, and δ core that are induced by eIF2-P, small molecules, and effector proteins to regulate eIF2B activity. Arrows depict the conformational changes imparted during I-state stabilization.

Comparing the parsed structure classes of unliganded eIF2B by cryo-EM 3D variability analysis, we demonstrate that apo-eIF2B exists in an equilibrium of A- and I-states, suggesting a relatively flat energy landscape between the two states. In the absence of an inhibitor, engagement of the eIF2Bβ latch-helix with the eIF2Bα latch-helix pocket biases the conformational equilibrium toward the A-state, which the viral effectors AcP10 and NSs further stabilize. By contrast, eIF2-P binding to eIF2B skews the equilibrium towards the I-state by stabilizing a remodeled latch-helix pocket. This interaction reorients eIF2Bα to drive the series of events along the allosteric axes that results in a widening of the substrate binding pocket to disfavor eIF2 binding. In doing so, eIF2-P binding stabilizes eIF2Bα in its intrinsic ground I-state conformation, as observed in crystal structures of eIF2Bα homodimers (Fig. S5C). Experimentally, the I-state can be further stabilized by a rationally designed ISR activator ISRACT-01 that, due to its increased length over the clinical ISR inhibitors based on ISRIB, buttresses the I-state-induced cleft between tetramer-tetramer interfaces.

Two mirrored allosteric axes connect the symmetry interface of the eIF2B complex. This observation expands the model of allosteric regulation proposed by Monod, Wyman, and Changeux (*78*) to include state-transitions of complexes formed by non-identical subunits. The experiments with the viral effector AcP10 provide first evidence that activation or inhibition of eIF2B on one side synchronizes the symmetry-related other side of the complex (Fig. S12F). We pose that such asymmetric communication is likely mimicked by eIF2B’s interactions with its substrate, eIF2, whose binding positions eIF2α as a tether between the two tetramers and hence stabilizes the A-state.

### Evolutionary considerations of latch-helix mediated regulation

Based on sequence and structural homology, the α, β, and δ subunits of eIF2B are evolutionarily related to archaeal aIF2B proteins (*79*). While eIF2Bδ has a large, flexible N-terminal expansion, only eIF2Bβ evolved the latch-helix containing insertion between helices α4 and α5 (Fig. S5A, S5D). Furthermore, analysis of eIF2Bβ sequences across eukaryotes reveals remarkable conservation of the latch-helix with hydrophobic side chains that likely fit snugly into the latch-helix pocket (Fig. S5B, S5D). Thus, it is plausible that a latch-helix based regulatory node in eIF2B co-evolved with its role as a GEF for eIF2 in eukaryotes. Notably, the amino acids defining the latch-helix binding pocket in eIF2Bα are likewise conserved (Fig. S5E) and thus eIF2Bα emerges from this and other studies (*73*) as a determinant of the allosteric modulation effected by metabolites, viral effectors, and eIF2-P.

While the function of the closest related aIF2Bs have not been elucidated, two families of proteins that are sequentially and structurally related to a/eIF2B are sugar phosphate isomerases (*79–81*), and human eIF2Bα retains the capability to bind sugar phosphates M6P and F6P *in vitro* (*73*). We show here that eIF2Bα dynamically closes its latch-helix binding pocket in response to eIF2-P, inducing the I-state. The consensus structures for M6P-bound eIF2Bα dimers and F6P-bound eIF2B decamers likewise adopt I-state conformations. Indeed, in support of our findings, the latch-helix is unresolved in the F6P-bound decamer, yet paradoxically, both M6P and F6P stimulate eIF2B activity *in vitro*—an observation that remains unexplained. In addition to eIF2Bα, eIF2Bβ and δ are also evolutionarily related to sugar phosphate binding proteins and may likewise retain metabolite-binding properties that conceivably regulate eIF2B function.

Surprisingly, the viral protein AcP10 is agnostic to the presence of the latch-helix in eIF2B (see Figs. 3-4). Through a series of intricate, precise, and evolutionarily convergent interactions that rely on aromatic amino acid “fingers”, both AcP10 and NSs coerce the α-subunit of eIF2B to reorient to the A-state conformation (Figs. 3D; 4D-E, S10). Viral protein use of aromatic fingers emerges as a compelling evolutionary means to force productive interaction surfaces, as often observed in high-affinity interactions, such as those mediated between antigens and antibodies (*82*, *83*). Using this trick to good effect, AcP10/NSs target eIF2B, activate the enzyme, and suppress the ISR.

### Considerations of small molecule ISR activators

While an increasing body of literature has focused on activating eIF2B for therapeutic benefit with small molecules (*49*, *54*, *63*, *64*, *84*), the characterization of the eIF2B I-state allowed us and others to design eIF2B inhibitors, ISRACTs, that target the same pocket as the ISR inhibitor ISRIB. ISRACT-01, in particular, is a valuable tool to study the ISR in cells by allowing specific ISR activation directly through eIF2B. This mode of action distinguishes ISRACT-01 from other compounds that induce eIF2 kinase-mediated increases in eIF2-P. Since kinases generally phosphorylate a spectrum of substrate proteins, it is unlikely that ISR kinases only phosphorylate eIF2α (*85*, *86*). Rather, ISR kinases likely change the cell’s phospho-proteome in yet uncharacterized ways and hence alter the molecular environment in which the ISR ultimately exerts its effects. Medicinally, further development of ISRACTs may serve as alternatives to ISR-activating phosphatase inhibitors such as salubrinal (*87*) which has shown promise in models of Charcot-Marie-Tooth disease (*88*) and sickle-cell anemia (*89*), and the GCN2 kinase activator halofuginone that protects from diabetes in a model of diet induced obesity (*90*).

We note that the modifications introduced to the ISRIB backbone to generate ISRACT impose some pharmacokinetic penalties. The binding pose of the added benzyl group restricts the movement of residue βH188 on eIF2B, which is generally observed as a mixture of rotamers, thereby potentially imposing an entropic penalty. Since the discovery of ISRIB, the molecule has been optimized into a clinical candidate through structure-activity relationship studies (*54*, *76*). Given their shared backbone, the principles of improving ISRIB’s pharmacological properties should be applicable to ISRACT as well. Furthermore, our increased structural understanding of eIF2B and its regulation via its allosteric signaling axes opens new ways to explore drugging alternative pockets, such as the latch-helix binding pocket and/or the eIF2- P/metabolite/viral effector binding pockets of the alpha subunit, for therapeutic benefit as we learn with increasing precision how to modulate the ISR across biological scales.

## Acknowledgements

This work was supported by Altos Labs, Inc. We thank Steve McKerrall and Deniz Eismann for helpful conversations, and members of the Altos Labs Institute of Science for comments on the manuscript. We also thank the Altos Labs cryo-EM hub for data collection and analysis, and Josh Lee and She-Rene Chen for support and review of this manuscript.

## Funding

### Author contributions

Conceptualization: TIC, AD, UD, AS, NJ, FZ, FJMvK, JJC, PW, AF

Methodology: UD, AS, TIC, DJL, PFE, CPA, JJC, AF, PW

Investigation: UD, AS, AD, JC, MV, MB, PFE, LH, DJL, YL, LCR, KS, HT, MLV, CPA, TIC

Resources: RJdG, FJMvK

Writing - Original Draft: UD, AS, PFE, TIC, PW, AF

Writing - Review & Editing: All authors

Visualization: UD, AS, PFE, TIC, PW, AF

Supervision: NT, CPA, JN, DAA, MCM, JJC, PW, AF

### Data and materials availability

Cryo-EM density maps and associated models will be publicly available through the wwPDB with accession codes: PDB: 9Y3P, 9Y3U, 9Y3Q, 9Y3T, 9Y3V, 9Y3W, 9Y4B, 9Y5S, 9Y5R, 9Y5T, 9Y5U; EMD-72462, 72467, 72463, 72466, 72468, 72499, 72477, 72521, 72520, 72522, 72523. All other raw data are available upon request through data transfer agreements.

## Supplementary Materials

### Materials and Methods

#### Protein expression and purification

##### eIF2Bβγδε tetramers

Plasmids containing *E. coli* codon-optimized dual ORFs of eIF2Bβ&δ (His-tagged WT: pAD025, His-Strep tagged WT: pPE-011, His-tagged Δ-latch: pAD028) and eIF2Bγ&ε (pAD024) were transformed into OneShot BL21 Star (DE3) chemically competent cells (Invitrogen). Following 1 h recovery in LB media, the entire transformation mixture was used to inoculate 50 mL of LB media containing chloramphenicol and kanamycin and grown overnight at 37 °C. 6 x 1 L of LB media (+chloramphenicol and kanamycin) were inoculated with 12 mL each of the overnight culture and grown at 37 °C until the culture reached an OD_600_ of 0.8. Expression was then induced by addition of IPTG to a final concentration of 0.8 mM and continued for 20 hours at 16 °C. Cells were harvested by centrifugation (10 minutes at 4000 x g) and stored at −80 °C or used immediately for purification. Pellets were resuspended with 4 mL/g wet pellet of cold lysis buffer (40 mM HEPES KOH pH 7.5, 250 mM KCl, 1 mM TCEP, 5 mM MgCl_2_, 15 mM Imidazole pH 8, and 1 tablet/50 mL of cOmplete EDTA-free protease inhibitor cocktail (Roche)). *E. coli* were lysed by three consecutive cycles through an EmulsiFlex-C3 (Avestin) homogenizer then cleared by centrifugation at 18,000 x g for 60 minutes at 4 °C. All subsequent steps were carried out using an AKTA Pure FPLC (Cytiva) at 4 °C. The clarified lysate was loaded onto a 5 mL HisTrap HP column equilibrated in HisTrap buffer A (20 mM HEPES KOH pH 7.4, 200 mM KCl, 5 mM MgCl_2_, 15 mM Imidazole, 1 mM TCEP), followed by washing with 5 CV of the buffer. Sample was then eluted from the column over a 0-100% linear gradient into HisTrap buffer B (20 mM HEPES KOH pH 7.4, 200 mM KCl, 5 mM MgCl_2_, 300 mM Imidazole, 1 mM TCEP). Peak fractions containing eIF2Bβγδε were pooled and applied to a MonoQ 10/100 GL column equilibrated in IEX buffer A (20 mM HEPES KOH pH 7.4, 200 mM KCl, 5 mM MgCl_2_, 1 mM TCEP) and then washed with 5 CV of the buffer. Bound proteins were eluted using a linear 0-100% gradient of IEX buffer B (20 mM HEPES KOH pH 7.4, 500 mM KCl, 5 mM MgCl_2_, 1 mM TCEP) over 10 CV. Peak fractions containing eIF2Bβγδε were pooled, concentrated, and loaded onto a Superdex 200 Increase 10/300 column equilibrated with SEC buffer (20 mM HEPES KOH pH 7.4, 200 mM KCl, 5 mM MgCl_2_, 5% glycerol, 1 mM TCEP). The final purified sample was concentrated using a 100k MWCO spin concentrator (Millipore), flash frozen in liquid nitrogen, and stored at −80 C until use. A typical yield was 3 mg of >95% pure tetramer from 6 L of *E. coli* culture.

##### eIF2Bα dimers

eIF2Bα dimers were initially purified as described previously (*65*). The following protocol was employed to increase yield and purity of the final protein. A modified pRSF Duet plasmid containing *E. coli* codon-optimized (His8-MBP-Thrombin cut site) tagged eIF2Bα ORF (pPFE ALTOS-041) was transformed into OneShot BL21 Star (DE3) chemically competent cells (Invitrogen) and grown on LB-Agar containing Kanamycin. A single colony was used to inoculate a 100 mL overnight culture. 20 mL of the overnight culture was used to inoculate 3 x 1 L of TB media (+Kanamycin) and the culture was grown at 37 °C until the culture reached an OD_600_ of 0.8. Expression was then induced by addition of IPTG to a final concentration of 0.8 mM and continued for 20 hours at 17 °C. Cells were harvested by centrifugation (17 minutes at 4000 x g) and stored at −80 °C or used immediately for purification. Pellets were resuspended with 100 mL of cold lysis buffer (40 mM Tris-HCl pH 7.5, 500 mM NaCl, 5% glycerol, 1 mM TCEP, 1 mM PMSF, and 1 tablet/50 mL of cOmplete EDTA-free protease inhibitor cocktail (Roche)).

*E. coli* were lysed by four consecutive cycles through an EmulsiFlex-C3 (Avestin) homogenizer then cleared by centrifugation at 23,000 x g for 60 minutes at 4 °C. The clarified lysate was incubated with 10 mL Cobalt IMAC resin (Takara) (equilibrated in lysis buffer) and incubated for 1 hour with gentle rotation to allow batch binding. The resin was applied to a gravity column and washed with 25 CV of lysis buffer containing 12.5 mM imidazole, and bound protein was eluted using 7 CV of lysis buffer containing 125 mM imidazole. The elution fraction was next incubated with 10 mL of equilibrated Amylose resin (NEB) for 30 minutes with gentle rotation. The resin was washed with 5 CV Amylose wash buffer (40 mM Tris-HCl pH 7.5, 300 mM NaCl, 5% glycerol) then eluted with 5 CV of Amylose wash buffer supplemented with 50 mM maltose. The eluted proteins were concentrated to 1 mL and purified further by size exclusion chromatography using a HiLoad Superdex200 Increase 16/60 column equilibrated in SEC buffer (20 mM HEPES-KOH pH 8, 200 mM KCl, 5 mM MgCl_2_, 0.5 mM TCEP) on an AKTA Pure instrument (Cytiva). The peak fractions containing His-MBP-eIF2Bα were pooled and the His-MBP tag was then cleaved using thrombin for 96 hours at 4 °C. The resultant mixture was purified by size exclusion chromatography using a HiLoad Superdex 75 16/60 column in SEC buffer. The peak fractions containing cleaved protein were pooled and passed over a Cobalt IMAC-Amylose mixed slurry to remove residual His-MBP fusions. The flow-through was collected, concentrated, flash frozen in liquid nitrogen and stored at −80 °C until use. A typical yield was ∼25 mg of >99% pure protein from 3 L of *E. coli* culture.

##### AcP10

*E. coli* BL21 (DE3) cultures expressing His-MBP-AcP10 or corresponding mutant plasmids were first cultured overnight at 37 °C. These overnight cultures were subcultured (1:100) in LB medium, supplemented with the appropriate antibiotics, to OD600 = 0.6. Protein expression was then induced with 1 mM IPTG and the culture was grown at 16 °C overnight. The next day, the bacteria were centrifuged at 6,000 r.p.m. for 5 min at 4 °C, followed by incubation of the pellets in bacterial lysis buffer (50 mM Tris-HCl pH 8, 300 mM NaCl, 5 mM Imidazole, 0.2 mM TCEP, 0.5 mM PMSF, 5% glycerol and complete protease inhibitor) for 60 min on ice. The mixtures were then lysed by three passes through an EmulsiFlex C3 high pressure homogenizer (Avestin). The lysate was then cleared by centrifugation at 20,000 x g for 45 min at 4 °C. The supernatant was applied onto a gravity column packed with TALON IMAC resin at 4 °C to enable batch binding for 1 hour at 4 °C. The resin was washed with five column volumes of wash buffer (50 mM Tris-HCl pH 8, 300 mM NaCl, 20 mM Imidazole, 0.1 mM TCEP, 0.1 mM PMSF, 5% glycerol and cOmplete protease inhibitor). His-MBP-AcP10 was eluted with elution buffer (50 mM Tris-HCl pH 8.0, 300 mM NaCl, 200 mM glutathione, 0.1 mM TCEP, and 5% glycerol) and concentrated at 4 °C using Amicon Ultra centrifugal filters with repeated mixing to prevent aggregate formation. The purified proteins were analyzed by SDS–PAGE and Coomassie brilliant blue staining. The protein concentrations were calculated by measuring absorbance at 280 nm and applying the Beer–Lambert law (A = εlc; ε = 82990). In certain cases, proteins were further purified by gel filtration chromatography using a Superdex 200 column attached to an AKTA Pure FPLC (Cytiva) and running buffer containing 50 mM Tris-HCl pH 8.0, 300 mM NaCl and protease inhibitor cocktail. Elution fractions containing protein were analysed by their A280 spectra, SDS–PAGE and Coomassie brilliant blue staining.

##### eIF2 and eIF2-P

Heretotrimeric eIF2 was generated by co-expression of full length eIF2α, N-term FLAG-Tev tagged eIF2β (2-333, K79R, K80R, K81R, K83R, K84R), eIF2γ (91–472), CDC123 in Expi293F cells. Equimolar ratios of codon-optimized plasmids (0.75 µg of each plasmid per mL of cell culture) were diluted in Opti-MEM media (Gibco) to a final concentration of 18 µg/mL. The diluted plasmids were then combined with an equivalent volume of PEI at 0.12 mg/mL in Opti-MEM. The mixture was added dropwise to cells at a density of ∼3×10^6^ cells/mL to transfect the plasmids. The cells were grown for 20 hours on an orbital shaker at 37 °C, 8% CO_2_ at a speed of 125 rpm before addition of Expi Enhancer to the culture. Cell pellets were harvested by centrifugation 72 h after transfection. Pellets were resuspended with 5 mL/g lysis buffer (20 mM HEPES KOH pH 7.4, 300 mM KCl, 1 mM MgCl_2_, 0.001 mM Triton X-100, 1 tablet/50 mL protease inhibitor cocktail) and lysed by sonication (3 s on, 5 s off, 200 W for 100 cycles).

Lysates were clarified by two consecutive centrifugations for 30 mins at 20300 x g, 4 °C. The supernatant was incubated with anti-FLAG M2 affinity gel (Sigma) and bound protein was eluted with 100 µg/ml of FLAG peptide. 1mM GDP was added to the eluate and incubated for one hour. For phosphorylation, GST-PKR was added to the solution at 1:20 (PKR:eIF2, w/w) and ATP was added to a final concentration of 1 mM. The protein treated with/without PKR kinase was further purified by size exclusion chromatography using a Superdex 200 Increase 10/300 GL (Cytiva) column and concentrated by ultra-filtration (Amicon, MWCO 30 kD). The final protein was analyzed using Phos-tag SDS-PAGE (Fujifilm) to determine the phosphorylation state, then flash frozen in liquid nitrogen and stored at −80°C until use.

##### eIF2α(NTD)-P

The coding sequence of the human eIF2α subunit N-terminal domain (residues 2-187) with an N-term 6xHis-Tev cut site tag was codon optimized, synthesized, and cloned into a pET28(a) vector. The plasmid was transformed into chemically competent OneShot Bl21 Star (DE3) *E. coli* and a single colony was picked to start an overnight culture. Following overnight growth at 37 °C, the culture was used to inoculate 2 L (1:250) of ZYP-5052 autoinduction media which was then grown for 20 hours at 20 °C. Cell pellets were harvested by centrifugation for 10 mins at 4000 x g, washed once with PBS and stored at −80 °C until use. Cells were resuspended in lysis buffer (20 mM Tris-HCl pH 7.5, 500 mM NaCl, 20 mM imidazole, 1 mM PMSF, and 1 tablet/50 mL EDTA-free protease inhibitor cocktail) and lysed by sonication using a Branson 450 sonifier (pulses of 10 s on, 10 s off for 20 minutes at 60% power). Lysates were cleared by centrifugation at 18000 x g for 30 min and passed through a 0.45 µm bottle-top filter before loading onto an equilibrated HisTrap FF column (Cytiva) using an AKTA Pure FPLC instrument. The column was then washed with 10 CV of eIF2α wash buffer (20 mM Tris-HCl pH 7.5, 500 mM NaCl) supplemented with 20 mM imidazole and a further 5 CV of buffer with 62.5 mM imidazole. Bound proteins were eluted with elution buffer (wash buffer + 250 mM imidazole). The elution fraction was supplemented with 5 mM MgCl_2_ and 2.5 mM ATP (Sigma) then phosphorylated by addition of 5 µg recombinant GST-PKR (Thermo) for 2 hours at 37 C with gentle stirring. The entire reaction was syringe filtered (0.22 µm filter), concentrated to 1 mL, and purified further by size exclusion chromatography using a Superdex 200 Increase pg 16/600 column. Phosphorylation and purity of the final sample was evaluated by SDS PAGE using a Phos-tag gel (Fujifilm). A typical yield was 2 mg of protein per liter of *E. coli* culture.

##### eIF2α-P

eIF2α-P was purified as previously described (*36*). N-terminally 6x-His-tagged human eIF2α was transformed into One-Shot BL21 Star (DE3) chemically competent *E. coli* cells and eIF2α expression was induced with 1 mM IPTG. The culture was grown overnight at 16°C. Cell pellets were then lysed using the EmulsiFlex-C3 in a buffer containing 100 mM HEPES-KOH, pH 7.5, 300 mM KCl, 2 mM dithiothreitol (DTT), 5 mM MgCl2, 5 mM imidazole, 10% glycerol, 0.1% IGEPAL CA-630, and protease inhibitor cocktail. The lysate was clarified at 20,000 rpm for 45 min at 4°C. Clarified lysate was loaded onto a AKTA Pure 5 ml His-Trap FF Crude column, washed in a buffer containing 20 mM HEPES-KOH, pH 7.5, 100 mM KCl, 5% glycerol, 1 mM DTT, 5 mM MgCl2, 0.1% IGEPAL CA-630, and 20 mM imidazole, and eluted with 75 ml linear gradient of 20 mM – 500 mM imidazole. The eIF2α-containing fractions were collected and applied to a MonoQ HR 10/100 GL column equilibrated in a buffer containing 20 mM HEPES-KOH pH 7.5, 100 mM KCl, 1 mM DTT, 5% glycerol, and 5 mM MgCl2 for anion exchange. The column was washed in the same buffer, and the protein was eluted with a gradient of 100 mM to 1 M KCl. eIF2α-containing fractions were concentrated with an Amicon Ultra-15 concentrator with a 30 kDa molecular mass cutoff. eIF2α was phosphorylated by incubating the concentrate with 1 mM ATP and 1 mg of PKR-GST enzyme (amino acids 252-551) per milligram of eIF2α overnight at 4°C. eIF2α-P was then purified on a Superdex 75 10/300 GL column equilibrated in a buffer containing 20 mM HEPES-KOH pH 7.5, 100 mM KCl, 1 mM DTT, 5 mM MgCl2, and 5% glycerol, and concentrated again using an Amicon Ultra-15 concentrator with a 10 kDa molecular mass cutoff and the protein was flash frozen in liquid nitrogen and stored at −80°C. Samples were run on a 12.5% PhosTag gel to verify phosphorylation status.

#### Cryo-EM sample preparation and data collection

eIF2B decamers were assembled on ice for 15 minutes by combining eIF2Bβγδε tetramers (WT or Δ-latch) with eIF2Bα_2_ at final concentrations of 12 µM and 9 µM, respectively. Where applicable, ligands were added after complex assembly at the following molar ratios to decamer: 3:1 eIF2α(NTD)-P, 2:1 MBP-AcP10, 15:1 ISRACT-01 (0.5% DMSO), 15:1 ISRACT-02 (0.5% DMSO). Following a 30-minute incubation on ice with ligands, samples were vitrified using a Vitrobot Mark IV plunge freezer (Thermo) set to 4 °C and 100% humidity. Quantifoil R 1.2/1.3, 300 mesh 25 nm gold grids were glow discharged using an EasiGlow glow discharger (Pelco) for 30 s at 15 mA immediately before use. 3.5 µL of sample was applied to a grid and excess liquid was removed using dual-side blotting with a blot force of −5 for 1.5 s. The grids were immediately plunged into liquid ethane then transferred to and stored in liquid nitrogen until clipping and imaging. The apo-eIF2B^WT^, eIF2B^WT^-AcP10, eIF2B^Δ-latch^-ISRACT-01, and eIF2B^Δ-latch^-ISRACT-02 datasets were collected using a Glacios TEM (Thermo) operating at 200 keV while apo-eIF2B^Δ-latch^ and eIF2B^Δ-latch^ -AcP10 datasets were collected using a Titan Krios G4 TEM (Thermo) operating at 300 keV. Both microscopes are equipped with Falcon 4i detectors and SelectrisX energy filters with a slit width of 10 eV. The microscope and data collection parameters for each dataset are provided in Supplemental table 1.

#### Cryo-EM image analysis

The specific details of image analysis are displayed in supplemental figures S1 (eIF2B^WT^), S4 (eIF2B^Δ-latch^), S7 (eIF2B^WT^-AcP10), S12 (eIF2B^Δ-latch^-AcP10), S14 (eIF2B^Δ-latch^-ISRACT-01, eIF2B^D-latch^-ISRACT-02) and CryoSPARC workflow files are available at https://github.com/udalwadi/latch-helix. For all datasets, patch-based global and local motion correction, and patch CTF estimation were performed within CryoSPARC live (Structura Biotechnology) using default settings. All subsequent image analysis was carried out using CryoSPARC v4.2 or newer (*91*). For the apo eIF2B^WT^ dataset, particles were initially picked with the blob picker using a particle diameter range of 160-220 Å and minimum separation distance of 0.7 diameters. Particles were extracted with a box size of 512 pixels and Fourier cropped to 400 pixels, then subjected to 2D classification into 50 classes with default settings. 2D classes resembling eIF2B were selected, then used for template-based particle picking for all datasets in this study, with a particle diameter of 220 Å. For all datasets, template-picked particles were extracted and subjected to one round of 2D classification to remove non-particles then subjected to *ab initio* reconstruction and heterogenous refinement into multiple classes. Particles from the best-refining classes were subjected to homogenous refinement with global and local CTF-refinement to generate the consensus map for each dataset. The refined particles were then subjected to 3D classification and 3DVA, where applicable, and each 3D class was refined further by homogenous refinement and used for subsequent model building.

#### Model building, refinement, and analysis

The initial apo eIF2B^WT^ A-state structural model was generated starting by global relaxation of 7L7G into the map using ISOLDE (*92*) within UCSF ChimeraX (*93*) by running the simulation with simple distance restraints enabled. Each chain and residue were then manually refined to fit the experimental map and the residues comprising the latch-helix were manually built and fit. This model was then used as a starting template to build into all subsequent experimental maps containing eIF2B following the same procedure. Models of ISRACT-01 and ISRACT-02 were imported to ChimeraX by their SMILES codes, parameterized using ISOLDE and fit manually into observed cryo-EM density. An AlphaFold prediction of the human uridine cytidine kinase UCK1 that shares homology with AcP10 (*71*) was used as a template to build the structure into the experimental density maps containing AcP10. A final refinement of all models was carried out against half-maps using Servalcat (*94*) and the resulting statistics are reported in **Supplemental table 1**. Experimental structures are deposited to the wwPDB with accession codes PDB: 9Y3P, 9Y3U, 9Y3Q, 9Y3T, 9Y3V, 9Y3W, 9Y4B, 9Y5S, 9Y5R, 9Y5T, 9Y5U; EMD-72462, 72467, 72463, 72466, 72468, 72499, 72477, 72521, 72520, 72522, 72523. Measurements of all reported structural features were done using the ‘distance’ command in ChimeraX between alpha-carbons of the appropriate residues. All figures and movies including cryo-EM maps and models were generated using ChimeraX.

#### Targeted reanalysis of hydrogen-deuterium exchange coupled to mass spectrometry on eIF2B to measure latch-helix dynamics

Hydrogen-deuterium exchange coupled to mass spectrometry (HDX-MS) datasets generated by Lawrence *et al.* (*67*) were mined to obtain measurements of relative latch-helix dynamics when eIF2B is in the apo-state compared to when it is stabilized in either the A-state (apo-eIF2B versus eIF2B:NSs complex) or the I-state (apo-eIF2B versus eIF2B:eIF2-P complex). The difference in deuteration levels (ΔD) of all detected peptides (n = 5) corresponding to different regions within amino acids 108 - 141 of eIF2Bβ at each exchange time point were plotted. Positive values represent peptides with less deuteration (more solvent protection) in eIF2B complexes with NSs/eIF2-P than in apo-eIF2B in isolation. Negative values represent peptides with more deuteration (less solvent protection) in eIF2B complexes with NSs/eIF2-P than in apo-eIF2B in isolation.

#### In vitro guanine nucleotide exchange assay

The nucleotide exchange activity of eIF2B was measured as either the loading of BODIPY-GDP to eIF2 (‘loading’) or subsequent exchange of GDP for BODIPY-GDP (‘unloading’) and the measurement is indicated in each relevant figure. Purified eIF2 (100 nM) was incubated with 100 nM BODIPY-FL-GDP (Thermo Fisher Scientific) in assay buffer (100 mM HEPES-KOH (pH 7.5), 100 mM KCl, 5 mM MgCl_2_, 1 mM TCEP and 1 mg/mL bovine serum albumin) to a volume of 500 µl in a black-walled 1.5-ml tube. eIF2B decamers were assembled for >45 minutes at room temperature with a molar ratio of eIF2Bβδγε tetramers to eIF2Bα dimers of 1.5 : 2.0. 10x concentrated GEF mixes were assembled with 100 mM eIF2B and a 10x concentration of any ligand as indicated in individual figures. 18 µL per well of the eIF2 - BODIPY-GDP - BSA mix was added to 384-square-well, black-walled, clear-bottom polystyrene assay plates (Corning #3766) and a baseline fluorescence intensity was measured for 30 seconds using a Clariostar Plus (BMG LabTech) plate reader (λ_Ex_ 497 nm, λ_Em_ 525 nm). 2 µl of the 10× GEF mixes were then spiked into the 384-well plate wells with a multichannel pipette to achieve the final concentration of proteins and drugs as indicated in each figure. Fluorescence intensity was recorded every 10 s for 60 min at room temperature. For unloading assays, wells were spiked with 1 mM GDP after completion of the loading reaction and fluorescence intensity measurement was resumed. Data collected were fit to a first-order exponential association (‘loading’) or first-order exponential decay (‘unloading’) with Prism (GraphPad) and the reaction half-time after curve fitting are reported as t_1/2_.

#### FAM-ISRIB fluorescence polarization affinity assay

All fluorescence polarization measurements were performed in 20 μl reactions with 2.5 nM FAM-ISRIB (Praxis Bioresearch) in FP buffer (20 mM HEPES-KOH (pH 7.5), 100 mM KCl, 5 mM MgCl_2_ and 1 mM TCEP) and measured in 384-well, non-stick black plates (Corning, 3820) using a ClarioStar Plus (BMG LabTech) plate reader at room temperature. Before the reaction setup, eIF2B decamers were assembled in FP buffer at 10 µM using eIF2Bβγδε and eIF2Bα_2_ in a 2:1.5 molar ratio for at least 1 h at 4 °C. FAM-ISRIB, FAM-ISRACT-01, or FAM-ISRACT-02 were always first diluted to 2.5 μM in 100% NMP before dilution to 50 nM in 2% NMP and then added to the reaction (0.1% NMP final). eIF2B decamers were serially diluted two-fold 10 times starting from 1.111 µM and the 18 µL of the dilutions were transferred to the 384-well assay plate. 2 µL of the 50 nM fluorescent compound was then added to each well. The plate was incubated at room temperature for 30 min before measurement of parallel and perpendicular intensities (λ_Ex_ 482 nm, λ_Em_ 530 nm).

#### Analytical ultracentrifugation

Analytical ultracentrifugation sedimentation velocity experiments were performed using the Optima AUC (Beckman Coulter). Samples were loaded into cells in AUC buffer (20 mM HEPES-KOH (pH 7.5), 150 mM KCl, 1 mM TCEP and 5 mM MgCl_2_) at 1 µM for eIF2Bβδγε tetramers and 0.75 µM for eIF2Bα dimers. A buffer-only reference control was also loaded. Samples were then centrifuged in an AN 50 Ti rotor at 40,000 r.p.m. at 20 °C, and 280-nm absorbance was monitored for 100 scans every 100 s. Subsequent data analysis was conducted with Sedfit using a non-model-based continuous c(*S*) distribution and graphs were generated using Prism (GraphPad).

#### In vitro immunoprecipitation assays

Assays were performed with a modified protocol from a previous report (*65*). For MBP-tag pulldowns, purified His-MBP-AcP10, His-MBP-AcP10 mutants, eIF2B(αβδγε)_2_, eIF2Bβδγε, eIF2B(αβ^Δ-latch^δγε)_2_, eIF2Bβ^Δ-latch^δγε and eIF2Bα_2_ were incubated (with gentle rotation at 4°C) with Amylose resin in Assay Buffer (20 mM HEPES-KOH, pH 7.5, 150 mM KCl, 5 mM MgCl_2_, 1 mM TCEP, 1 mg/ml bovine serum albumin (BSA), 5 mM CaCl_2_). After 1 h, the resin was pelleted by benchtop centrifugation and the supernatant was removed. Resin was washed 3 times with 500 µl of ice cold Assay Buffer before resin was resuspended in Elution Buffer (Assay Buffer with 5 mM EDTA and 200mM Maltose) and incubated with gentle rocking for 30 minutes. The resin was then pelleted and the eluate was incubated with Laemmeli buffer and boiled. Samples were analyzed by Western Blotting.

For eIF2B pulldowns, purified eIF2Bβδγε tetramers with a Strepavidin-eIF2Bβ were used. Purified His-MBP-AcP10, His-MBP-AcP10 mutants and preassembled eIF2B^WT^ decamers were first incubated (with gentle rotation at 4°C) in Assay Buffer (20 mM HEPES-KOH pH 7.5, 150 mM KCl, 5 mM MgCl_2_, 1 mM TCEP, 1 mg/ml bovine serum albumin (BSA), 5 mM CaCl_2_). After 1 h, eIF2ɑ-P was added to the samples where indicated for another hour at 4°C. Streptavidin-Sepharose resin was then added to the mixtures and incubated for a further hour with gentle rotation. The resin was pelleted by benchtop centrifugation and the supernatant was removed. Resin was washed 3 times with 500 µl of ice cold Assay Buffer before the resin was resuspended in Elution Buffer (Assay Buffer with 5 mM EDTA and 2.5 mM Desthiobiotin) and incubated with gentle rocking for 30 minutes. The resin was then pelleted and the eluate was incubated with Laemmeli buffer and boiled. Samples were analyzed by Western Blotting.

#### Crosslinking mass spectrometry of AcP10-eIF2B complexes

##### PhoX crosslinking and protein digestion

His-MBP-AcP10 was incubated with the eIF2B decamer at a molar ratio of 5:1 for 1 hour at 25°C in a final volume of 40 µL. 10% of the input was removed for SDS-Page analysis, and Disuccinimidyl Phenyl Phosphonic Acid (DSPP or PhoX) was added to the remaining solution at a final concentration of 1mM. The crosslinking reaction was carried out at 25°C for 1 hour before quenching with Tris pH 8.5 to a final concentration of 20 mM. To check cross-linking efficiency, 2.5 μL of each sample (non-crosslinked and crosslinked) was mixed with 2.5 μL of 2x Laemmli loading buffer containing 5 mM TCEP (Bond-Breaker). Samples were then heated for 5 minutes at 95°C. 5 μL of the prepared SDS-PAGE samples were loaded onto a 12-well stain-free 4-15% Criterion gel. The gel was imaged under UV light using a Bio-Rad ChemiDoc system. Successful crosslinking was determined by the upward shift of the AcP10 and eIF2B bands.

Protein samples were reduced with 5 mM TCEP for 30 minutes at 37°C on a ThermoMixer shaking at 1000 rpm. Alkylation was then performed using 15mM 2-chloroacetamide for 30 minutes in the dark at 25°C on a ThermoMixer shaking at 1000 rpm. Excess 2-chloroacetamide was quenched by DTT at a final concentration of 10mM and incubating for 30 minutes at 25°C. For protein precipitation, 300μL of cold acetone was added to each sample and vortexed briefly, followed by overnight precipitation at −30°C. Precipitated protein was centrifuged at 21,130 x g at 4°C for 20 minutes and acetone was removed carefully without disturbing the pellet. Samples were airdried for 15 minutes at 25°C and resuspended in 25 μL of 8M Urea, 0.1M Tris pH 8.5. For digestion, LysC was added at a 1:40 ratio of enzyme:protein sample and incubated at 25°C on a ThermoMixer at 1000rpm for 3 hours. To dilute the urea concentration down to 1M before trypsin digestion, 175μL of 0.1M Tris pH 8.5 was added to each sample. Trypsin was then added at a 1:40 enzyme:protein ratio and digested overnight at 25°C.

##### Peptide C18 Clean-up and IMAC Enrichment

Digested peptides were brought to 1% trifluoroacetic acid (TFA) by adding 22.2 μL of 10% TFA in water. Samples were briefly vortexed, and droplets were collected by centrifugation. 210μL of each sample was then transferred to a LoBind 96-well plate. 200μL of peptide sample was desalted by C18 clean-up performed on an AssayMAP Bravo automated liquid handling system (Agilent Technologies; Santa Clara, Ca). Peptides were eluted from the C18 cartridges with 25μL of 80% acetonitrile (ACN) in 0.1% TFA. Crosslinked peptides were enriched on the AssayMAP Bravo using 5μL Fe(III)-NTA cartridges. The cartridges were primed with 200 μL of 0.1% TFA in ACN and equilibrated with 200 μL of loading buffer (80% ACN/0.1% TFA). Peptides were resuspended in a final volume of 210μL 80% ACN/0.1% TFA and 200μL was loaded onto the cartridge. The cartridges were washed with 50 μL of 80% ACN/0.1% TFA, and the crosslinked peptides were eluted with 25 μL of a 3:1 ratio of H_2_O: NH_4_OH into a 96-well LoBind plate containing 25μL of 10% formic acid in H_2_O to immediately neutralize the pH. The plate was immediately dried using a speedvac and stored at −80°C. The peptides were resuspended in 3% ACN:0.1% formic acid and identified by LC-MS/MS analyses.

##### Database Search

Spectral raw data files were first processed using MSFragger (version 4.0) with a FASTA file combining the H. sapiens and E. coli proteomes (downloaded on April 11, 2024, UP000005640 and UP000000625) including the relevant fusion protein sequences and a list of common contaminant proteins. Cysteine carbamidomethylation (57.02146) was set as fixed modification. Methionine oxidation (15.9949), protein N-term acetylation (42.0106), phosphorylation (79.96633 on S, T, Y), and hydrolysed and Tris-quenched PhoX crosslinker modifications (227.98238 and 331.0457 on K respectively) were set as dynamic modifications. The proteins identified by MSFragger were combined into a FASTA file to generate a searchable library that was subsequently used with the XlinkX node 3.0 of the Proteome Discoverer 3.0 software (Thermo Fisher Scientific) to identify crosslinked peptides using the noncleavable open search algorithm. The analysis was performed using cysteine carbamidomethylation as a fixed modification and methionine oxidation as a variable modification. PhoX (+209.97181 Da) was set as the crosslinker modification on lysine residue while also allowing protein N-terminus linkage with the following parameters: minimum peptide length 5, missed cleavages 2, precursor mass tolerance 10 ppm, fragment mass tolerance 20 ppm, and 5% FDR on CSM and cross-link levels. The crosslinks were visualized using the xMAS plugin in ChimeraX.

#### Cell lines and cell culture

HEK293T (ATCC, CRL-3216), MEFs (gift from R. J. Kaufman, Sanford Burnham Prebys Medical Research Institute) and BJ fibroblasts (ATCC, CRL-2522) were cultured in DMEM-high glutamine medium. DMEM media was supplemented with 10% fetal bovine serum and the cells were maintained at 37 °C and 5% CO2. For the MEF and BJ fibroblast cell cultures, the DMEM medium was further supplemented with penicillin– streptomycin antibiotic, sodium pyruvate and non-essential amino-acid mixtures. The HEK293T cells were seeded in poly-L-lysine–treated plates. The cells were cultured to approximately 75–90% confluency for most experiments. Each cell line was routinely screened for mycoplasma contamination by the Tissue Culture Hub at Altos-BAI. HEK293T and BJ fibroblast cells from ATCC were authenticated by short-tandem-repeat profiling.

#### Transfection

HEK293T cells were transfected with plasmid vectors using the TransIT-2020 transfection reagent for 24 hours. Lipofectamine RNAiMax was used to carry out siRNA treatments in HEK293T cells for 72 hours. For co-expression of eIF2Bβ siRNA and eIF2Bβ siRNA resistant plasmids, HEK293T cells were first transfected in suspension with eIF2Bβ siRNA using Lipofectamine RNAiMax for 24 hours. The next day, the cell culture media was replaced with fresh media containing the eIF2Bβ siRNA resistant plasmid:TransIT-2020 mixture and cells were cultured for a further 48 hours. Expression or knockdown efficiencies were checked in parallel for every experiment by immunoblotting. The eIF2Bβ siRNA sequence is provided in Supplementary Table 3. Plasmids with ORFs expressing AcP10, AcP10 mutants, NSs and siRNA resistant eIF2Bβ^WT^ and eIF2Bβ^Δ-latch^ were sequenced and verified before use. The ORF sequences are provided in Supplementary Table 3.

#### Puromycin pulse–chase assays

Assays were performed as previously reported (*95*). Cells were treated with 2 µg/ml puromycin for 10 min, followed by puromycin wash out with PBS. The cells were chased with complete medium with or without the indicated stressor for 50 min, washed with PBS and lysed with RIPA buffer (1% Triton X-100, 20 mM Tris–HCl pH 8.0, 0.1% SDS, 0.05% sodium deoxycholate, 150 mM NaCl, 10 mM Na3VO4, 40 mM β-glycerophosphate and 10 mM NaF). The lysates were then subjected to immunoblotting.

#### ISR activation assays with sodium arsenite

HEK293T cells were treated with 50 µM of sodium arsenite (LabChem Cat. No.LC229001) dissolved in DMEM media supplemented with 10% FCS for 4 hours to induce the ISR.

#### Cell-extract preparation and immunoblotting

Cells were washed three times with PBS and immediately collected at 4 °C in RIPA lysis buffer containing phosphatase inhibitors (1% Triton X-100, 20 mM Tris–HCl pH 8.0, 0.1% SDS, 0.05% sodium deoxycholate, 150 mM NaCl, 10 mM Na3VO4, 40 mM β-glycerophosphate and 10 mM NaF) and complete protease inhibitors (Roche). The detergent-soluble supernatant fractions were immediately processed for SDS–PAGE and immunoblotting with antibodies. A list of antibodies used in this study are provided in Supplementary Table 4.

#### IncuCyte caspase assay

BJ fibroblasts were seeded in 96-well plates with 100 µL EMEM (ATCC, 30-2003) supplemented with 10% FBS (Omega Scientific, FB-01), 1x GlutaMAX (Thermo Fisher Scientific, 35050061), and 1x MEM non-essential amino acid (Thermo Fisher Scientific, 11140035). The following day, the medium was changed to 80 µL of medium supplemented either with ISRIB (200 nM), ISRACT-01 (3 µM), or the control medium without ISRIB and ISRACT-01. After 4 hours of pre-incubation with ISRIB or ISRACT-01, tunicamycin treatment was started by adding 20 µL of the medium supplemented with tunicamycin (5 µM) and reagents for Incucyte analyses: Incucyte Caspase-3/7 Dye for Apoptosis (Sartorius, 4804, 1:1000 dilution) and Incucyte Nuclight Rapid NIR Dye for Live-Cell Nuclear Labelling (Sartorius, 4440, 1:1000 dilution). The final concentrations of the compounds were 1 µM for tunicamycin, 160 nM for ISRIB, and 2.4 µM for ISRACT-01. Immediately, the 96-well plates were placed in the Incucyte S3 Live-Cell Analysis System (Sartorius), and the cells were recorded with the 10X objective every 2 hours for 5 days (excluding one time point every 24 hours). Using Incucyte Analysis Software (Sartorius), Caspase 3/7 intensity of each well was obtained as the area with the signal above a cutoff for Incucyte Caspase-3/7 Dye divided by the total cell count obtained from the Incucyte Nuclight Rapid NIR Dye.

#### Polysome profiling

Ribosome profiling was performed as we previously described with some modifications (*64*). Briefly, we prepared fresh 12 ml of 10 – 50% (m/v) sucrose gradients [20 mM Tris-HCl (pH 8), 5 mM MgCl_2_, 150 mM KCl] in open-top Polyclear tubes (Seton Scientific, 9/16×3-3/4 inch) with gradient master that comes with the fractionator (Biocomp; we used the long-cap protocol to accommodate the lysate volume). Gradients were kept at 4 °C for at least 30 min before use. WT mouse fibroblasts in 15-cm dishes at ∼80% confluency were treated with vehicle (DMSO), thapsigargin (50nM) or ISRACT-01 (10μM) for 30 min before subject to a wash with cold PBS and lysis in 3x polysome lysis buffer [1x concentrations: 20 mM Tris-HCl (pH 8), 5 mM MgCl_2_, 150 mM KCl, 1 mM DTT, 1% NP-40, 100 μg/ml cycloheximide, and EDTA-free protease inhibitors (Roche Indianapolis, IN)] and centrifuged at 14,000 rpm for 10 min at 4 °C. The supernatant was transferred to a pre-chilled tube. We normalized the input amount by RNA concentration measured by qubit (RNA broad range kit, Fisher Scientific). The supernatant was layered onto sucrose gradient carefully. Gradients were centrifuged in a SW-40Ti rotor at 40,000 rpm at 4 °C for 2 hr and then analyzed by piston gradient fractionator (Biocomp) and the absorbance of the material was monitored and recorded using Triax flow cell. Ribosome profile traces were then plotted using Prism (GraphPad).

#### Chemical synthesis

Visual schematics of chemical synthesis are provided in Fig. S13. Data files of ^1^H NMR, ^13^C NMR, and LC-MS analyses of synthesized products are included as Data S1.

##### ISRACT-01

*Step 1*. Tert-butyl N-[trans-4-[N-benzyl-2-(4-chlorophenoxy)acetamido]cyclohexyl]carbamate

To a stirred solution of tert-butyl N-[trans-4-(benzylamino)cyclohexyl]carbamate (3.00 g, 9.85 mmol, 1.00 equiv) and TEA (2.99 g, 29.6 mmol, 3.00 equiv) in DCM (30 mL) was added 4-chlorophenoxyacetyl chloride (2.22 g, 10.8 mmol, 1.10 equiv) at 0 °C. The reaction was stirred for 2 h at 0 °C. The reaction was quenched with water (40 mL). The resulting mixture was extracted with DCM (2 x 40 mL). The combined organic layers were washed with brine (30 mL), dried over anhydrous Na_2_SO_4_. After filtration, the filtrate was concentrated under reduced pressure to give tert-butyl N-[trans-4-[N-benzyl-2-(4-chlorophenoxy)acetamido]cyclohexyl]carbamate (3.70 g, 79% yield) as a white solid. LCMS (ES, m/z): 473.20, 475.20 [M+H]^+^

*Step 2*. N-benzyl-2-(4-chlorophenoxy)-N-[trans-4-aminocyclohexyl]acetamide hydrochloride

To a stirred solution of tert-butyl N-[trans-4-[N-benzyl-2-(4-chlorophenoxy)acetamido]cyclohexyl]carbamate (2.00 g, 4.23 mmol, 1.00 equiv) in ethyl acetate (10 mL) was added HCl (4 M in 1,4-dioxane) (10.6 mL, 42.3 mmol, 10.0 equiv) at 25 °C. The reaction was stirred for 2 h at 25 °C. The mixture was concentrated and washed with Et_2_O (20 mL) to give N-benzyl-2-(4-chlorophenoxy)-N-[trans-4-aminocyclohexyl]acetamide hydrochloride (1.5 g, 87% yield) as a white solid. LCMS (ES, m/z): 373.20, 375.20 [M-HCl +H]^+^

*Step 3.* 3-(4-Chlorophenoxy)-N-[trans-4-[N-benzyl-2-(4-chlorophenoxy)acetamido]cyclohexyl]propanamide

To a stirred solution of N-benzyl-2-(4-chlorophenoxy)-N-[trans-4-aminocyclohexyl]acetamide hydrochloride (2.00 g, 4.90 mmol, 1.00 equiv) and 3-(4-chlorophenoxy)propanoic acid (1.18 g, 5.88 mmol, 1.20 equiv) in DMF (20 mL) was added DIEA (1.88 g, 14.7 mmol, 3.00 equiv) and HATU (2.42 g, 6.37 mmol, 1.30 equiv) at 25 °C. The reaction was stirred for 2 h at 25 °C. The reaction was purified by reversed flash chromatography (column: C18 column, 120 g, 20-35 µm; mobile phase, A: water (10 mM NH_4_HCO_3_) and B: ACN (0% to 75% gradient in 30 min); detector: 254/220 nm). The collected fraction was lyophilized to give 3-(4-chlorophenoxy)-N- [trans-4-[N-benzyl-2-(4-chlorophenoxy)acetamido]cyclohexyl]propanamide (1.08 g, 36% yield, 98.8% purity) as a white solid.

##### ISRACT-02

*Step 1.* 2-(4-Chlorophenoxy)-N-[trans-4-[(pyridin-4-ylmethyl)amino]cyclohexyl]acetamide

To a stirred solution of 2-(4-chlorophenoxy)-N-[trans-4-aminocyclohexyl]acetamide (440 mg, 1.56 mmol, 1.00 equiv) in DCM (5.0 mL) was added 4-formylpyridine (168 mg, 1.56 mmol, 1.00 equiv) and AcOH (9.4 mg, 0.156 mmol, 0.10 equiv) at 25 °C. The reaction was stirred for 0.5 h at 25 °C. Then STAB (996 mg, 4.68 mmol, 3.00 equiv) was added and the mixture was stirred for 2 h at 25 °C. The reaction was quenched with water (20 mL). The mixture was extracted with DCM (2 x 20 mL). The combined organic layers were washed with brine (30 mL), dried over anhydrous Na_2_SO_4_. After filtration, the filtrate was concentrated under reduced pressure. The crude product was purified by silica gel column, eluting with PE/EA (1:2) to afford 2-(4-chlorophenoxy)-N-[trans-4- [(pyridin-4-ylmethyl)amino]cyclohexyl]acetamide (300 mg, 50% yield) as a white solid. LCMS (ES, m/z):374.15, 376.15 [M+H]^+^

*Step 2*. 3-(4-Chlorophenoxy)-N-(pyridin-4-ylmethyl)-N-[trans-4-[2-(4-chlorophenoxy)acetamido]cyclohexyl]propanamide

To a solution of 2-(4-chlorophenoxy)-N-[trans-4-[(pyridin-4-ylmethyl)amino]cyclohexyl]acetamide (240 mg, 0.642 mmol, 1.00 equiv), 3-(4-chlorophenoxy)propanoic acid (193 mg, 0.963 mmol, 1.50 equiv) in DCM (4.0 mL) was added HOBt (173 mg, 1.28 mmol, 2.00 equiv) and EDCI (185 mg, 0.963 mmol, 1.50 equiv) at 25°C. The reaction was stirred for 2 h at 25 °C and concentrated under reduced pressure. The residue was purified by reversed flash chromatography (column: C18 column, 40 g, 20-35 µm; mobile phase, A: water (10 mM NH_4_HCO_3_) and B: ACN (10% to 70% gradient in 40 min); detector: 254/220 nm). The collected fraction was lyophilized to afford 3-(4-chlorophenoxy)-N-(pyridin-4-ylmethyl)-N-[trans-4-[2-(4-chlorophenoxy)acetamido]cyclohexyl]propanamide (41.0 mg, 11% yield, 97.3% purity) as a white solid.

#### Data Reproducibility

The number of replicates for each experiment are ≥ 2 and statistical parameters for each experiment are listed in the corresponding figure legends. No statistical methods were used to pre-determine sample sizes. Data distribution was assumed to be normal, but this was not formally tested. Experimental design and data collection was randomized. For each experiment with multiple conditions, recombinant protein preparations or cultured cells were from the same batch and allocated at random in test-tubes or culture dishes. The independent variable was controlled by direct manipulation. All other sources of extraneous variability were controlled by randomization or chance. As a collaborative study, each investigator was blinded from experiments and data collection conducted by every other investigator. Data collection and analysis by an individual investigator was however not performed blind to the conditions of the experiments. For the densitometric analysis of western blots, protein bands on membranes were captured on a BioRad Chemidoc Touch or a LI-COR Chemidoc apparatus. The exposure times were varied to obtain appropriate signal intensities of the protein bands. The intensities of each of the bands were quantitated using the Image-J gel analysis tool.

---

**Supplemental Table 1.1:** Cryo-EM data collection, analysis, and refinement statistics of wild-type and Δ-latch eIF2B.

|  | eIF2B<sup>WT</sup> |  | eIF2B<sup>Δ-latch</sup> |  |
| --- | --- | --- | --- | --- |
|  | A-state | I-state | I-state | A-state |
|  | PDB: 9Y3P<br>EMD-72462 | PDB: 9Y3Q<br>EMD-72463 | PDB: 9Y3U<br>EMD-72467 | PDB: 9Y3T<br>EMD-72466 |
| Data collection and processing |  |  |  |  |
| Microscope | Thermo Scientific Glacios<br>TEM |  | Thermo Scientific Krios G4 TEM |  |
| Magnification | 130,000x |  |  |  |
| Voltage (kV) | 200 |  | 300 |  |
| Election exposure<br>(e<sup>-</sup>/Å<sup>2</sup>) | 50 |  |  |  |
| Defocus range<br>(μm) | -0.6 to -1.6 |  |  |  |
| Pixel size (Å) | 0.8936 |  | 0.9286 |  |
| Symmetry<br>imposed | C1 |  |  |  |
| Initial particle<br>images (no.) | 890,578 |  | 907,870 |  |
| Final particle<br>images (no.) | 60,012 | 55,475 | 96,151 | 88,960 |
| Map resolution (Å) | 2.94 | 3.30 | 2.42 | 2.50 |
| FSC threshold | 0.143 |  |  |  |
| Map resolution<br>range (Å) | 2.52-38.2 | 2.50-42.7 | 2.05-31.8 | 2.03-35.7 |
| Refinement |  |  |  |  |
| Initial model used<br>(PDB code) | 7L7G | 7L7G | 7L7G | 7L7G |
| Map sharpening B<br>factor (Å<sup>2</sup>) | 56.3 | 56.8 | 47.2 | 48.4 |

| Model composition |  |  |  |  |
| --- | --- | --- | --- | --- |
| Nonhydrogen atoms | 28946 | 28563 | 28169 | 28492 |
| Protein residues | 3709 | 3709 | 3613 | 3657 |
| Ligands | 15 | 15 | 4 | 8 |
| Root mean square deviations |  |  |  |  |
| Bond lengths (Å) | 0.008 | 0.0073 | 0.0078 | 0.0073 |
| Bond angles (°) | 1.715 | 1.609 | 1.625 | 1.6 |
| Validation |  |  |  |  |
| MolProbity score | 1.01 | 1.13 | 1.13 | 1.31 |
| Clashscore | 0.81 | 1.27 | 1.58 | 1.74 |
| Poor rotamers (%) | 1.27 | 0.94 | 1.18 | 1.32 |
| Ramachandran |  |  |  |  |
| Favoured (%) | 96.95 | 95.86 | 97.03 | 95.89 |
| Allowed (%) | 3 | 4 | 3 | 4 |
| Disallowed (%) | 0.05 | 0.06 | 0.03 | 0.19 |

**Supplemental Table 1.2:** Cryo-EM data collection, analysis, and refinement of wild-type and Δ-latch eIF2B bound to ligands.

|  |  |  |  |  |  |  |
| --- | --- | --- | --- | --- | --- | --- |
|  | eIF2B<sup>WT</sup>-<br>AcP10 | eIF2B<sup>Δ-latch</sup> – AcP10 |  | eIF2B<sup>Δ-latch</sup><br>– ISRACT-01 | eIF2B<sup>Δ-latch</sup> – ISRACT-02 |  |
|  | 2 AcP10 | 2 AcP10 | 1 AcP10 | I-state | A-state | I-state |
|  | PDB:<br>9Y3V<br>EMD-72468 | PDB: 9Y4B<br>EMD-72477 | PDB:<br>9Y4W<br>EMD-72499 | PDB: 9Y5S<br>EMD-72521 | PDB: 9Y5T<br>EMD-72522 | PDB: 9Y5U<br>EMD-72523 |
| Data collection and processing |  |  |  |  |  |  |
| Microscope | Thermo<br>Scientific<br>Glacios<br>TEM | Thermo Scientific Krios<br>G4 TEM |  | Thermo Scientific Glacios TEM |  |  |
| Magnification | 130,000x |  |  |  |  |  |
| Voltage (kV) | 200 | 300 |  | 200 |  |  |
| Election<br>exposure (e<sup>-</sup><br>/Å<sup>2</sup>) | 50 |  |  |  |  |  |
| Defocus<br>range (μm) | -0.6 to -1.6 |  |  |  |  |  |
| Pixel size (Å) | 0.8936 | 0.9286 |  | 0.8936 |  |  |
| Symmetry<br>imposed | C1 |  |  |  |  |  |
| Initial particle<br>images (no.) | 1,805,332 | 867,729 |  | 2,714,171 | 2,376,924 |  |
| Final particle<br>images (no.) | 220,193 | 171,498 | 112,494 | 157,984 | 418,565 | 325,317 |
| Map<br>resolution (Å) | 2.90 | 2.13 | 2.42 | 3.10 | 2.78 | 2.90 |
| FSC<br>threshold | 0.143 |  |  |  |  |  |
| Map<br>resolution<br>range (Å) | 1.93-34.8 | 2.08-27.8 | 2.05-31.7 | 1.92-39.6 | 1.93-30.6 | 1.93-35.0 |
| Refinement |  |  |  |  |  |  |
| Initial models used (PDB code) | 7L7G, AF-Q9HA47-F1 |  |  | 7L7G |  |  |
| Map sharpening B factor (Å<sup>2</sup>) | 100.5 | 37.4 | 43.6 | 82.3 | 87.4 | 85.2 |
| **Model composition** |  |  |  |  |  |  |
| Nonhydrogen atoms | 32247 | 32179 | 30512 | 28748 | 28652 | 28658 |
| Protein residues | 4126 | 4076 | 3869 | 3650 | 3631 | 3634 |
| Ligands | 3 | 13 | 10 | 11 | 7 | 5 |
| **Root mean square deviations** |  |  |  |  |  |  |
| Bond lengths (Å) | 0.0095 | 0.0081 | 0.0076 | 0.0081 | 0.0092 | 0.008 |
| Bond angles (°) | 1.796 | 1.655 | 1.607 | 1.709 | 1.840 | 1.71 |
| **Validation** |  |  |  |  |  |  |
| MolProbity score | 1.13 | 0.95 | 0.95 | 1.30 | 1.19 | 1.28 |
| Clashscore | 1.15 | 1.19 | 1.04 | 1.62 | 1.39 | 1.58 |
| Poor rotamers (%) | 1.48 | 0.76 | 0 | 1.5 | 1.51 | 1.50 |
| **Ramachandran** |  |  |  |  |  |  |
| Favoured (%) | 97.02 | 97.4 | 97.2 | 96.16 | 96.9 | 96.3 |
| Allowed (%) | 3 | 3 | 3 | 4 | 3 | 4 |
| Disallowed (%) | 0.05 | 0 | 0.03 | 0.08 | 0.03 | 0.08 |

**Supplemental Table 2:** Apo eIF2B^WT^ 3D classification model A- and I-state analysis.

| Class | Particles | Substrate interface |  | Fulcrum-helix |  | Latch-helix |  |
| --- | --- | --- | --- | --- | --- | --- | --- |
|  |  | Side 1 | Side 2 | Side 1 | Side 2 | Side 1 | Side 2 |
| 1 | 54965 | 32.8 | 33.2 | A-state | A-state | Present | Present |
| 2 | 57555 | 33.4 | 33.5 | A-state | A-state | Absent | Present |
| 3 | 54977 | 33.4 | 33.1 | A-state | A-state | Present | Present |
| 4 | 55271 | N/A | N/A | N/A | N/A | N/A | N/A |
| 5 | 57477 | 32.7 | 33.5 | A-state | A-state | Present | Absent |
| 6 | 57885 | 33.1 | 33.0 | A-state | A-state | Present | Present |
| 7 | 59301 | N/A | N/A | N/A | N/A | N/A | N/A |
| 8 | 55468 | 35.7 | 35.4 | I-state/<br>A-state | I-state/<br>A-state | Absent | Absent |
| 9 | 59983 | 33.3 | 33.4 | A-state | A-state | Present | Present |
| 10 | 56533 | 33.9 | 33.6 | A-state | A-state | Present | Present |

**Supplemental Table 3:**
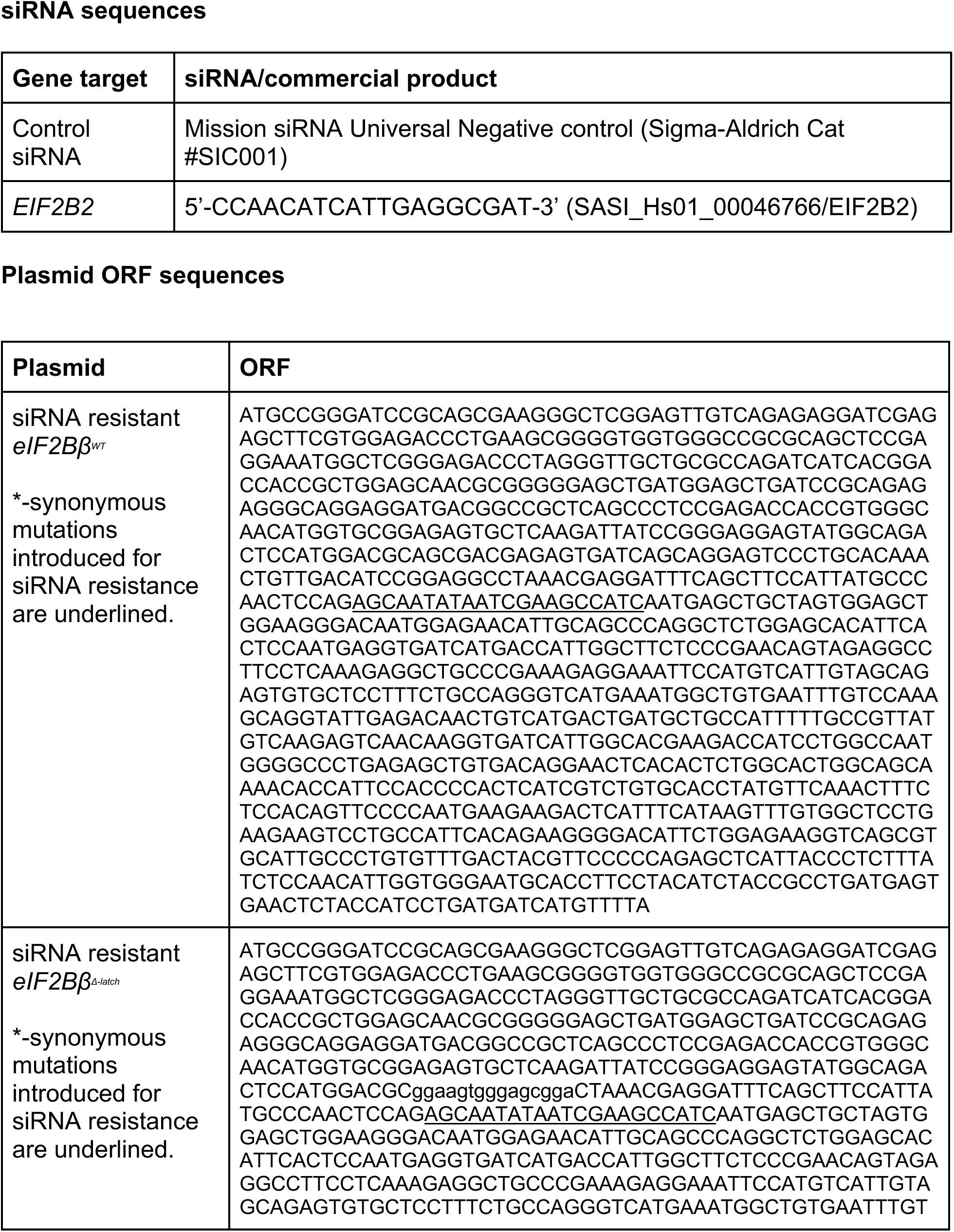
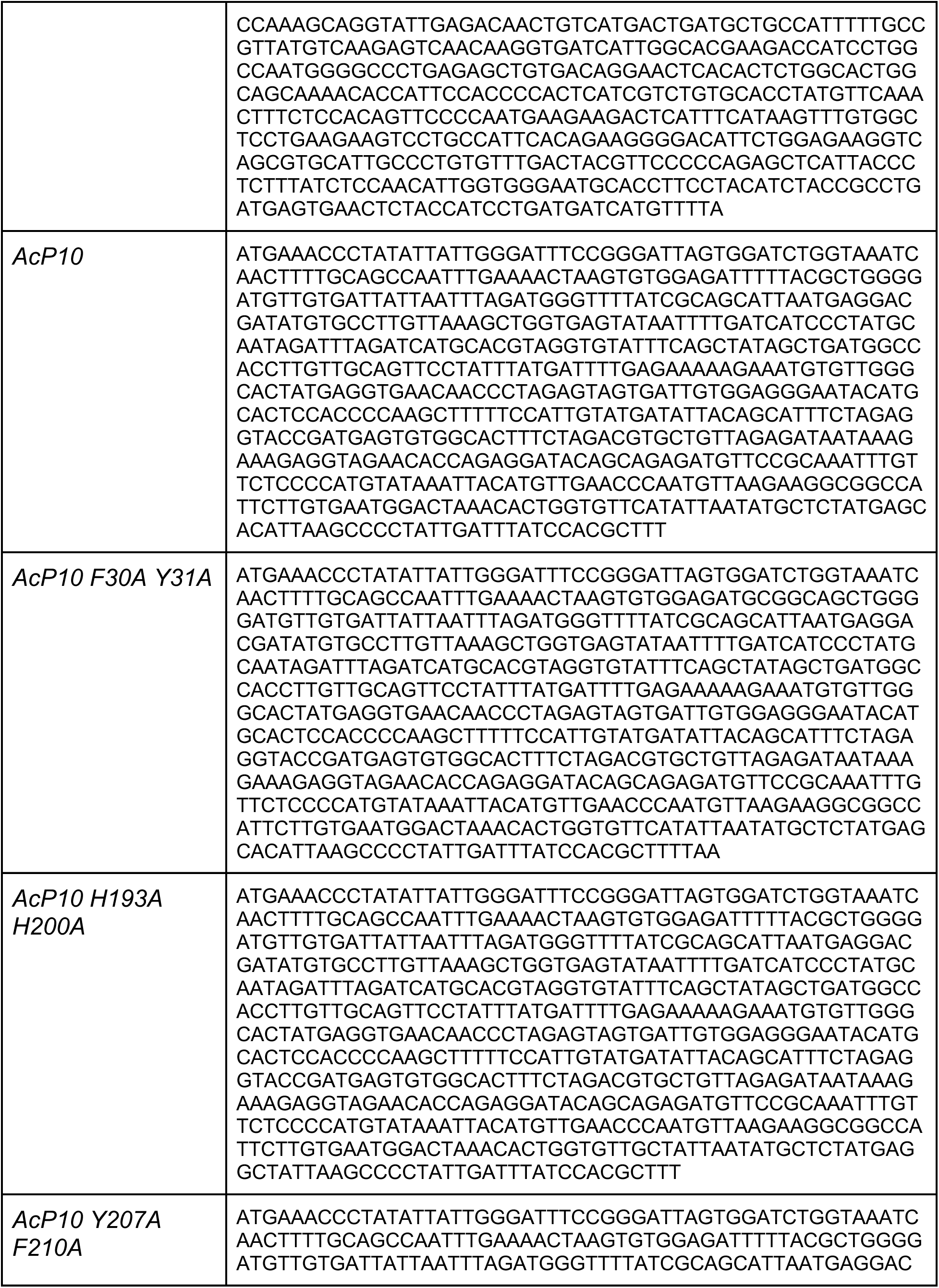
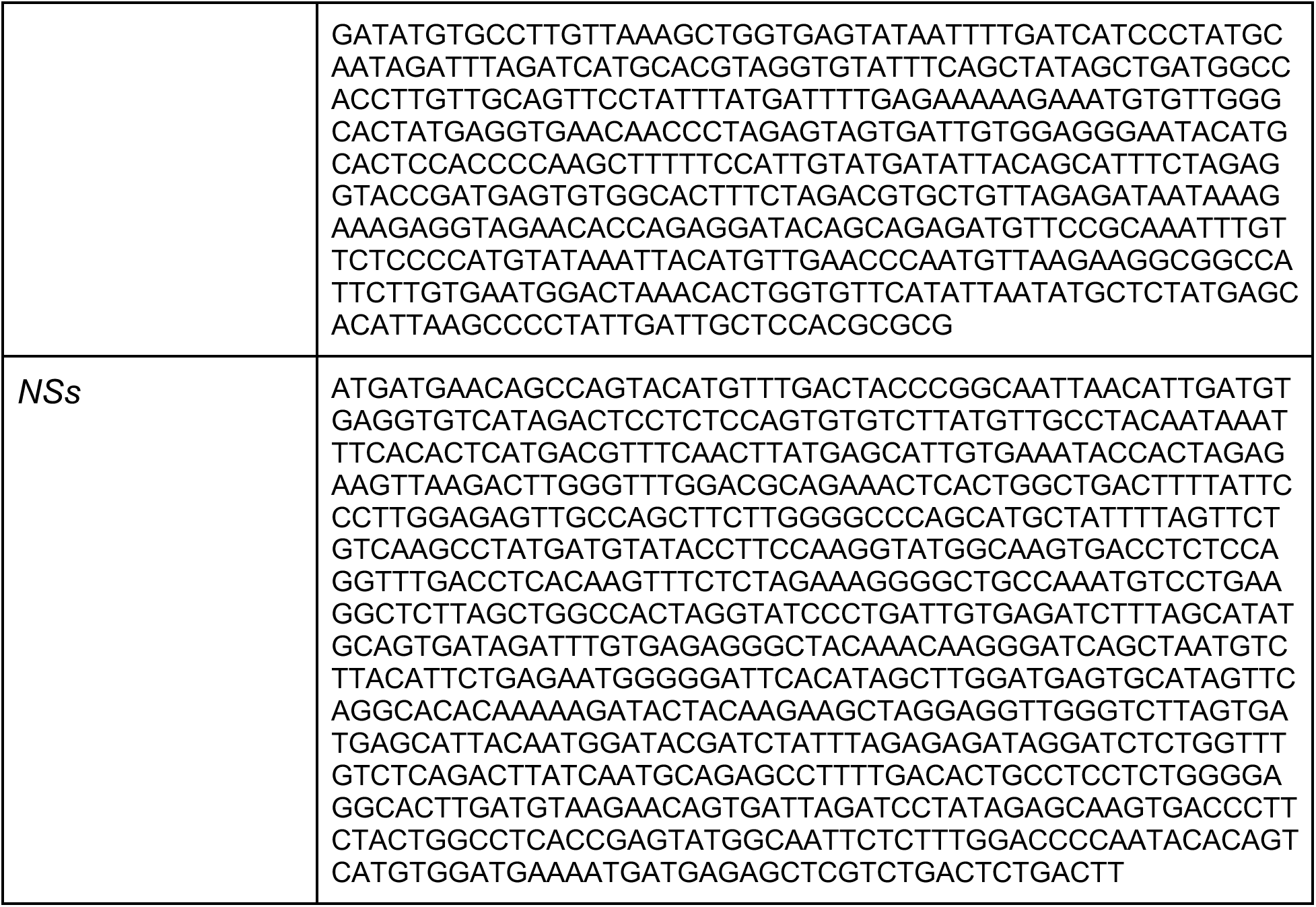
siRNA and plasmid ORF sequences siRNA sequences.

**Supplemental Table 4:**
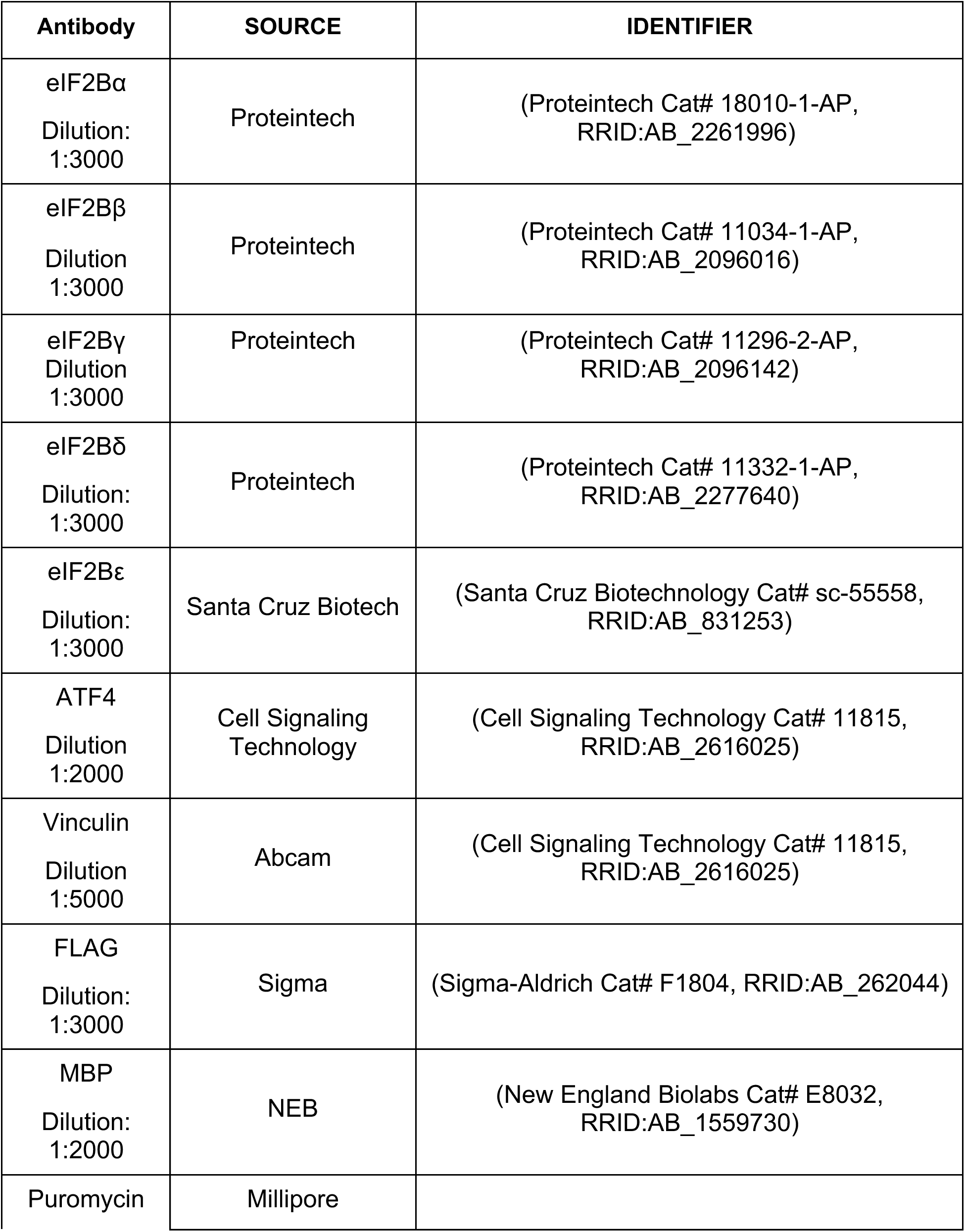
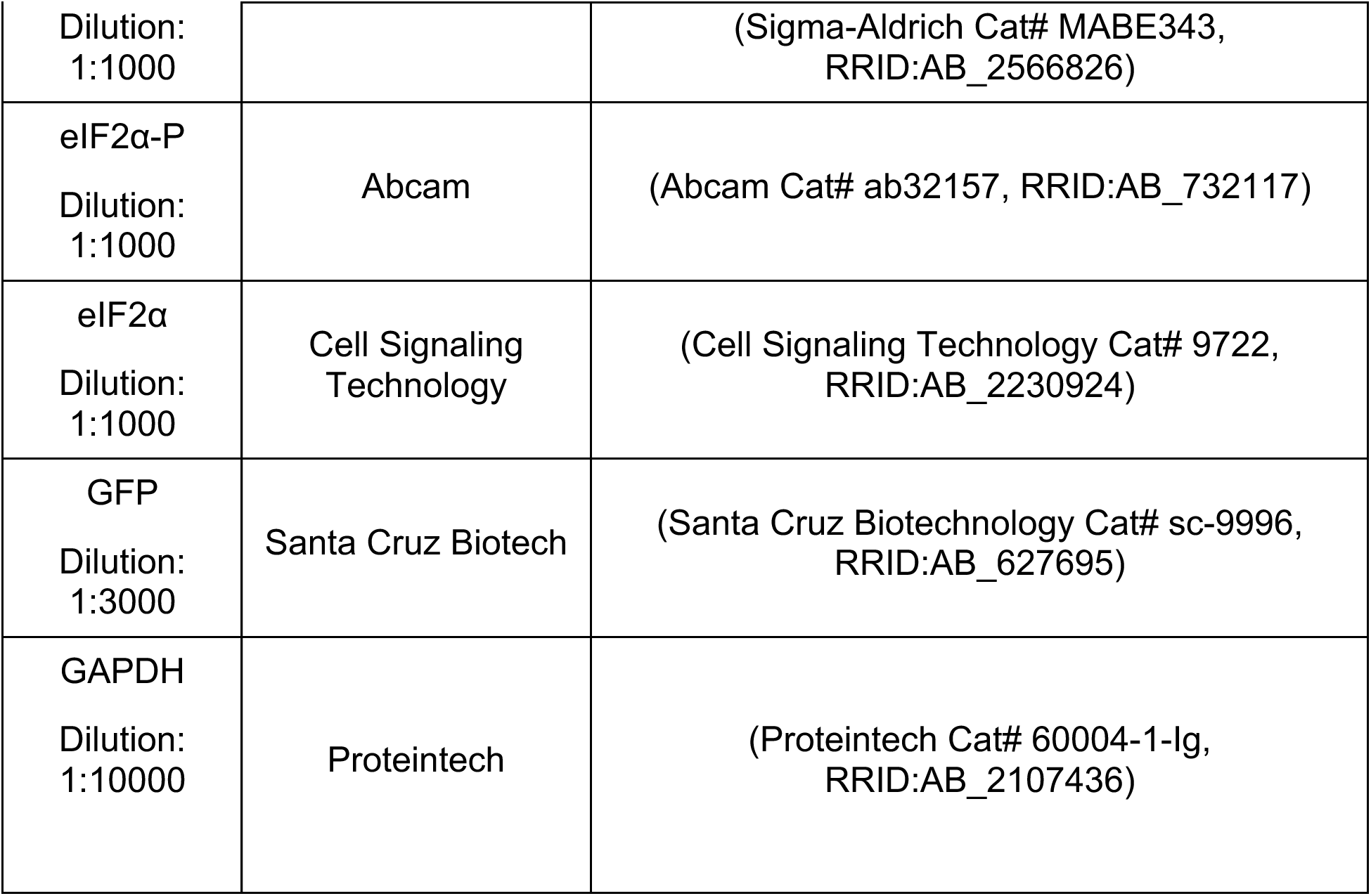
Immunoblotting antibodies used in this study.

