## Supplementary Figures for "Allosteric Disordering of eIF2B Regulates the Integrated Stress Response"

Figure S1

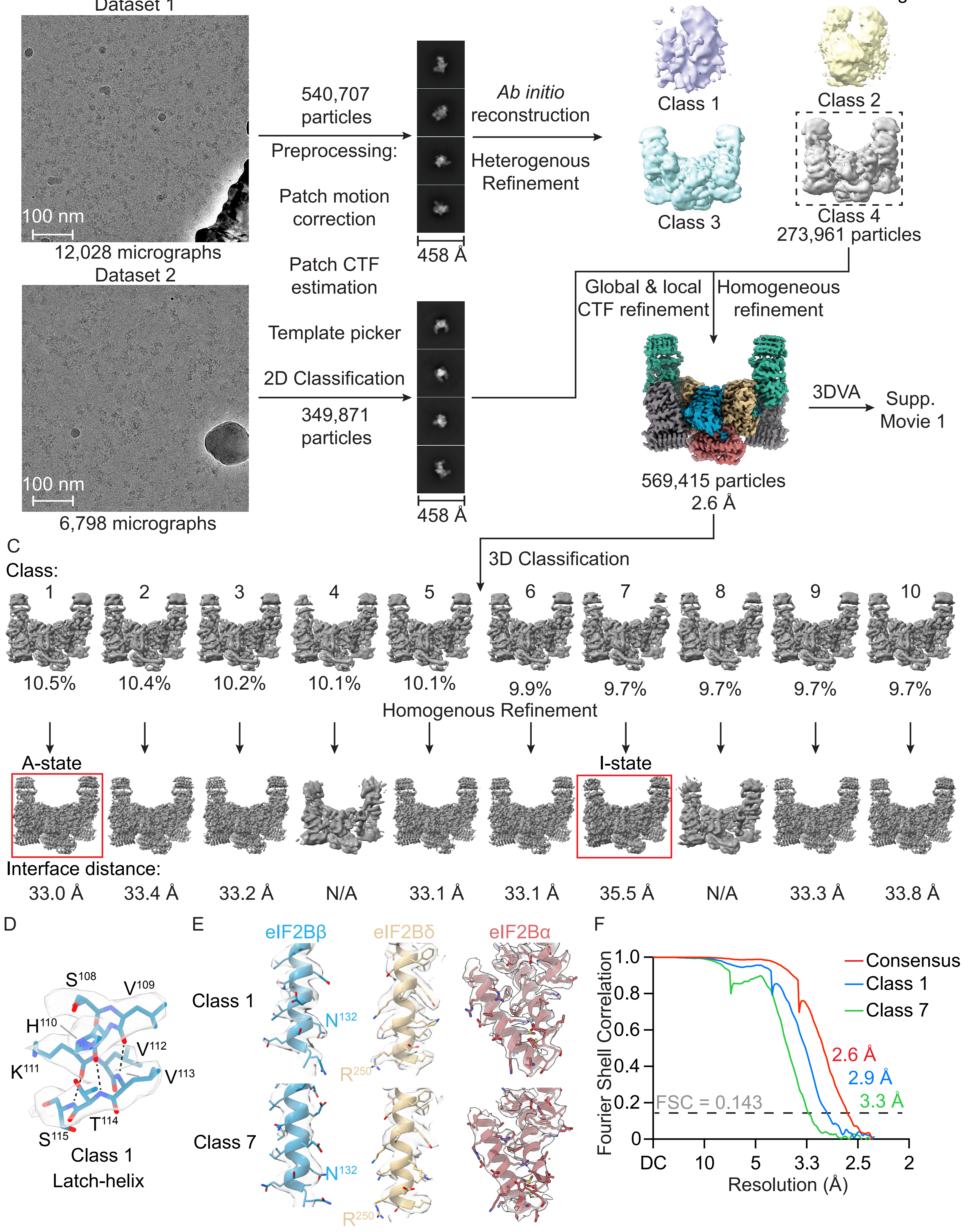
A
Dataset 1
100 nm
12,028 micrographs
Dataset 2
100 nm
6,798 micrographs
B
540,707
particles
Preprocessing:
Patch motion
correction
Patch CTF
estimation
Template picker
2D Classification
349,871
particles
458 Å
458 Å
Ab initio
reconstruction
Heterogenous
Refinement
Class 1
Class 2
Class 3
Class 4
273,961 particles
Global & local
CTF refinement
Homogeneous
refinement
3DVA
Supp.
Movie 1
569,415 particles
2.6 Å
C
3D Classification
Class:
1
2
3
4
5
6
7
8
9
10
10.5%
10.4%
10.2%
10.1%
10.1%
9.9%
9.7%
9.7%
9.7%
9.7%
Homogenous Refinement
A-state
I-state
Interface distance:
33.0 Å
33.4 Å
33.2 Å
N/A
33.1 Å
33.1 Å
35.5 Å
N/A
33.3 Å
33.8 Å
D
S108
V109
H110
V112
K111
V113
T114
S115
Class 1
Latch-helix
E
eIF2Bβ
eIF2Bδ
eIF2Bα
Class 1
N132
R250
Class 7
N132
R250
F
Consensus
Class 1
Class 7
Fourier Shell Correlation
1.0
0.8
0.6
0.4
0.2
0
2.6 Å
2.9 Å
3.3 Å
FSC = 0.143
DC
10
5
3.3
2.5
2
Resolution (Å)

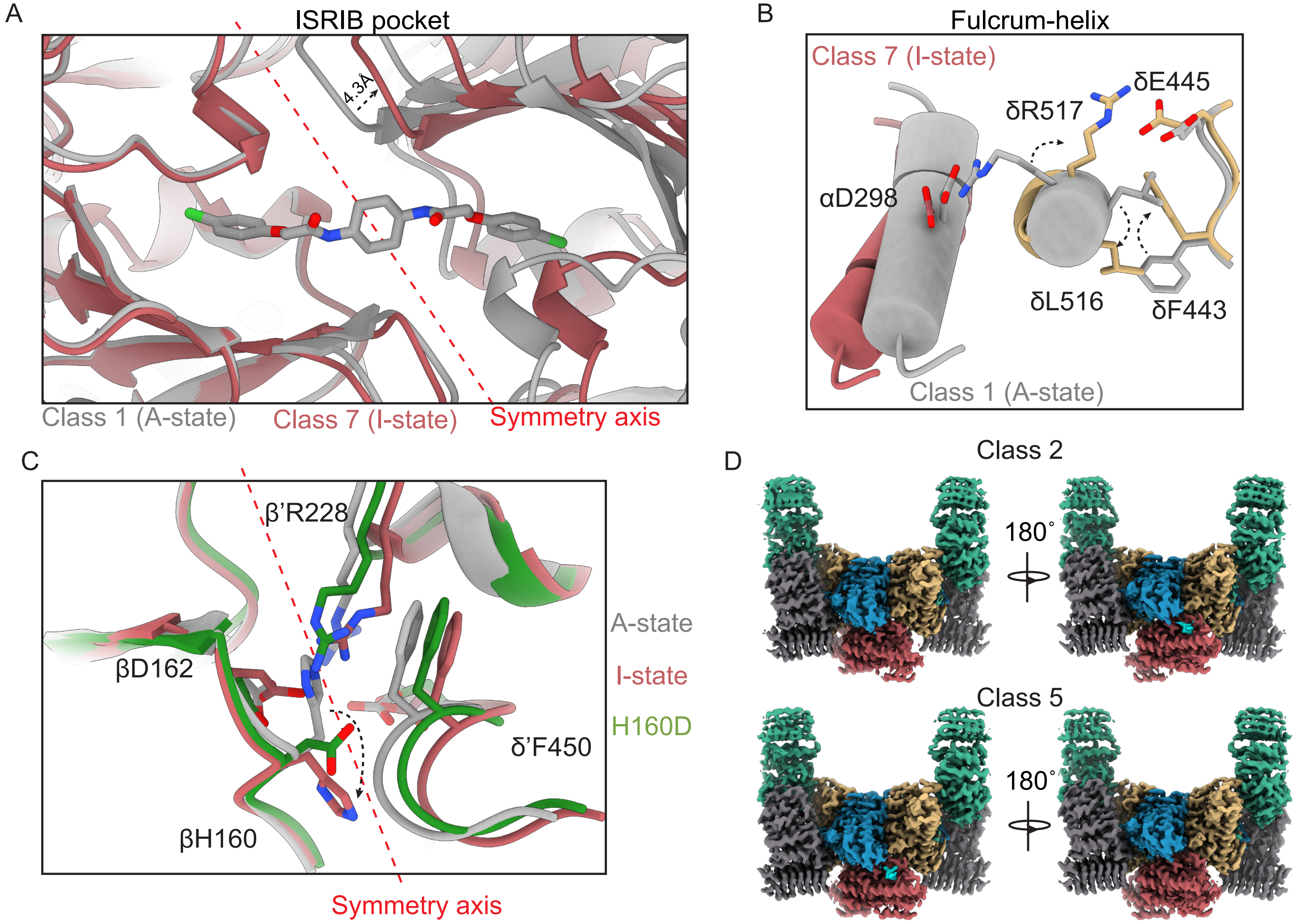
A
ISRIB pocket
4.3Å
Class 1 (A-state)
Class 7 (I-state)
Symmetry axis
B
Fulcrum-helix
Class 7 (I-state)
δE445
δR517
αD298
δL516
δF443
Class 1 (A-state)
C
β'R228
A-state
βD162
I-state
H160D
δ'F450
βH160
Symmetry axis
D
Class 2
180°
Class 5
180°

Figure S3

**>sp|P49770|EI2BB_HUMAN Translation initiation factor eIF2B subunit beta OS=Homo sapiens**

MPGSAAKGSELSERIESFVETLKRGGGPRSSEEMARETLGLLRQIITDHRWSNAGELMEL
IRREGRRMTAAQPSETTVGNMVRRVLKIIREEYGRLHGR**SDESDQQESLHKLLTSG**GLNE
DFSFHYAQLQSNIIEAINELLVELEGTMENIAAQALEHIHSNEVIMTIGFSRTVEAFLKE
AARKRKFHVIVAECAPFCQGHEMAVNLSKAGIETTVMTDAAIFAVMSRVNKVIIGTKTIL
ANGALRAVTGTHTLALAAKHHSTPLIVCAPMFKLSPQFPNEEDSFHKFVAPEEVLPFTEG
DILEKVSVHCPVFDYVPPELITLFISNIGGNAPSYIYRLMSELYHPDDHVL

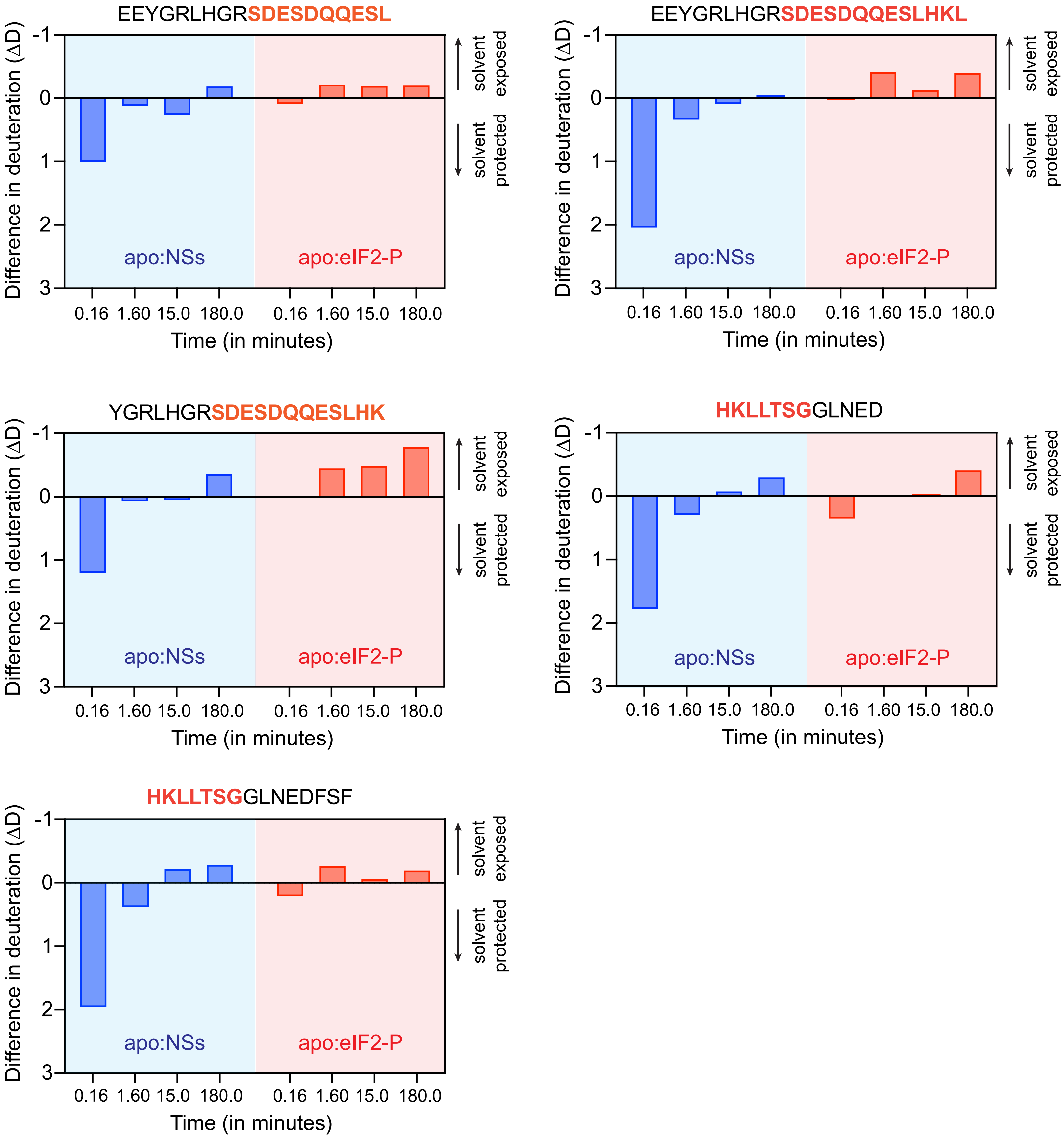

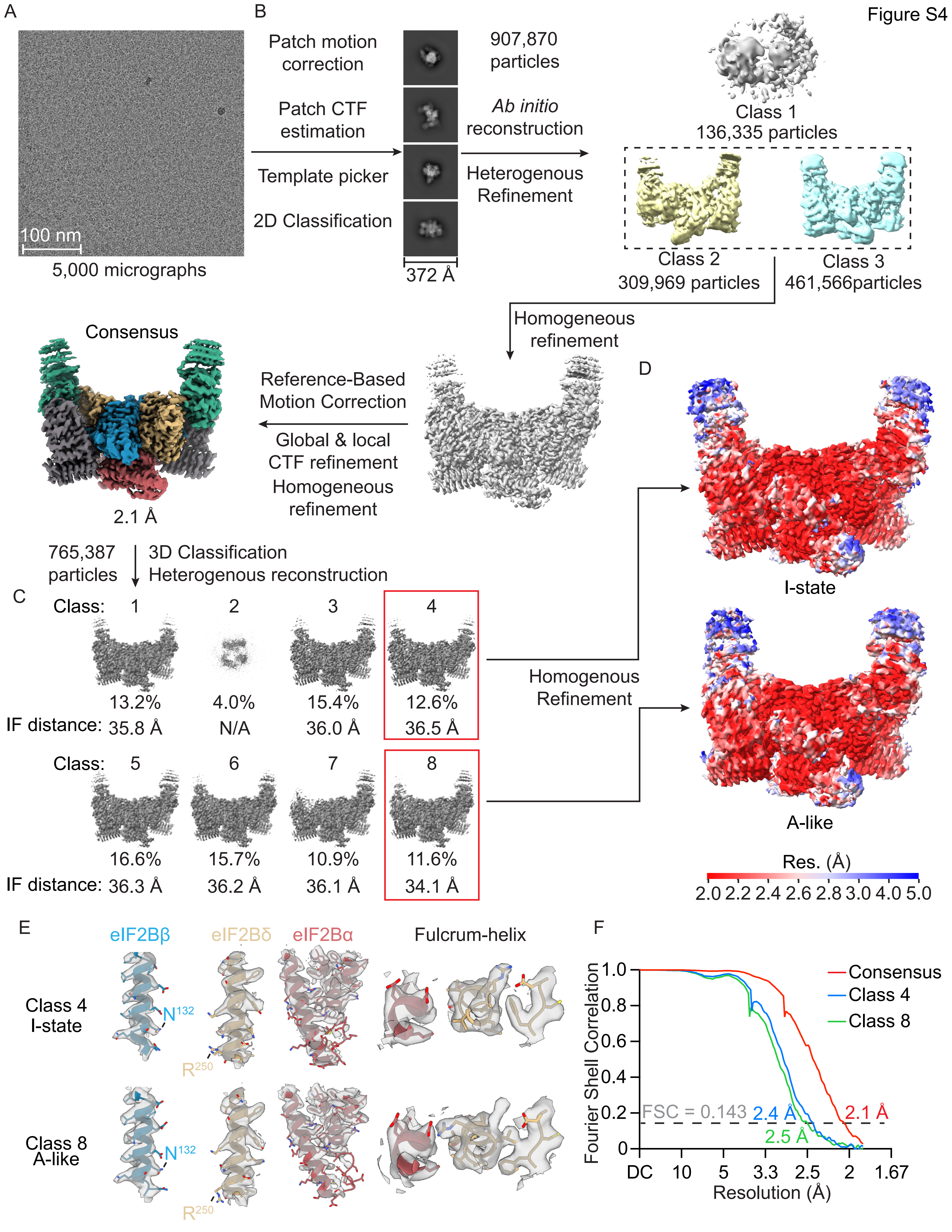
Figure S4
A
100 nm
5,000 micrographs
B
Patch motion correction
Patch CTF estimation
Template picker
2D Classification
372 Å
907,870 particles
Ab initio reconstruction
Heterogenous Refinement
Class 1
136,335 particles
Class 2
309,969 particles
Class 3
461,566particles
Homogeneous refinement
Reference-Based Motion Correction
Global & local CTF refinement
Homogeneous refinement
Consensus
2.1 Å
765,387 particles
3D Classification
Heterogenous reconstruction
C
Class: 1 2 3 4
13.2% 4.0% 15.4% 12.6%
IF distance: 35.8 Å N/A 36.0 Å 36.5 Å
Class: 5 6 7 8
16.6% 15.7% 10.9% 11.6%
IF distance: 36.3 Å 36.2 Å 36.1 Å 34.1 Å
Homogenous Refinement
D
I-state
A-like
Res. (Å)
2.0 2.2 2.4 2.6 2.8 3.0 4.0 5.0
E
eIF2Bβ
eIF2Bδ
eIF2Bα
Fulcrum-helix
Class 4 I-state
N132
R250
Class 8 A-like
N132
R250
F
Fourier Shell Correlation
1.0 0.8 0.6 0.4 0.2 0
Consensus
Class 4
Class 8
FSC = 0.143
2.4 Å
2.1 Å
2.5 Å
DC 10 5 3.3 2.5 2 1.67
Resolution (Å)

Figure S5

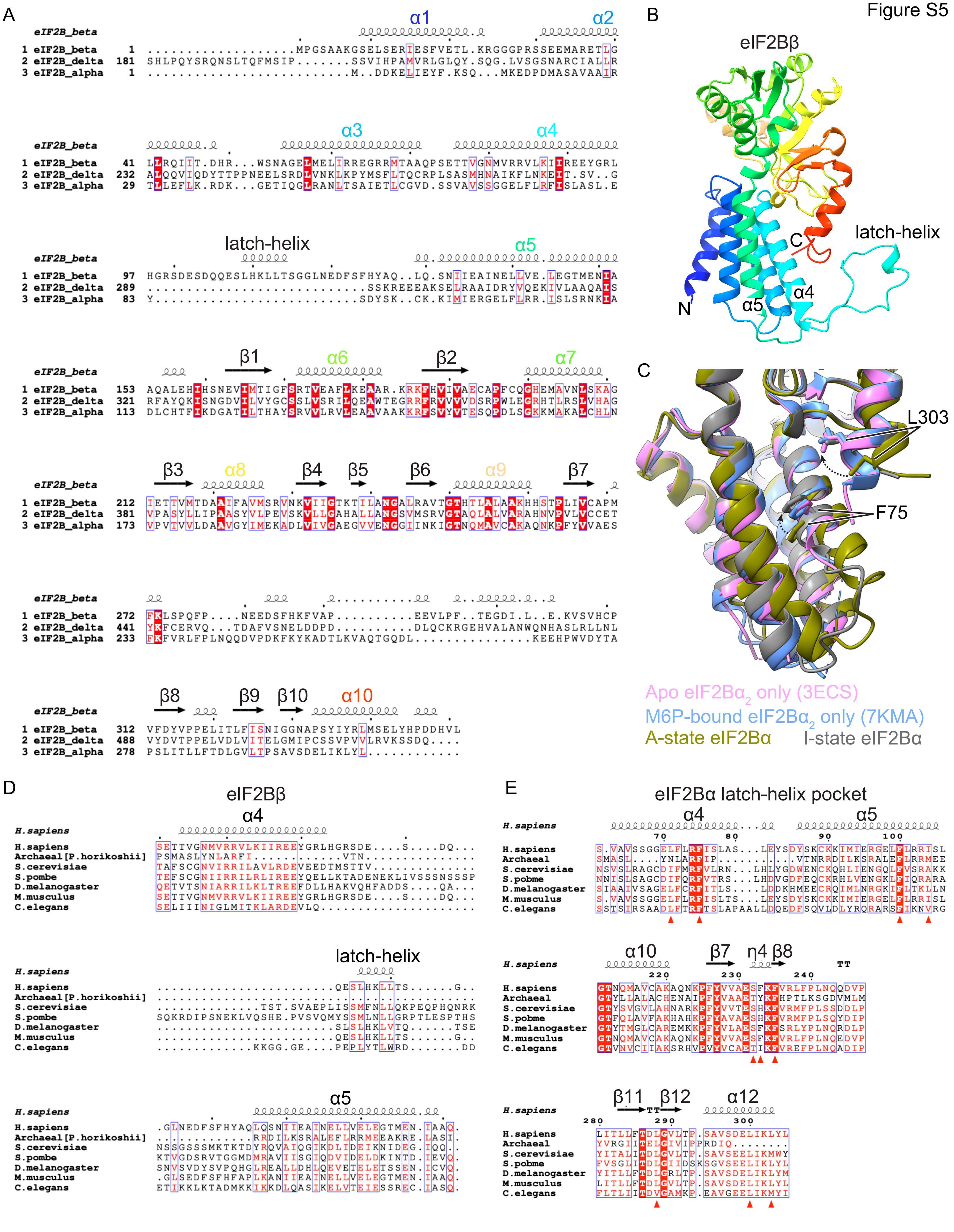
A
eIF2B_beta
1 eIF2B_beta
2 eIF2B_delta
3 eIF2B_alpha
α1
α2
α3
α4
latch-helix
α5
β1
α6
β2
α7
β3
α8
β4
β5
β6
α9
β7
β8
β9
β10
α10
B
eIF2Bβ
latch-helix
C
N
α5
α4
C
L303
F75
Apo eIF2Bα2 only (3ECS)
M6P-bound eIF2Bα2 only (7KMA)
A-state eIF2Bα
I-state eIF2Bα
D
eIF2Bβ
α4
latch-helix
α5
H.sapiens
Archaeal [P.horikoshii]
S.cerevisiae
S.pombe
D.melanogaster
M.musculus
C.elegans
E
eIF2Bα latch-helix pocket
α4
α5
α10
β7
η4
β8
β11
β12
α12
H.sapiens
Archaeal
S.cerevisiae
S.pombe
D.melanogaster
M.musculus
C.elegans

Figure S6

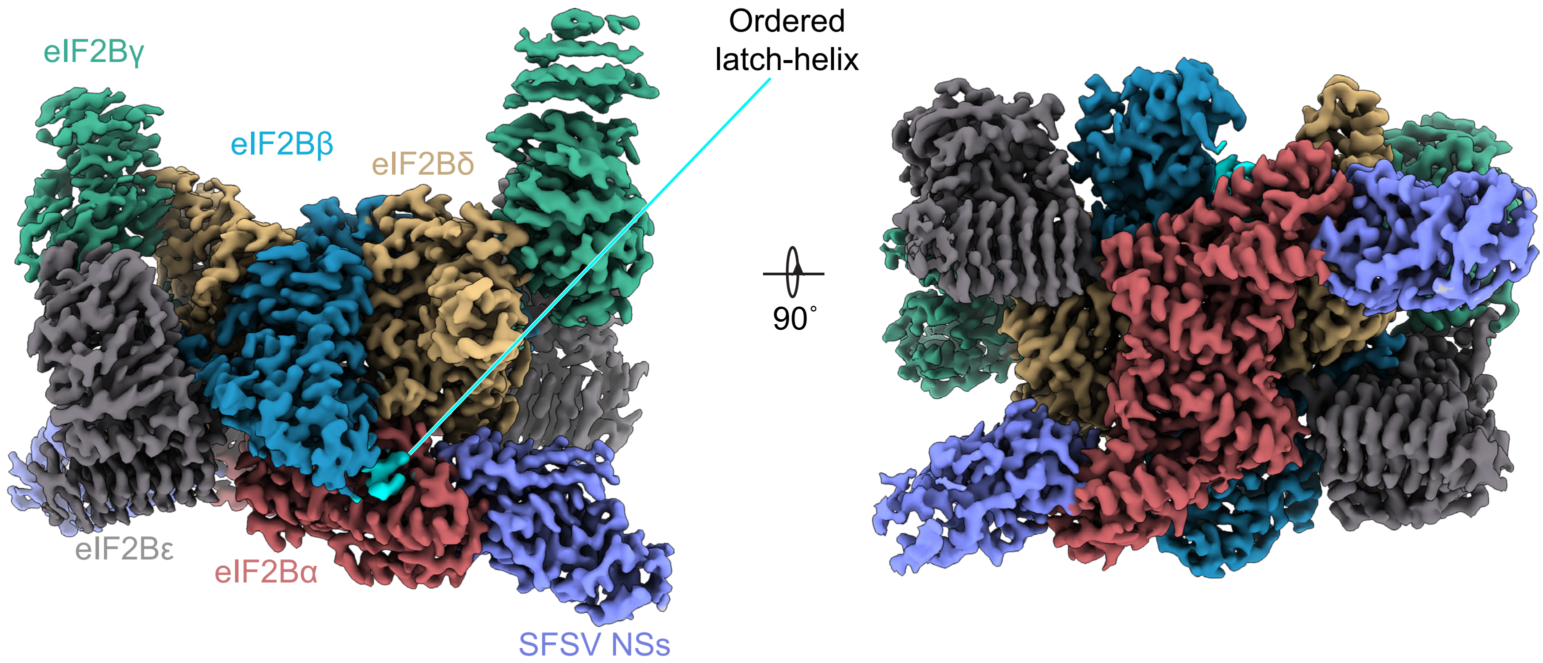

eIF2Bγ
eIF2Bβ
eIF2Bδ
Ordered
latch-helix
90˚
eIF2Bε
eIF2Bα
SFSV NSs

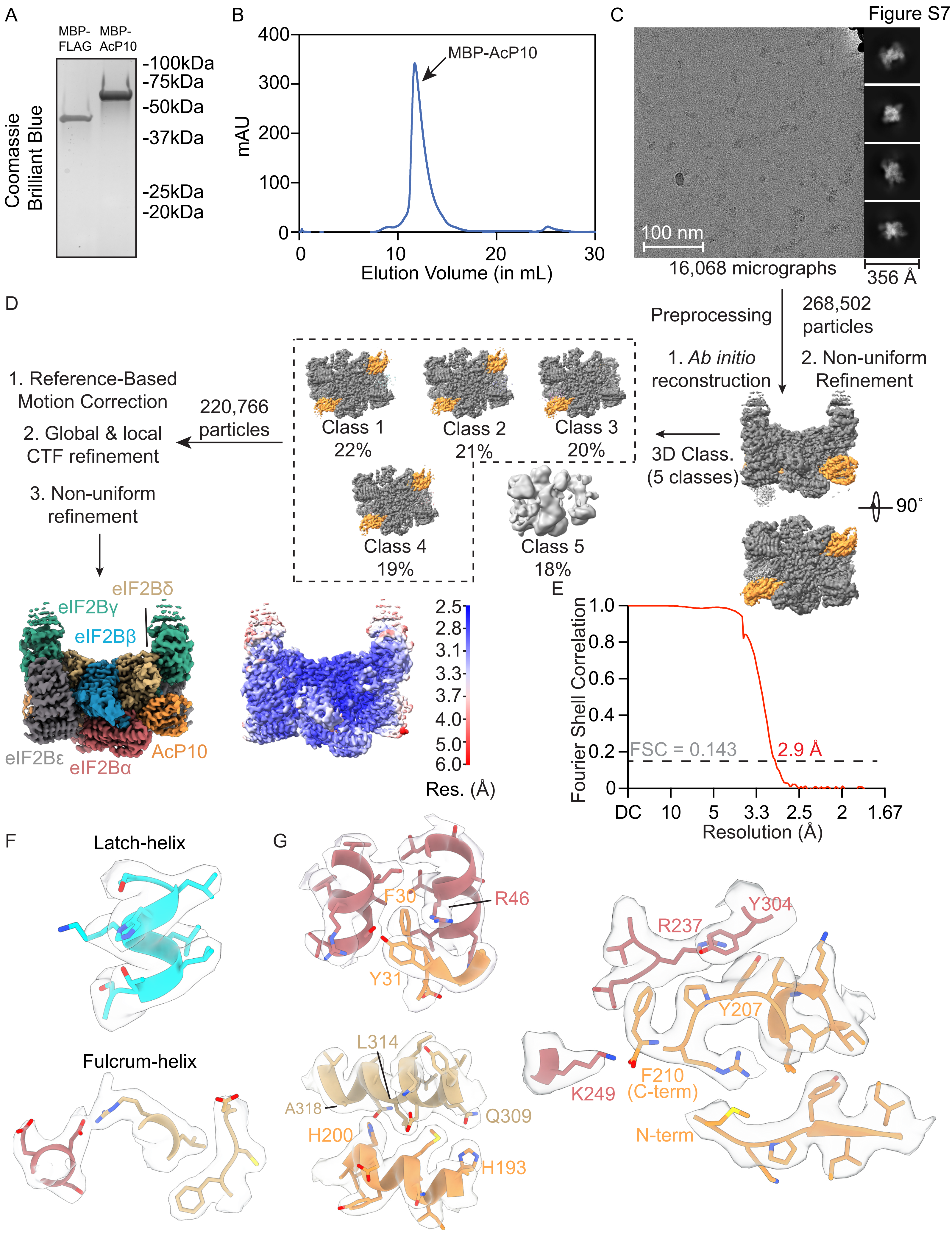

Figure S7
A
MBP-FLAG
MBP-AcP10
-100kDa
-75kDa
-50kDa
-37kDa
-25kDa
-20kDa
Coomassie Brilliant Blue
B
MBP-AcP10
mAU
Elution Volume (in mL)
C
100 nm
16,068 micrographs
356 Å
D
Preprocessing
268,502 particles
1. Ab initio reconstruction
2. Non-uniform Refinement
3D Class. (5 classes)
90˚
Class 1 22%
Class 2 21%
Class 3 20%
Class 4 19%
Class 5 18%
220,766 particles
1. Reference-Based Motion Correction
2. Global & local CTF refinement
3. Non-uniform refinement
eIF2Bδ
eIF2Bγ
eIF2Bβ
eIF2Bε
eIF2Bα
AcP10
Res. (Å)
E
Fourier Shell Correlation
FSC = 0.143
2.9 Å
Resolution (Å)
F
Latch-helix
Fulcrum-helix
G
F30
R46
Y31
L314
A318
Q309
H200
H193
R237
Y304
Y207
K249
F210 (C-term)
N-term

Figure S8

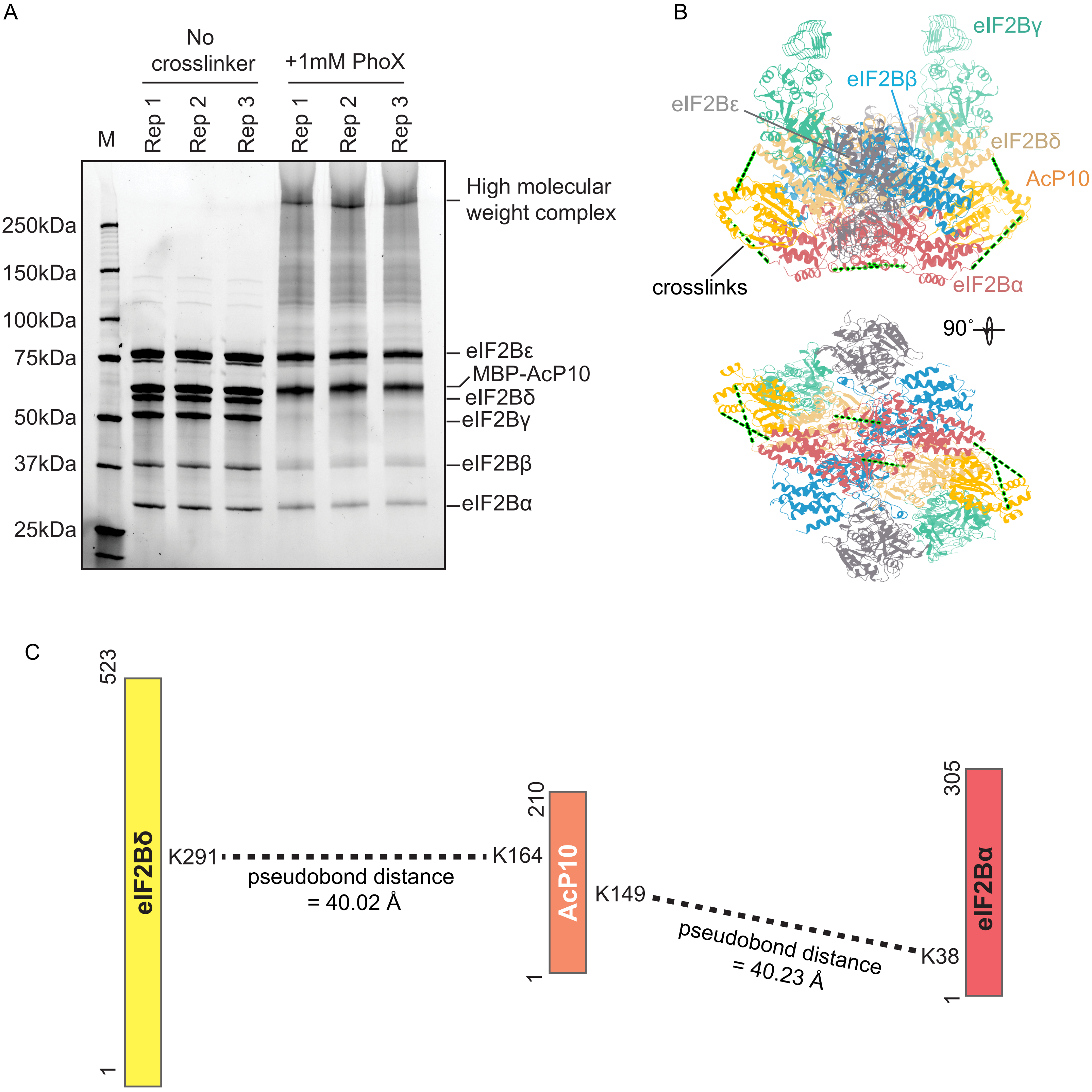

A
No crosslinker
+1mM PhoX
M
Rep 1
Rep 2
Rep 3
Rep 1
Rep 2
Rep 3
250kDa
150kDa
100kDa
75kDa
50kDa
37kDa
25kDa
High molecular weight complex
eIF2Bε
MBP-AcP10
eIF2Bδ
eIF2Bγ
eIF2Bβ
eIF2Bα
B
eIF2Bγ
eIF2Bβ
eIF2Bε
eIF2Bδ
AcP10
crosslinks
eIF2Bα
90˚
C
523
eIF2Bδ
1
K291
pseudobond distance
= 40.02 Å
K164
210
AcP10
1
K149
pseudobond distance
= 40.23 Å
K38
305
eIF2Bα
1

Figure S9

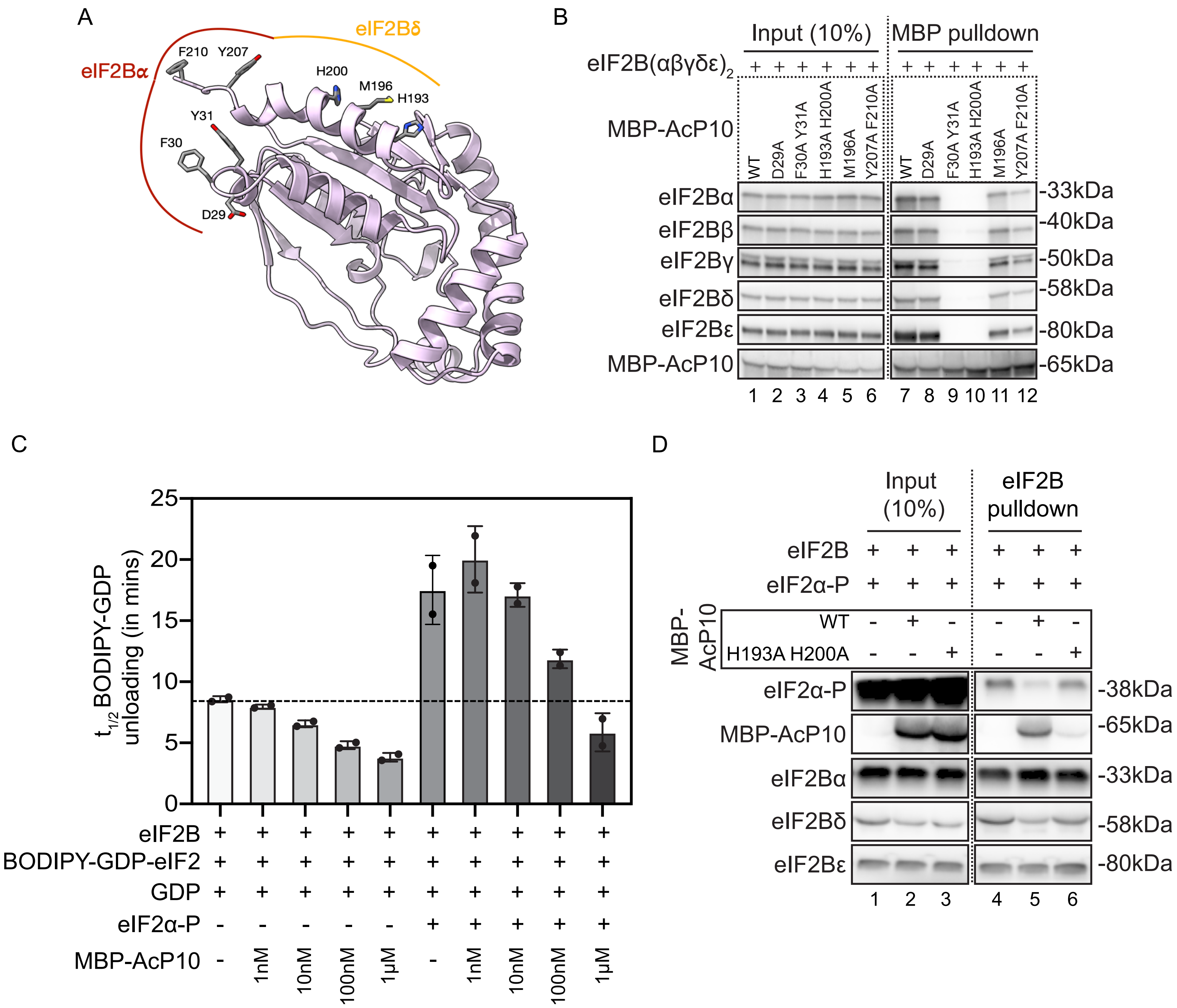
A
eIF2Bα
eIF2Bδ
F210
Y207
H200
M196
H193
Y31
F30
D29
B
Input (10%)
MBP pulldown
eIF2B(αβγδε)2
MBP-AcP10
WT
D29A
F30A Y31A
H193A H200A
M196A
Y207A F210A
eIF2Bα
eIF2Bβ
eIF2Bγ
eIF2Bδ
eIF2Bε
MBP-AcP10
-33kDa
-40kDa
-50kDa
-58kDa
-80kDa
-65kDa
1 2 3 4 5 6 7 8 9 10 11 12
C
t1/2 BODIPY-GDP unloading (in mins)
0
5
10
15
20
25
eIF2B
BODIPY-GDP-eIF2
GDP
eIF2α-P
MBP-AcP10
1nM
10nM
100nM
1μM
D
Input (10%)
eIF2B pulldown
eIF2B
eIF2α-P
MBP-AcP10
WT
H193A H200A
eIF2α-P
MBP-AcP10
eIF2Bα
eIF2Bδ
eIF2Bε
-38kDa
-65kDa
-33kDa
-58kDa
-80kDa
1 2 3 4 5 6

Figure S10

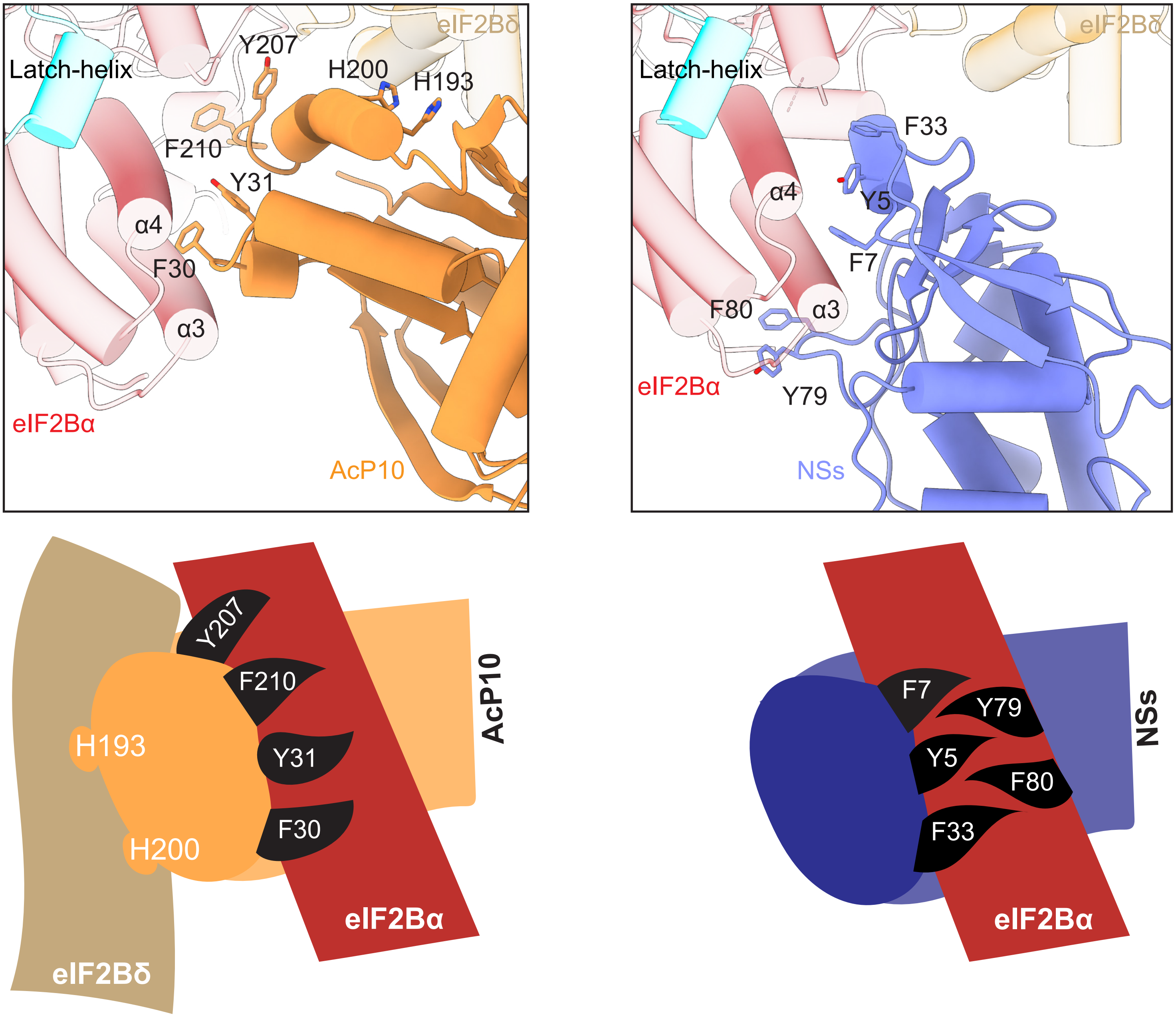

Latch-helix
Y207
H200
H193
eIF2Bδ
F210
Y31
α4
F30
α3
eIF2Bα
AcP10
Latch-helix
eIF2Bδ
F33
α4
Y5
F7
F80
α3
eIF2Bα
Y79
NSs
Y207
F210
H193
Y31
F30
H200
AcP10
eIF2Bα
eIF2Bδ
F7
Y79
Y5
F80
F33
NSs
eIF2Bα

Figure S11

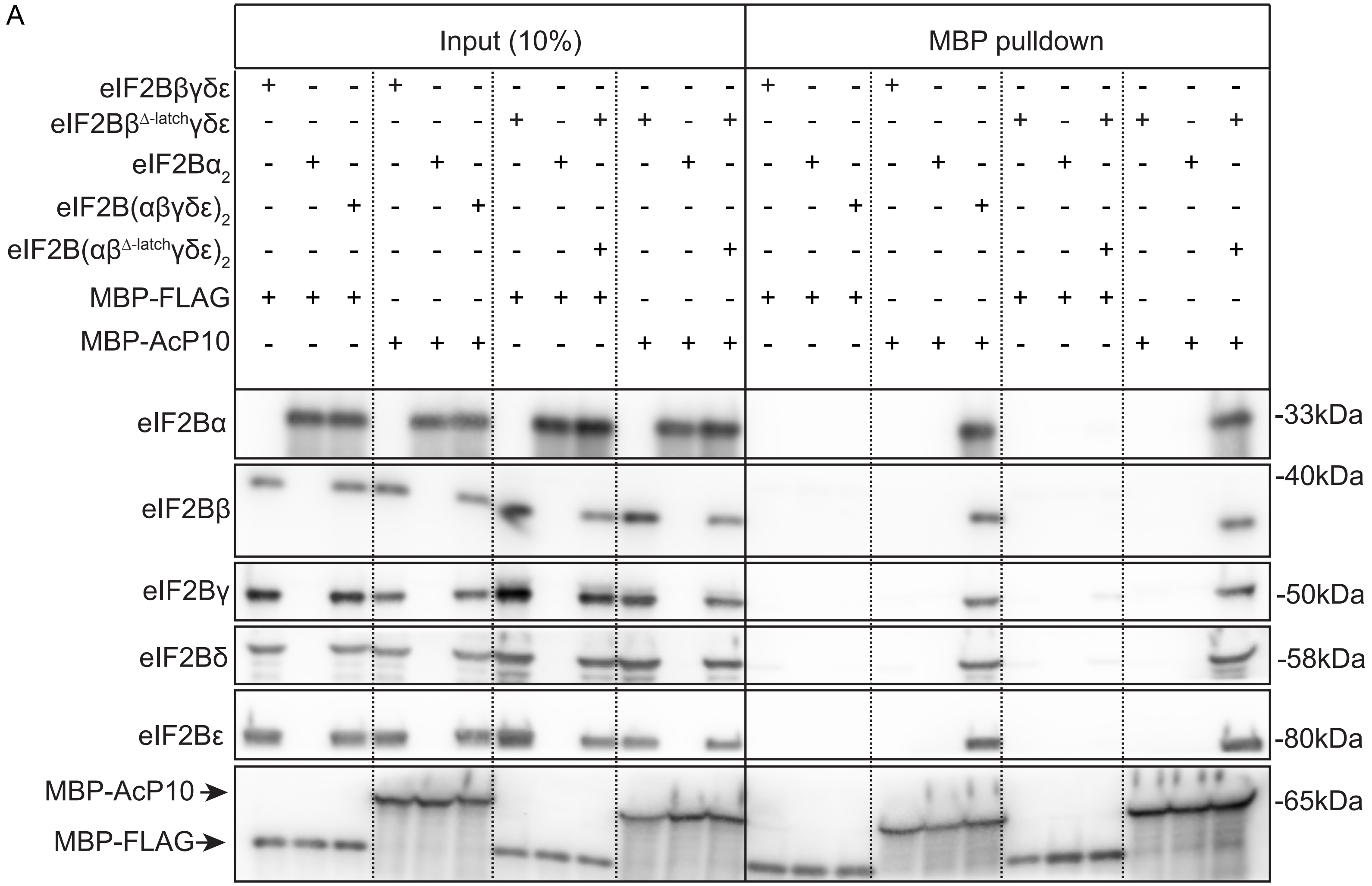
A
Input (10%)
MBP pulldown
eIF2Bβγδε
eIF2Bβ^Δ-latch γδε
eIF2Bα2
eIF2B(αβγδε)2
eIF2B(αβ^Δ-latch γδε)2
MBP-FLAG
MBP-AcP10
eIF2Bα
eIF2Bβ
eIF2Bγ
eIF2Bδ
eIF2Bε
MBP-AcP10
MBP-FLAG
-33kDa
-40kDa
-50kDa
-58kDa
-80kDa
-65kDa

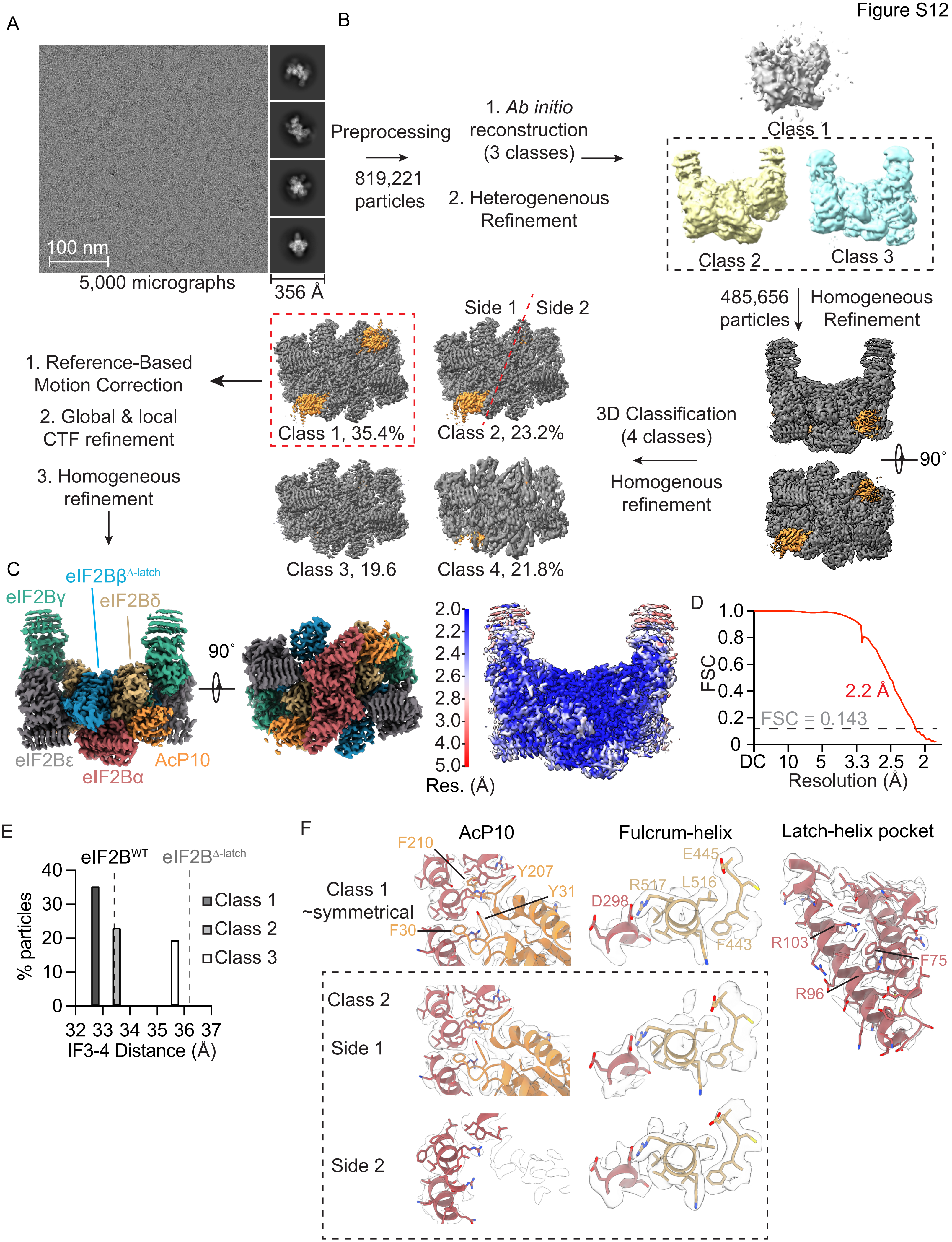
A
100 nm
5,000 micrographs
356 Å
B
Preprocessing
819,221 particles
1. Ab initio reconstruction (3 classes)
2. Heterogenenous Refinement
Class 1
Class 2
Class 3
485,656 particles
Homogeneous Refinement
90°
3D Classification (4 classes)
Homogenous refinement
Side 1
Side 2
Class 1, 35.4%
Class 2, 23.2%
Class 3, 19.6
Class 4, 21.8%
1. Reference-Based Motion Correction
2. Global & local CTF refinement
3. Homogeneous refinement
C
eIF2Bβ$^{\Delta\text{-latch}}$
eIF2Bγ
eIF2Bδ
eIF2Bε
eIF2Bα
AcP10
90°
Res. (Å)
D
FSC
2.2 Å
FSC = 0.143
Resolution (Å)
DC 10 5 3.3 2.5 2
E
eIF2B$^{WT}$
eIF2B$^{\Delta\text{-latch}}$
% particles
Class 1
Class 2
Class 3
IF3-4 Distance (Å)
F
AcP10
Fulcrum-helix
Latch-helix pocket
Class 1 ~symmetrical
F210
Y207
Y31
F30
E445
R517
L516
D298
F443
R103
F75
R96
Class 2
Side 1
Side 2

Figure S13

A

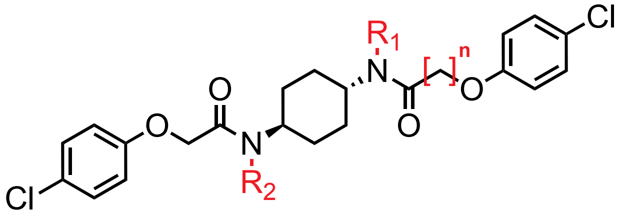


B

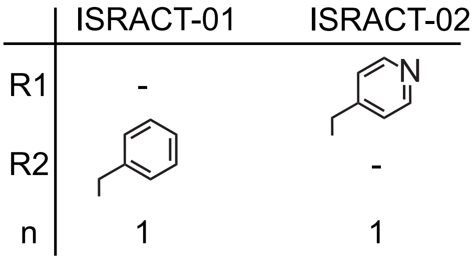


C

ISRACT-01:

3-(4-chlorophenoxy)-N-[trans-4-[N-benzyl-2-(4-chlorophenoxy)acetamido]cyclohexyl]propanamide

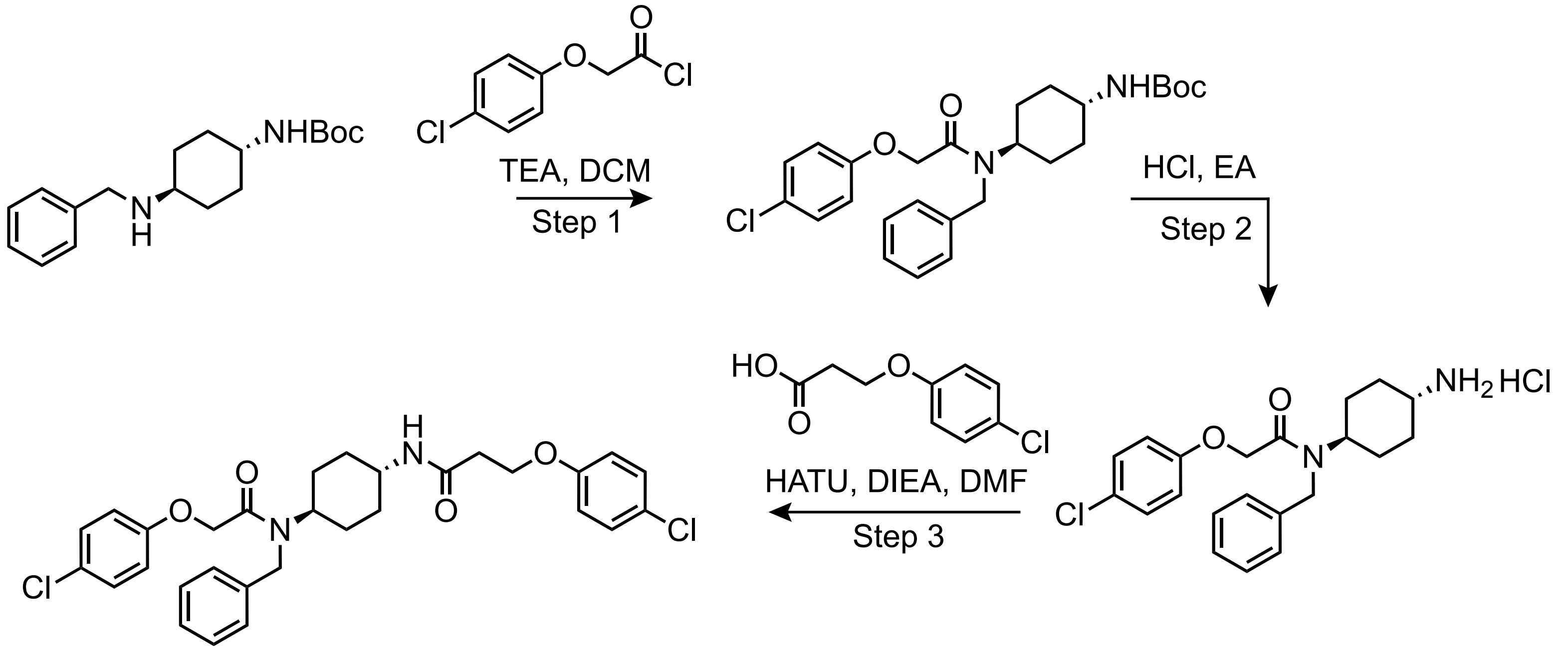


ISRACT-01

D

ISRACT-02:

3-(4-chlorophenoxy)-N-(pyridin-4-ylmethyl)-N-[trans-4-[2-(4-chlorophenoxy)acetamido]cyclohexyl]propanamide

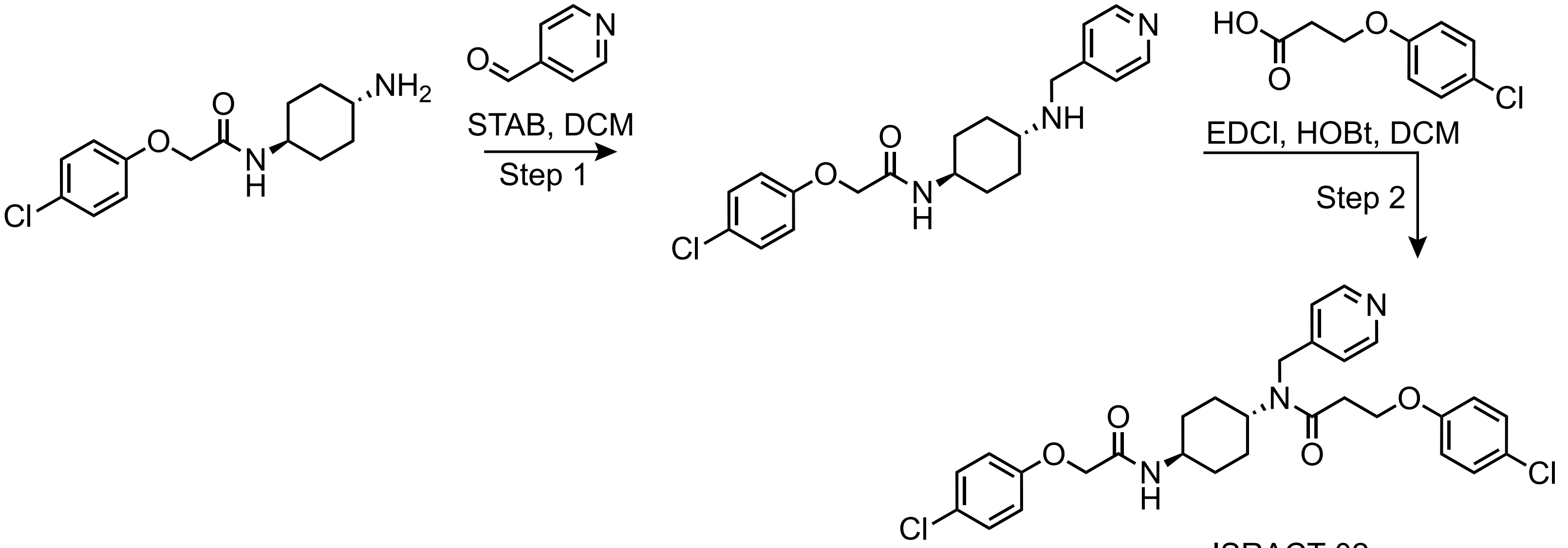


ISRACT-02

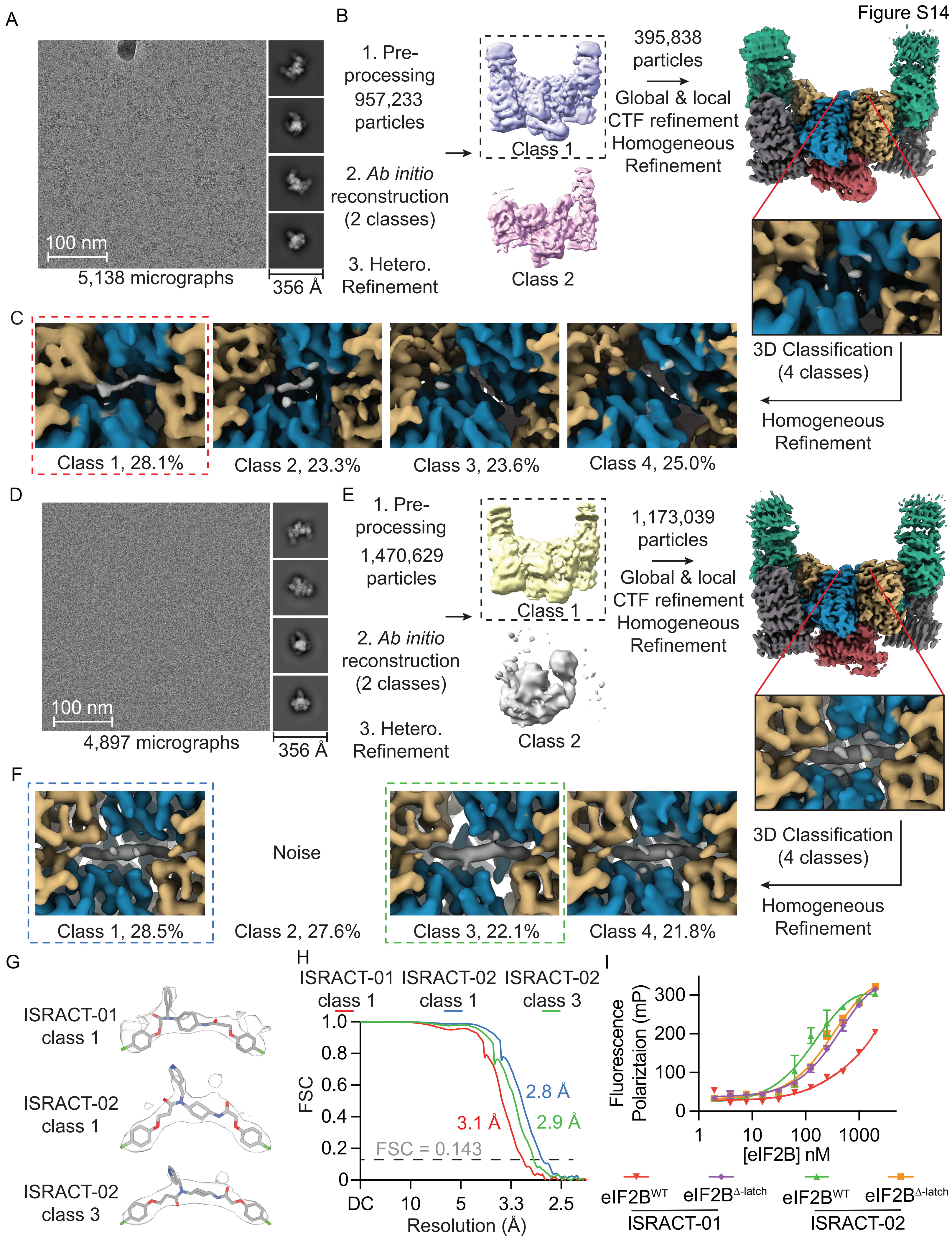

Figure S14
A
100 nm
5,138 micrographs
356 Å
B
1. Pre-processing 957,233 particles
2. Ab initio reconstruction (2 classes)
3. Hetero. Refinement
Class 1
Class 2
395,838 particles
Global & local CTF refinement
Homogeneous Refinement
3D Classification (4 classes)
Homogeneous Refinement
C
Class 1, 28.1%
Class 2, 23.3%
Class 3, 23.6%
Class 4, 25.0%
D
100 nm
4,897 micrographs
356 Å
E
1. Pre-processing 1,470,629 particles
2. Ab initio reconstruction (2 classes)
3. Hetero. Refinement
Class 1
Class 2
1,173,039 particles
Global & local CTF refinement
Homogeneous Refinement
3D Classification (4 classes)
Homogeneous Refinement
F
Class 1, 28.5%
Noise
Class 2, 27.6%
Class 3, 22.1%
Class 4, 21.8%
G
ISRACT-01 class 1
ISRACT-02 class 1
ISRACT-02 class 3
H
ISRACT-01 class 1
ISRACT-02 class 1
ISRACT-02 class 3
FSC
Resolution (Å)
FSC = 0.143
3.1 Å
2.8 Å
2.9 Å
1.0
0.8
0.6
0.4
0.2
0
DC
10
5
3.3
2.5
I
Fluorescence Polariztaion (mP)
[eIF2B] nM
300
200
100
0
1
10
100
1000
eIF2B^WT
eIF2B^Δ-latch
ISRACT-01
eIF2B^WT
eIF2B^Δ-latch
ISRACT-02

Figure S15

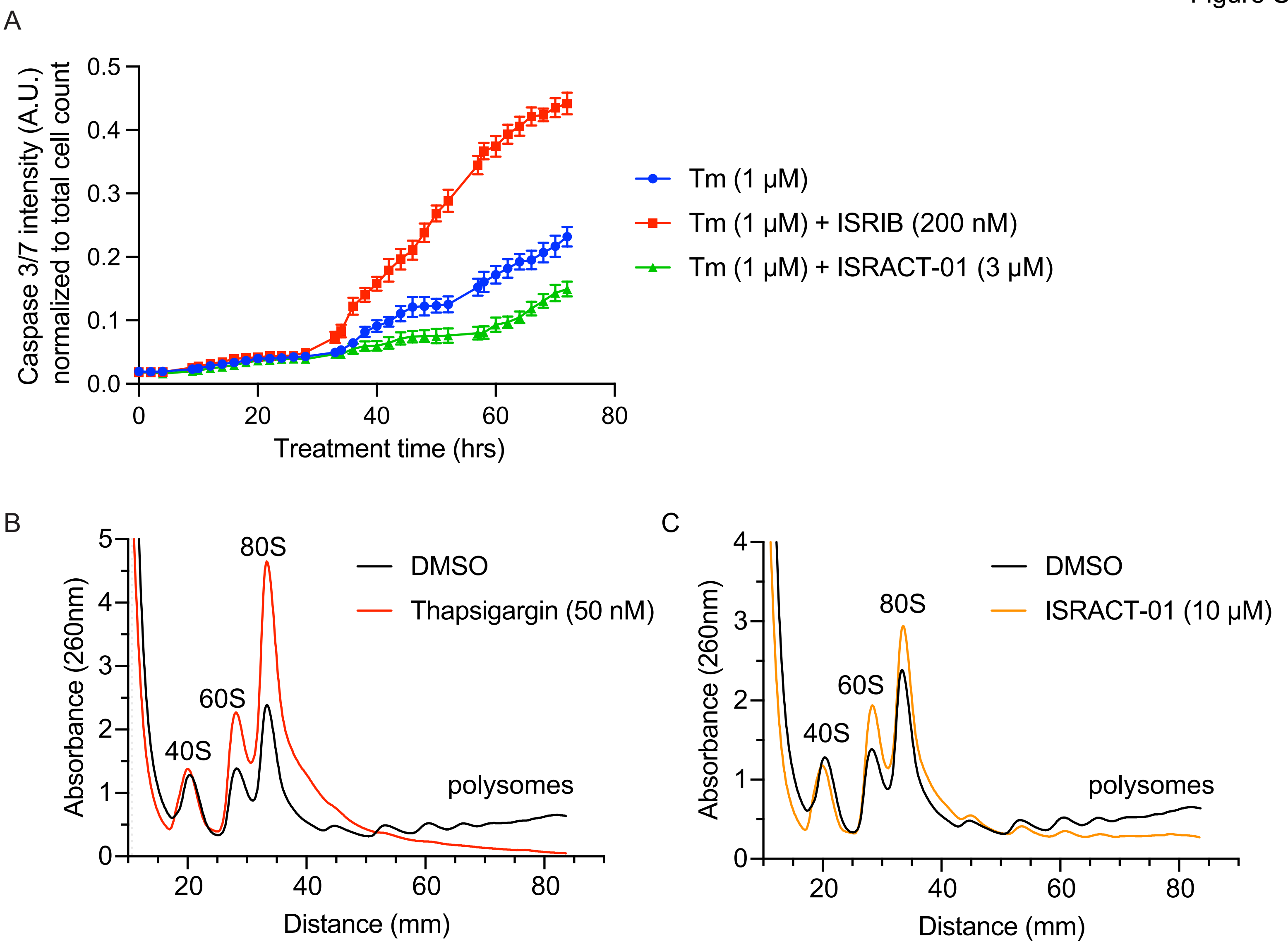
A
Caspase 3/7 intensity (A.U.) normalized to total cell count
0.0
0.1
0.2
0.3
0.4
0.5
0
20
40
60
80
Treatment time (hrs)
Tm (1 μM)
Tm (1 μM) + ISRIB (200 nM)
Tm (1 μM) + ISRACT-01 (3 μM)
B
Absorbance (260nm)
0
1
2
3
4
5
40S
60S
80S
polysomes
20
40
60
80
Distance (mm)
DMSO
Thapsigargin (50 nM)
C
Absorbance (260nm)
0
1
2
3
4
40S
60S
80S
polysomes
20
40
60
80
Distance (mm)
DMSO
ISRACT-01 (10 μM)
